# Enhancer-driven cell type comparison reveals similarities between the mammalian and bird pallium

**DOI:** 10.1101/2024.04.17.589795

**Authors:** Nikolai Hecker, Niklas Kempynck, David Mauduit, Darina Abaffyová, Roel Vandepoel, Sam Dieltiens, Ioannis Sarropoulos, Carmen Bravo González-Blas, Elke Leysen, Rani Moors, Gert Hulselmans, Lynette Lim, Joris De Wit, Valerie Christiaens, Suresh Poovathingal, Stein Aerts

## Abstract

Combinations of transcription factors govern the identity of cell types, which is reflected by enhancer codes in cis-regulatory genomic regions. Cell type-specific enhancer codes at nucleotide-level resolution have not yet been characterized for the mammalian neocortex. It is currently unknown whether these codes are conserved in other vertebrate brains, and whether they are informative to resolve homology relationships for species that lack a neocortex such as birds. To compare enhancer codes of cell types from the mammalian neocortex with those from the bird pallium, we generated single-cell multiome and spatially-resolved transcriptomics data of the chicken telencephalon. We then trained deep learning models to characterize cell type-specific enhancer codes for the human, mouse, and chicken telencephalon. We devised three metrics that exploit enhancer codes to compare cell types between species. Based on these metrics, non-neuronal and GABAergic cell types show a high degree of regulatory similarity across vertebrates. Proposed homologies between mammalian neocortical and avian pallial excitatory neurons are still debated. Our enhancer code based comparison shows that excitatory neurons of the mammalian neocortex and the avian pallium exhibit a higher degree of divergence than other cell types. In contrast to existing evolutionary models, the mammalian deep layer excitatory neurons are most similar to mesopallial neurons; and mammalian upper layer neurons to hyper- and nidopallial neurons based on their enhancer codes. In addition to characterizing the enhancer codes in the mammalian and avian telencephalon, and revealing unexpected correspondences between cell types of the mammalian neocortex and the chicken pallium, we present generally applicable deep learning approaches to characterize and compare cell types across species via the genomic regulatory code.

## Introduction

Genomic enhancers form the core part of gene regulatory networks (GRNs) that maintain the identity of cell types. GRNs comprise combinations of transcription factors (TFs) that bind to specific transcription factor binding sites (TFBS) in enhancer regions to regulate the expression of target genes. These combinations of TFBS form enhancer codes that are characteristic for the identity of cell types (*1–3*). Several tools have been developed to leverage single-cell RNA-seq (scRNA-seq) and chromatin accessibility (scATAC-seq) data to identify cell type specific GRNs (*4, 5*). However, characteristic combinations of TFBS for a cell type are difficult to detect using these methods. Sequence-based deep learning models have shown major advances to delineate which sequence patterns in enhancer regions are important for cell type specific chromatin accessibility or gene expression (*6, 7*). They have contributed substantially to identify TFBS specific to mammalian interneurons (*8, 9*), fly brain cell types (*10*), mouse liver cells (*11*), and mouse embryonic stem cells (*12*). Furthermore, deep learning models have been applied to predict chromatin accessibility across mammalian brain cell types (*13, 14*), to compare enhancer codes of melanocytes across species (*15*), and to identify potential enhancer regions linked to the evolution of neocortex expansion and vocal learning (*8, 16*). As these deep learning models allow us to identify enhancer codes in cell type-specific enhancer regions, we hypothesized that they may shed light on cell type conservation across species.

Vertebrate telencephala pose ideal examples to study the conservation of enhancer codes as they comprise a variety of cell types that are expected to be maintained by either similar or diversified GRNs across species (*17*). Despite shared developmental trajectories, telencephala of different vertebrate brains display a strikingly different neuroanatomy (*18*). As an iconic example, the six-layered neocortex is found in mammals but is absent in other vertebrates (*17*). Different homologies between structures of the vertebrate telencephalon have been suggested based on their developmental origin and circuitry (*19–21*). Single cell sequencing has been used to compare the transcriptome of cells from the mammalian neocortex, reptilian three-layered cortex, and three telencephalic nuclei in song birds (*22–24*). Both non-neuronal and neuronal cortical cell types have overall conserved molecular identities between human, marmoset, and mouse despite transcriptomic differences (*24*). GABAergic neurons were found to be conserved between the reptilian cortex and mammalian neocortex based on scRNA-seq data, whereas cross-species relationships of excitatory neurons could not be clearly assigned based on transcriptome comparisons (*22*). However, groups of turtle excitatory neurons resemble either upper layer or deep layer neurons of the mammalian neocortex (*22*). For songbirds, two nuclei related to vocal learning (HVC and robust nucleus of the arcopallium) have been suggested to exhibit similarities to distinct mammalian neocortical neurons but do not have the same developmental origin as their potential mammalian counterparts (*23*). Thus, how the majority of the avian telencephalic cell types relate to those of other vertebrates has yet to be deciphered. While transcriptome comparisons suggest some similarities between telencephalic cell types across vertebrates, they do not take into account genomic signatures in enhancer regions that may provide additional insights into the conservation of cell types.

## Results

### Transcriptome similarities between avian and mammalian telencephalic cell types

To compare cell types and dissect enhancer codes in the mammalian telencephalon, we analyzed two mouse brain and two human cortex single-cell datasets (Fig. 1) (*4, 24–26*). Cell types in human and mouse cerebral cortex have 1-1 homology relationships at a subclass level which we recapitulate based on transcriptome comparisons with SAMap, a method that optimizes transcriptome correspondences and gene orthology relationships (*27*) (Methods, fig. S1A and B). These 1-1 correspondences agree with previous findings by Bakken et al. (*24*). To compare mammalian cell types with those from another amniote lineage, we generated chicken telencephalon single-cell multiome and spatially-resolved transcriptomics (Stereo-seq) datasets containing 23,179 cells and 27,487 spatial bins (*28*).

**Fig. 1.**
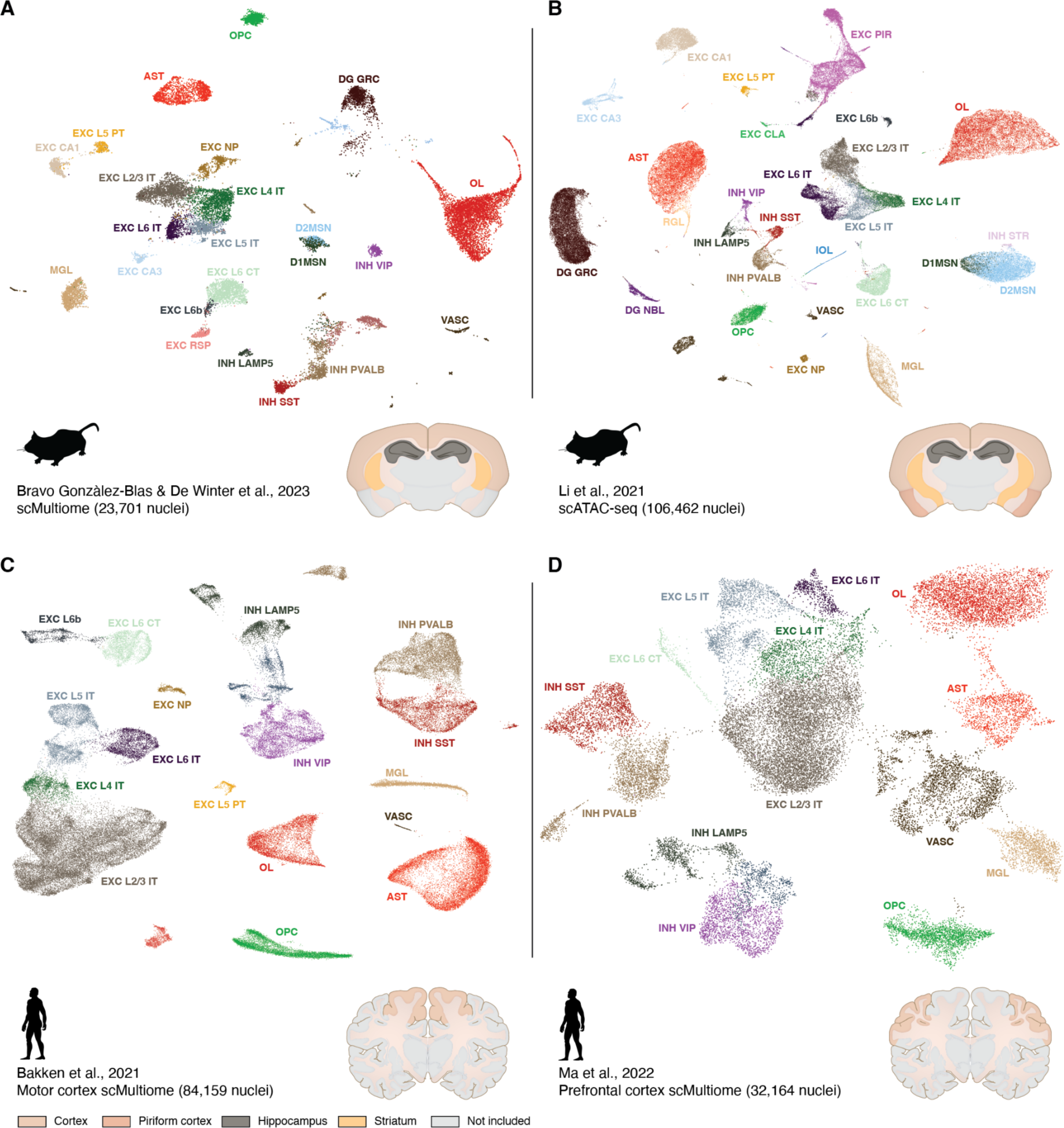
Conserved cell types in mouse and human single cell data sets. UMAPs show clusters of cells and corresponding cell types for mouse telencephalon (**A, B**) and human cortex single cell data sets (**C, D**). Pictograms indicate the approximate brain regions from which the data was sampled. AST, Astrocytes; D1/2MSN, D1/2 medium spiny neurons; DG GRC, Dentate gyrus granule cells; DG NBL, Dentate gyrus granule cell neuroblasts; EXC CA1/3, Cornu Ammonis 1/3 hippocampal excitatory neurons; EXC CLA, Claustrum excitatory neurons; EXC L2/3/4/5/6 IT, Layer 2/3/4/5/6 intra-telencephalic excitatory neurons; EXC L5 PT, Layer 5 pyramidal tract excitatory neurons; EXC L6 CT, Layer 6 corticothalamic excitatory neurons; EXC L6b, Layer 6b excitatory neurons; EXC NP, Layer 5 near-projecting excitatory neurons; EXC PIR, Piriform cortex excitatory neurons; EXC RSP, Retrosplenial excitatory neurons; INH LAMP5, LAMP5 lateral ganglionic eminence derived interneurons; INH PVALB, PVALB medial ganglionic eminence derived interneurons; INH SST, SST medial ganglionic eminence derived interneurons; INH STR, Striatum derived interneurons; INH VIP, VIP lateral ganglionic eminence derived interneurons; IOL, Intermediate oligodendrocytes; MGL, Microglia; OL, Oligodendrocytes; OPC, Oligodendrocyte precursors; RGL, Radial-glial like cells; VASC, Vascular endothelial cells.

Clusters of chicken telencephalon cells map to distinct locations in the chicken telencephalon including the striatum, nidopallium or hyperpallium, mesopallium, entopallium, and hippocampal region based on spatially-resolved transcriptomics data (Fig. 2A). Besides non-neuronal cell types, we identified seven clusters of GABAergic and eight clusters of glutamatergic neurons using known marker genes for mammalian brain cell types (Fig. 2B). We detected clusters corresponding to the major interneuron subclasses PVALB+, SST+ and LAMP5+ by their cognate markers supporting previous findings, and VIP-like interneurons by known marker genes though not by expression of VIP itself (*23*). Similarly, cell types of the striatum, D1/D2 medium spiny neurons (D1/2MSN) and other striatal-like GABAergic neurons express mammalian marker genes. Glutamatergic neurons could not easily be assigned to mammalian cell types by known marker genes, although we detected one cluster of RORB+ cells in the entopallium, in-line with previous findings (*29*). To identify potential additional homologies between cell types and to corroborate our marker-based assignments, we employed SAMap to match chicken to mouse cell types (*27*) (Methods). SAMap suggests an unambiguous 1-1 correspondence between mammalian and bird GABAergic neuron subclasses, as well as between all non-neuronal cell types (Fig. 2C). The identification of correspondences of most excitatory neurons are more ambiguous although several chicken telencephalon clusters exhibit preferences for different mouse neurons subclasses, including EXC GLU-1 (nido- and hyperpallium) to neocortex layer 2/3 intratelencephalic (L2/3 IT) neurons, EXC GLU-2 (nido- and hyperpallium) and EXC GLU-3 (entopallium) to L4 IT neurons, EXC GLU-4/5/7 (mesopallium) to deeper layer neurons, and EXC GLU-6 (medial pallium) and EXC GLU-8 to hippocampal or dentate gyrus cell types. In addition to SAMap derived cell type correspondences, we confirmed the transcriptome similarities by comparing the correlation of expression levels of 1:1 orthologous genes across avian and mammalian cell types (fig. S2) (*22*). In conclusion, most chicken cell types which transcriptomes exhibit correspondences to the different mouse brain cell types are localized in distinct nuclei or regions of the chicken telencephalon.

**Fig. 2.**
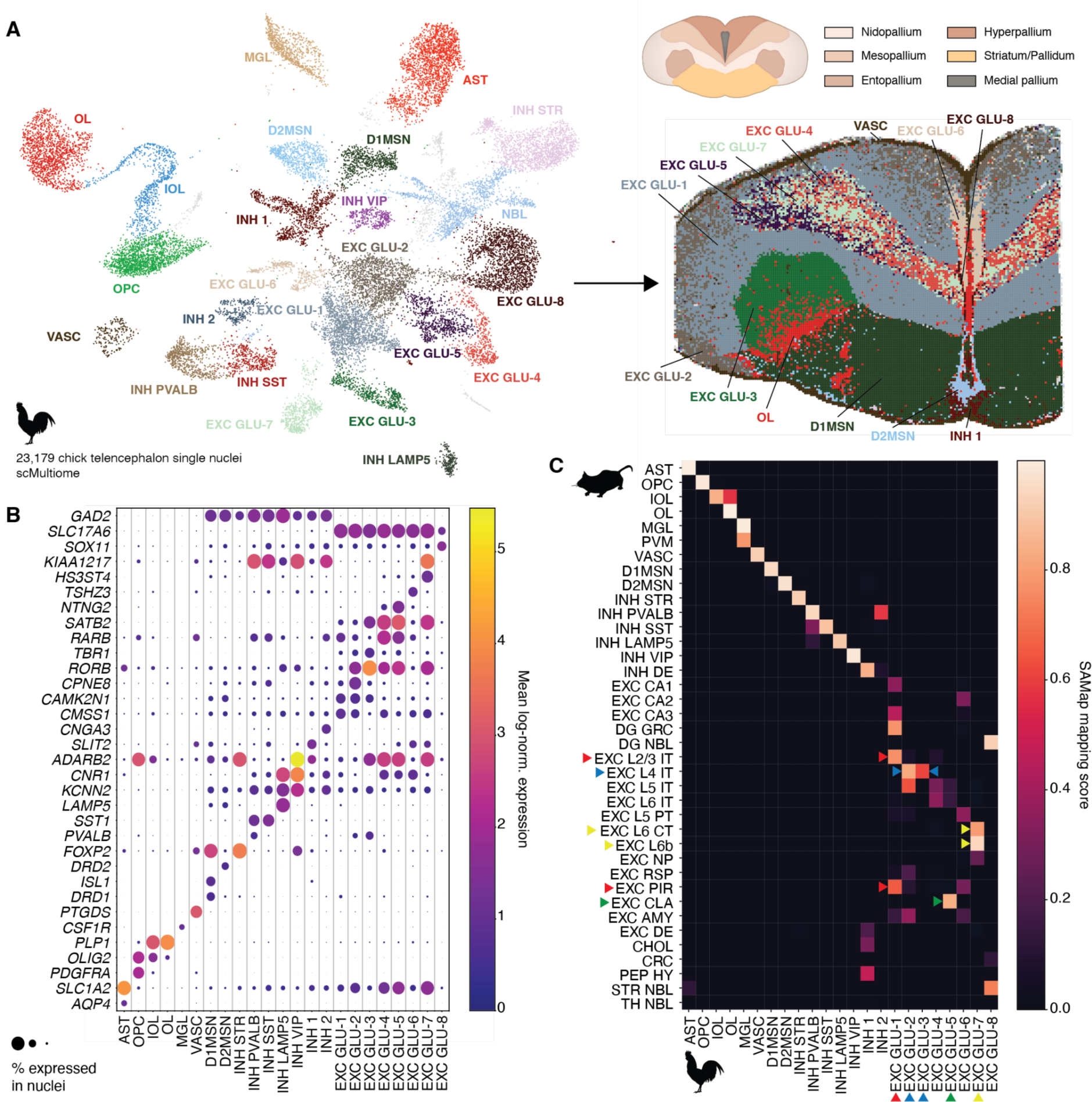
Cell types of the chicken telencephalon and expression of marker genes. (**A**) The UMAP shows clusters of single cells and inferred cell types or assigned cluster labels (left). Single cell clusters were mapped to spatially-resolved transcriptomics data (Stereo-seq). Spatial spots are assigned to single cell clusters based on the highest density of mapped clusters labels (right). The pictogram shows anatomical subdivisions of the chicken telencephalon. (**B**) The expression of characteristic marker genes is shown per chicken telencephalon cell type. Dot sizes indicate the fraction of nuclei of a cluster that express the gene. (**C**) Transcriptomes of chicken telencephalon cell types were compared to mouse brain cell type transcriptomes using a mouse brain single nuclei gene expression data set (*30*). Triangles highlight potential cell type homologs for excitatory neurons.

### Enhancer codes of non-neuronal cell types are highly conserved between birds and mammals

After finding similarities between avian and mammalian cell types based on transcriptomes, the question arises whether we can find genomic signatures in the form of genomic enhancer codes in accessible regions that reflect cell type homologies. Our rationale is that while enhancer regions are not necessarily sequence-conserved across more distantly related species, cell-type specific gene regulation through TFs is often conserved (*31–33*). In agreement with this assumption, the correlation of 1-1 ortholog TF expression is sufficient to delineate which cell types are similar between mouse and chicken (fig. S2) (*22*). If TF mediated gene regulation is conserved during evolution, we expect selective pressures that preserve specific combinations of TFBS that form enhancer codes in cis-regulatory regions.

To test this assumption, we used genomic regions with differential chromatin accessibility between telencephalic cell types as a proxy for potential enhancer regions and investigated them with deep learning models. In particular, we trained deep learning models to predict cell types directly from DNA sequences of differentially accessible regions (DARs) for the different telencephalic cell types (Fig. 3A). We trained separate models for the two mouse brain (DeepMouseBrain1&2), two human cortex (DeepHumanCortex1&2), and the chicken telencephalon (DeepChickenBrain) datasets, respectively (fig. S3). The use of independently trained models for two mouse and two human datasets allows us to infer robust cross-species predictions. To verify that our models are capable of identifying homologous cell types based on their enhancer codes, we first evaluated whether the deep learning models can recapitulate the cell type homologies between birds and mammals that we would expect based on transcriptome comparisons. We evaluated their prediction performance on cell type-specific DARs of non-neuronal cell types and neurons grouped into three broad categories: medium spiny neurons (MSN), interneurons, and excitatory neurons. For each chicken cell type in our dataset, we used the top 100 DARs with the highest log-fold change per cell type and predicted mammalian cell types with the DeepMouseBrain and DeepHumanCortex models (Fig. 3B). Scoring mouse or human regions in the same manner gives identical results (fig. S4, A and B). Similarly to the SAMap comparison between chicken and mouse (Fig. 2C), we find near 1:1 correspondences between avian and mammalian non-neuronal cell types directly from accessible DNA sequences. Excitatory neurons, MSN and interneurons grouped together are also classified correctly. The same regions scored by the human models validate these matches. These findings confirm the transcriptomic cell type matches and suggest that our models show a robust generalization in classifying previously unseen sequences from different species.

**Fig. 3.**
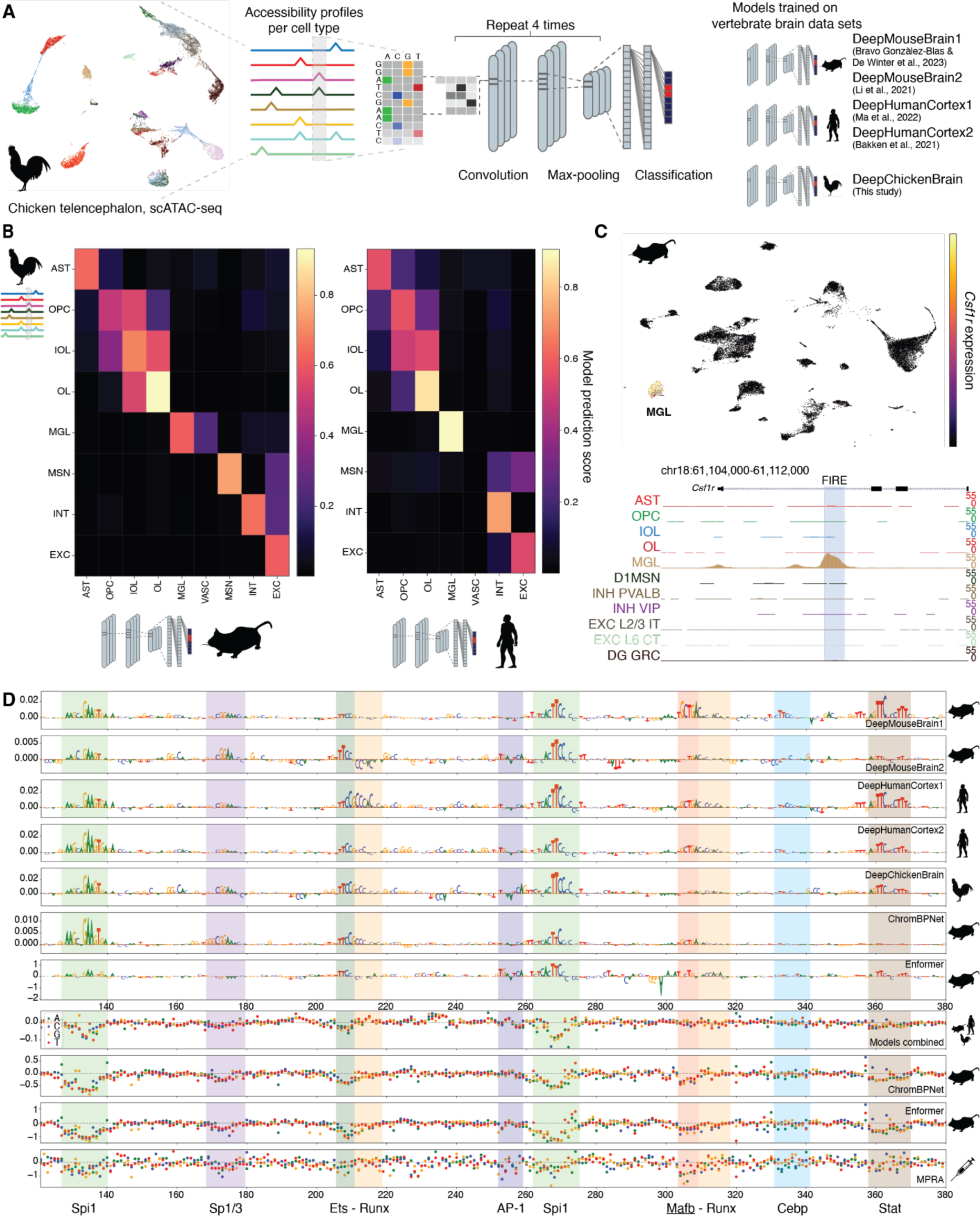
Cross species enhancer modeling. (**A**) Overview of enhancer model architecture and workflow. For each of the five brain datasets, a model was trained. (**B**) Cross-species model predictions from the mouse (left) and human (right) models on the top 100 DARs per chicken cell type. The median prediction score per region is shown. Both mouse and human predictions are the consensus of the two respective models per species. Excitatory (EXC), medium spiny neuron (MSN) and interneuron (INT) cell types are grouped into broad cell type categories. (**C**) scRNA-seq UMAP of mouse brain cells from the DeepMouseBrain1 dataset (*4*) showing the microglia-specific expression of the *Csf1r* gene (top). The cell-type specific chromatin accessibility of the microglia enhancer FIRE (mm10 chr18:61108475-61108975), regulating *Csf1r* expression, in microglia is depicted underneath. (**D**) Nucleotide contribution scores of FIRE obtained from the five enhancer models (order: DeepMouseBrain1, DeepMouseBrain2, DeepHumanCortex1, DeepHumanCortex2, DeepChickenBrain), from a microglia ChromBPNet model and from Enformer are shown. ISM scores are shown for a consensus of all five enhancer models, the ChromBPNet model and Enformer. Finally, IVM based MPRA activity in mouse BV2 cells is shown. Previously identified TF binding sites are highlighted, as well as an uncharacterized potential Mafb binding site.

Secondly, we asked whether the deep learning models that were trained independently on data from different species learned the same sequence features, that is, enhancer codes that correspond to TFBS. To identify sequence features, we used both gradient based (SHAP) contribution scores (*34, 35*) and in-silico-mutagenesis (ISM) (*36, 37*) to derive nucleotide contribution scores at each position of the sequence of a DAR. These nucleotide contribution scores describe the importance of each nucleotide per sequence for predicting a specific cell type and allow us to identify important regulatory subsequences, or motifs, corresponding to TFBS that are characteristic for cell type identity. As an illustrative example, we investigated the enhancer code of the mouse “fms intronic regulatory element” (FIRE) that regulates the expression of the *Csf1r* gene in microglia (*38*). This region has indeed microglia-specific chromatin accessibility in our dataset and *Csf1r* is specifically expressed in microglia (Fig. 3C). The enhancer’s nucleotide importance scores are highly correlated (average pairwise Spearman correlation = 0.45, average P-value = 1.78 x 10^-11^) for all of our models and all species, and detect previously validated TFBS as the most important features of the sequence (Fig. 3D). Nucleotide importance scores derived from other models with different architectures and training methods agree with the ones obtained from our models (*39, 40*) (Fig. 3D, Methods). To further validate the model predictions, we performed in vitro mutagenesis via massive parallel reporter assays (MPRA) in the BV2 mouse cell line. The in vitro enhancer activity profile of this microglia cell line shows decreases in activity specifically for nucleotide changes that distort predicted TFBS. This confirms the nucleotide importance scores, as well as previously experimentally identified TFBS in this enhancer (Fig. 3D) (*38*). Additionally, we find a previously unidentified Mafb-like binding site in all mammalian models that is confirmed by the MPRA activity. These results indicate that our models independently learned the same features for explaining their predictions for the mouse microglia enhancer. This suggests conservation of the microglia enhancer code across the three amniote species, and provides confidence in our trained model to enable cross-species comparisons and to detect important cis-regulatory features.

Given that our models trained on data from different species learned similar enhancer codes for different cell types, we can utilize them for assessing homologies between cell types. We illustrate this for a candidate enhancer in mouse astrocytes (Fig. 4A). This region is located inside an intron of *Prdm16* and is conserved (sequence identity 60%) and specifically accessible in both chicken and mouse astrocytes (log-fold change 5.33 and 5.32) (Fig. 4A). During development, *Prdm16* is expressed in radial glia and contributes to cell migration through transcriptional silencing (*41*). In adult mammalian and avian brains, *Prdm16* is a characteristic astrocyte marker gene (Fig. 4B). The DeepMouseBrain, DeepChickenBrain, and DeepHumanCortex models accurately predict this region as being specific to astrocytes (Fig. 4A & fig. S5A). To compare the similarity of learned astrocyte enhancer codes between our chicken, human, and mouse models, we computed nucleotide contribution scores for all cell types and compared them with contribution scores derived from the other models (Methods). This allowed us to assess for which cell types two models learned the most similar enhancer codes. In particular, we compared nucleotide contribution scores that our DeepMouseBrain1 model learned for explaining astrocytes with nucleotide contribution scores for all cell types derived from our DeepChicken and DeepHuman brain models (Methods). Nucleotide contribution scores for the astrocyte class of the DeepChicken model show a high degree of similarity to nucleotide contributions scores for the astrocyte class of the DeepMouseBrain1 model suggesting similar enhancer code specific to astrocytes (Spearman correlation coefficient of 0.63, p-value = 9.81 x 10^-55^). As we observed for the microglia enhancer (Fig. 4D), the sequence features with high importance scores correspond to potential TFBS. The DeepMouseBrain2 and the DeepHumanCortex models identify these TFBS as well (fig. S5B). In addition, the sequence alignment between the mouse and chicken regions shows that point mutations and insertions occur at nucleotides that the models do not deem important, while the identified TFBS remain conserved. The same applies for the importance scores of the chicken homologous region (fig. S5C). As a negative control, contribution scores for the chicken intermediate oligodendrocyte (IOL) cell type do not indicate any important TFBS in this region and are anti-correlated with the mouse astrocyte contribution scores (Spearman correlation of -0.25, p-value= 1.66 x 10^-8^) (Fig. 4C). Furthermore, the presence of Rora/b binding sites exhibits negative contribution scores towards predicting this sequence as an IOL region. Hence, the correlation between nucleotide contribution scores can be used to compare for which cell types two different models learned the most similar enhancer codes, which reflects cell type homologies.

**Fig. 4.**
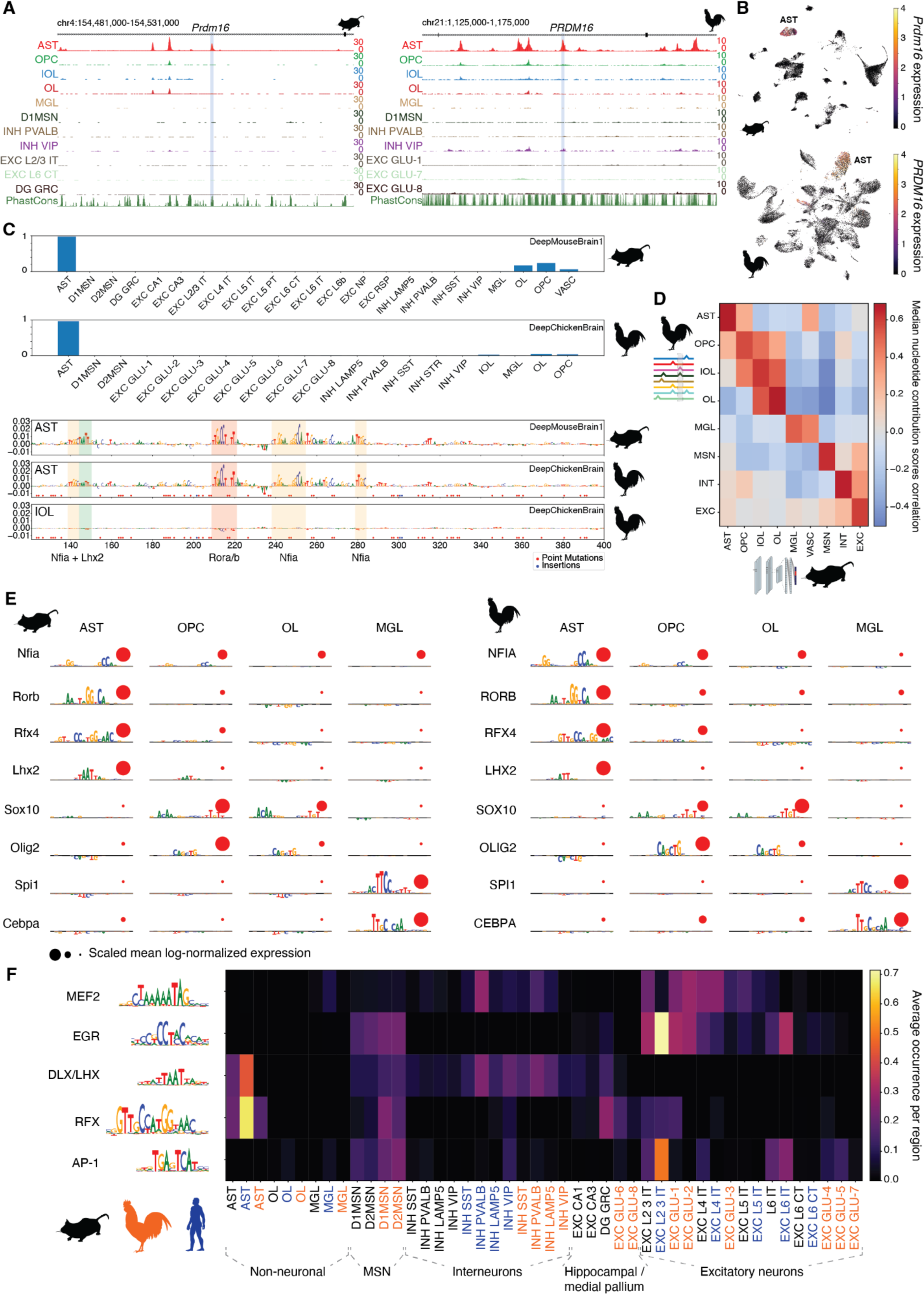
Conservation of enhancer code in vertebrate brain cell types. (**A**) scATAC tracks of the *PRDM16* region in chicken (galGal6 chr21:1148446-1148946) and mouse (mm10 chr4:154506677-154507177) showing astrocyte specificity. (**B**) scRNA-seq UMAPs of *Prdm16* expression in mouse (DeepMouseBrain1 dataset) and chicken. (**C**) DeepMouseBrain1 and DeepChickenBrain predictions on the mouse region, accompanied by DeepMouseBrain1 contribution scores for the astrocyte class and DeepChickenBrain contribution scores for the astrocyte and intermediate oligodendrocyte classes. Point mutations and insertions in the mouse sequence when aligned to the chicken sequence are indicated in red and blue respectively. TFs that potentially bind to nucleotide features with high contribution scores are indicated and assigned based on their similarity to known TFBS motifs and cell-type specific expression. (**D**) Cross-species nucleotide contribution scores Spearman correlation between the DeepMouseBrain models and DeepChickenBrain for the top 100 DARs per chicken cell type. The median of the correlation per region is shown. The correlations are the consensus of two comparisons between DeepMouseBrain1&2 and DeepChickenBrain. Excitatory neuron, MSN and interneuron cell types are grouped in broad cell type categories. (**E**) Averaged contribution scores of characteristic sequence patterns are indicated for mouse (left) and chicken (right) astrocytes (AST), oligodendrocyte precursors (OPC), mature oligodendrocytes (OL), and microglia (MGL). The size of letters indicates their nucleotide contribution. Potential TFs that correspond to the sequence patterns are indicated based on known TFBS and the correlation of their expression with averaged importance scores. The scaled mean expression of TFs per cell type is shown by red circles. (**F**) Heatmap depicting the average number of instances of sequence patterns that are characteristic for broad cell type categories. Cell types of the different species are indicated in black, orange, and blue for mouse, chicken, and human cell types respectively.

Next, we used the nucleotide importance score correlation to compare the similarity of enhancer codes across all mammalian and avian telencephalic cell types. In particular, we calculated the median Spearman correlation between the nucleotide contribution scores of the top 100 DARs in chicken ranked by log-fold change and all of the per cell type contribution scores of the mouse and human models for those DARs. As expected, the correlation between contribution scores is higher for homologous cell types than for non-homologous cell types, also when the comparison is done on the top 100 mouse and human DARs (Fig. 4D, fig. S4, C to E). This strategy enables cell type comparison based on cell-type-specific nucleotide contribution scores, which represent the underlying gene regulatory code that the models learn.

To further investigate the specific TFBS per cell type learned by our models, we used TF-MoDISco (*42*) to infer cell type characteristic sequence patterns for each set of cell type specific DARs (log-fold change=1.5). To identify learned patterns that are shared across cell types and species, we compared the learned patterns of all models against each other using the MEME suite and clustered them into 42 groups of motifs (fig. S6-7) (*43, 44*). In addition, we screened all learned patterns against the *cistarget* database (*4*). The majority of learned patterns resembles TFBS (Data availability). The learned TFBS patterns correlate with differential TF expression across astrocytes, oligodendrocytes, and microglia (Fig. 4E). For example, concordant patterns and expression of genes are found for Nfia, Rorb, Rfx4, and Lhx2 in astrocytes, Sox10 and Olig2 in Oligodendrocytes; and Spi1 and Cebpa in microglia. In agreement with astrocyte specific TFBS that we identified with TF-MoDISco, the potential astrocyte enhancer region within the *PRDM16* intron harbors Nfia, Rorb, and Lhx2 binding sites that our models consider important for identifying this region as astrocyte specific (Fig. 4A).

To evaluate whether the learned TFBS motifs can be used to distinguish different cell types, we evaluated the correlation of their average number of instances across cell types and species to cluster cell types (fig. S8). We then performed hierarchical clustering based on the correlation of motifs between the human, mouse, and chicken cell types. The cell types of the three different species cluster into astrocytes, oligodendrocytes, microglia, interneurons, MSN, and excitatory neurons based on the correlation of learned TFBS motifs, which indicates shared enhancer codes across the species (fig. S7). Furthermore, broad cell type categories including MSN, interneurons, neurons of the hippocampus or medial pallium, and excitatory neurons of the neocortex and pallium can be characterized by a set of five TFBS motifs, which are shared between human, mouse, and chicken. (Fig. 4F). MEF2 binding sites are important to predict most neuronal cell types; and EGR TFBS are important for MSN and IT cortical or hyper-, nido-, or entopallial excitatory neurons (Fig. 4F). In contrast LHX/DLX binding sites are characteristic for astrocytes, and GABAergic neurons including MSN and interneurons. RFX binding sites were detected for astrocytes, MSN, neurons of the hippocampus or medial pallium, cortical layer L2/3, and chicken hyper-/nidopallium (EXC GLU-1). AP-1 factor family binding sites are learned by our models for predicting MSN and mammalian IT neurons, similar to previous findings (*13*). Hence, our models learned the same TFBS for the amniote species to distinguish broad neuronal cell type categories. The comparison of patterns thus represents a third strategy to employ enhancer models across species, besides the model prediction scores and the correlation of attribution scores.

### Enhancer codes of GABAergic neurons are conserved between birds and mammals

Using the same three strategies for comparing enhancer codes, we investigated avian and mammalian MSN and interneurons in more detail. In both mouse and chicken, lateral ganglionic eminence (LGE) derived D1/2MSN are exclusively found in the striatum, whereas medial (MGE) and caudal ganglionic eminence derived (CGE) interneurons occupy cortical areas in mammals and pallial areas in chicken (Fig. 5A). Two clusters of inhibitory neurons are preferentially mapped to the septum and globus pallidus in the chicken Stereo-seq data (fig. S9); structures that are not part of any of the analyzed mammalian data sets. The entopallium contains exclusively PVALB+ interneurons based on our mapping from single-cell to spatially resolved transcriptomics data. This is supported by the localized expression of *PVALB* in the entopallium (Fig. 5A). The transcriptomes of GABAergic neuron types can be clearly matched across mammals and birds supporting their previously suggested conservation across amniotes (Fig. 2C) (*22, 23*).

**Fig. 5.**
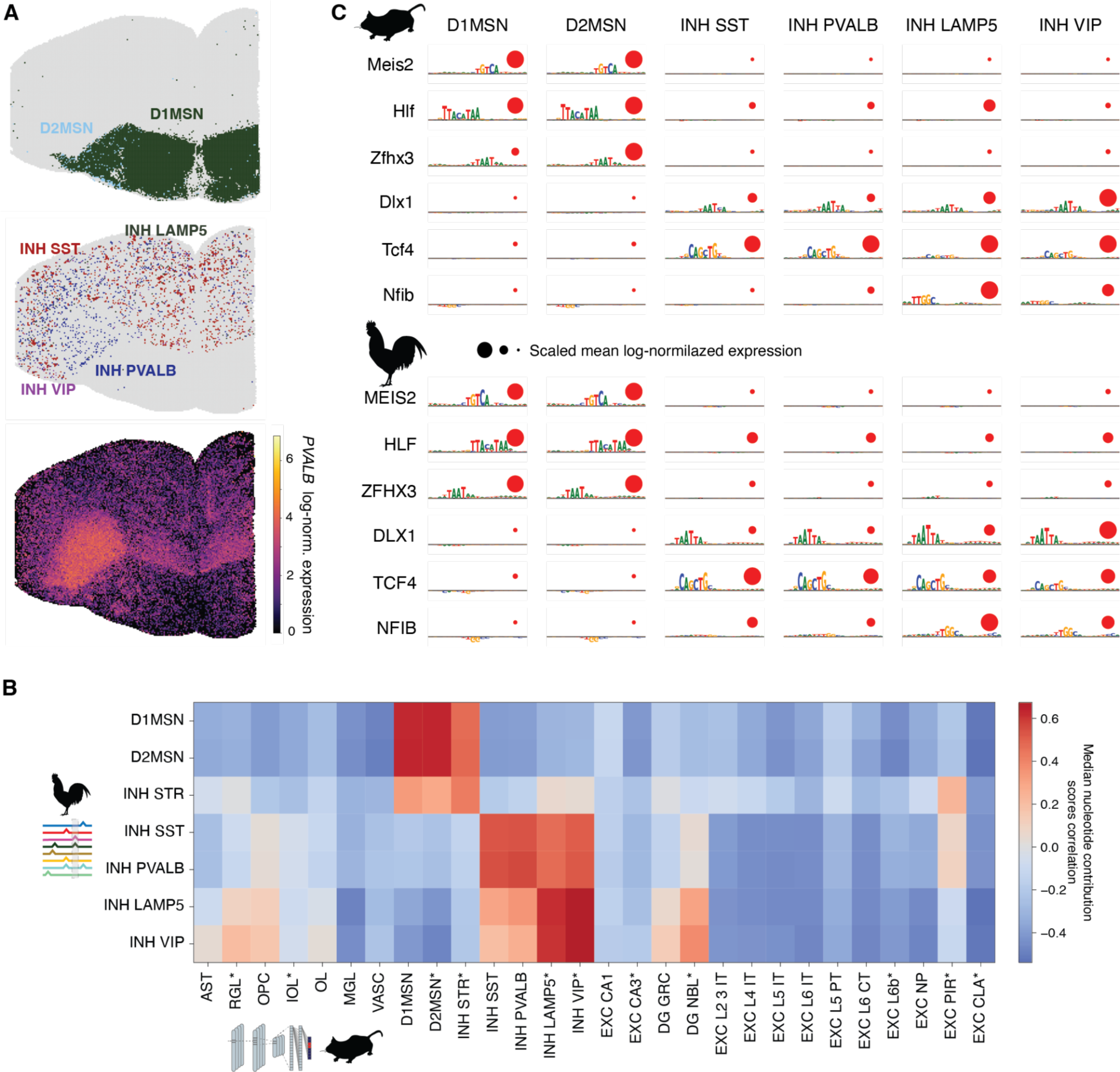
Conserved enhancer codes of GABAergic neurons across mouse and chicken. (**A**) Localization of MSN (top), interneurons (middle), and *PVALB* expression are shown in chicken spatially-resolved transcriptomics data (Stereo-seq) (bottom). Cell types are assigned to spatial spots based on the highest density of mapped single cell clusters. (**B**) Cross-species nucleotide contribution scores Spearman correlation between the DeepMouseBrain models and DeepChickenBrain for the top 100 DARs of medium spiny neurons and interneurons. The heatmap shows the consensus of the median correlation coefficients from the two comparisons between DeepChickenBrain and the two mouse models (DeepMouseBrain1&2). Mouse cell types annotated with an asterisk only contain contribution scores from DeepMouseBrain2 (Methods). (**C**) Averaged contribution scores of characteristic sequence patterns are indicated for mouse (top) and chicken (bottom) medium spiny and interneurons. The size of letters indicates their nucleotide contribution. Potential TFs that correspond to the sequence patterns are indicated based on known TFBS and the correlation of their expression with the averaged nucleotide importance scores. The scaled mean expression of TFs per cell type is shown by red circles.

To assess whether we can recapitulate the conservation of GABAergic neurons also at the level of enhancer codes, we investigated the predictions and learned features of our deep learning models. Note that the resolution of the ATAC signal limits the ability to distinguish between SST and PVALB interneuron types, and between LAMP5 and VIP at the level of chromatin accessibility (fig. S10). Nonetheless, our models are able to distinguish between D1/2MSN, MGE-derived and CGE-derived interneurons based on their prediction scores (fig. S11) and based on the correlation of derived nucleotide contribution scores (Fig. 5B).

Learned enhancer codes for predicting D1/2MSN and interneurons correspond to binding sites for key TFs and are correlated with TF expression (Fig. 5C). Particularly, Meis2 and Hlf binding sites are characteristic for spiny neuron enhancers. In contrast, basic helix-loop-helix (bHLH) factor binding sites likely corresponding to Tcf4 and binding sites of Nfib are characteristic for interneurons. Homeodomain factor binding sites, which could correspond to different DLX-family TFs or Zhfx3, are characteristic for all MSN and interneurons. Dlx1/2/5 are key TFs of cortical GABAergic neurons (*3*). As an example for a MSN specific putative enhancer, we identified a region in an intron of *FOXP2* that is specifically accessible in D1/2MSN and striatum-like inhibitory neurons (INH STR). This region is highly conserved in sequence (sequence identity: 94.9%) and ATAC signal (log-fold change 3.66 and 3.80) between mouse and chicken (Fig. 6A & B). *FOXP2* is a marker gene for D1/2MSN (Fig. 6A) and an important gene for plasticity in the striatum, as well as striatum-dependent learning of skills, including speech (*45, 46*). Mutations in *FOXP2* alter the length of MSN dendrites and have been linked to the evolution of language and speech deficits in humans (*47*). The mouse, human, and chicken models detect MEIS2 and homeodomain factor binding sites to be characteristic of D1/2MSN in agreement with the TFBS identified by TF-MoDISco (Fig. 4F). We assessed the activity of this enhancer candidate in vivo by incorporating it into an AAV vector along with a GFP-reporter and administering it to a mouse brain via injection (Methods). The reporter specifically expressed GFP in the striatum and striatum-like amygdalar nucleus, in agreement with our predictions and observed scATAC signal (Fig. 6D, fig. S12A and B, fig. S13). Additionally, there was a slight expression observed in the thalamus, a brain area absent from our current mouse datasets and models. Although the sequence is highly conserved, the chicken enhancer lacks an Egr1 binding site due to two point mutations compared to the mouse enhancer. As an example for an interneuron specific enhancer candidate, we identified regions near the interneuron-marker gene *ELAVL2*, that are specifically accessible for SST-interneurons in chicken and mouse, showing conserved enhancer codes of SST interneurons (fig. S14). Our analyses suggest that enhancer codes of GABAergic neurons are conserved between mammals and birds.

**Fig. 6:**
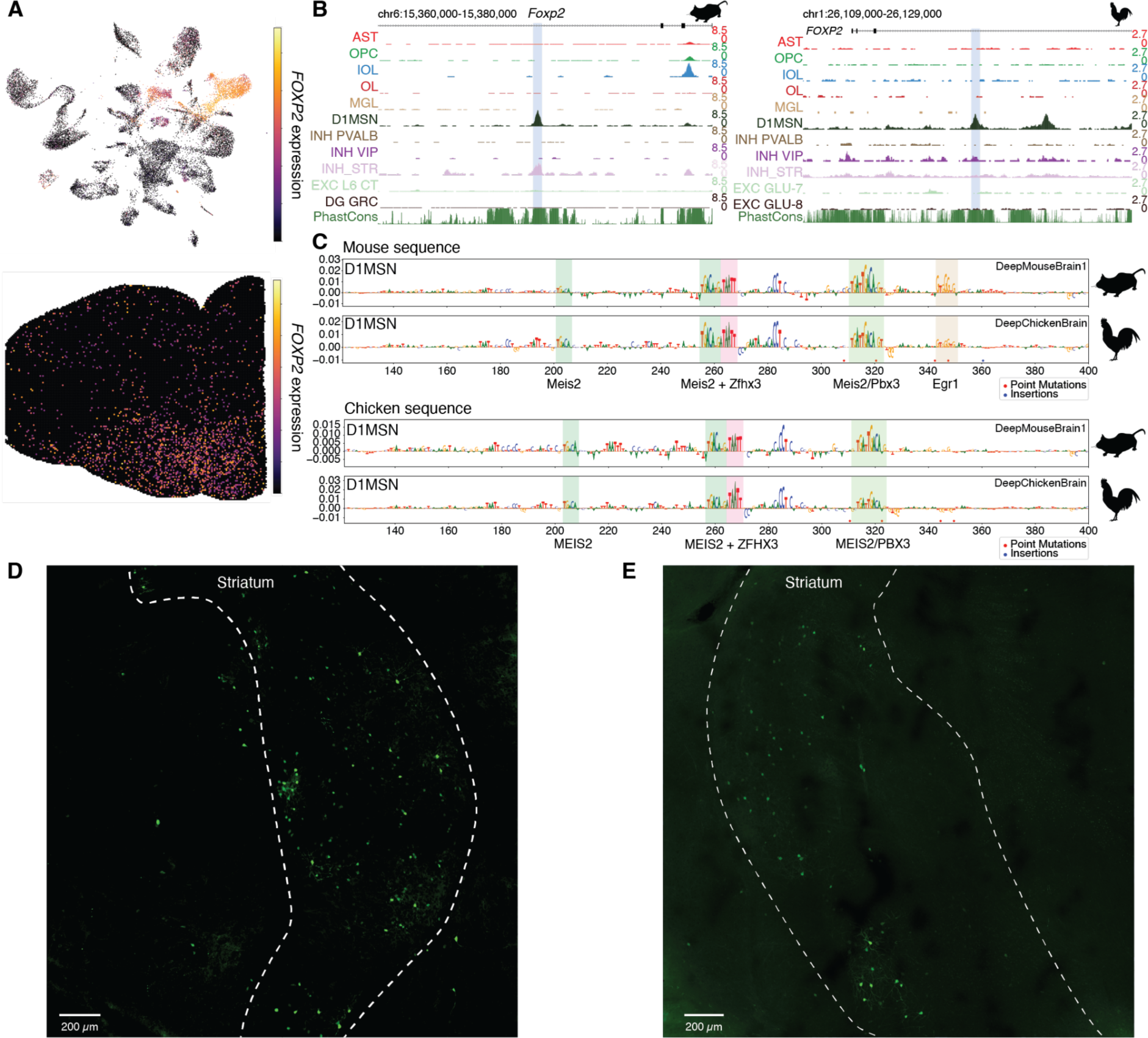
Identification of an intronic medium spiny neuron enhancer inside *Foxp2*. (**A**) scRNA-seq UMAP showing *FOXP2* expression is specific to D1MSN and striatal-like inhibitory neurons (INH STR) in chicken telencephalon single cell data (top), and highest expression in striatum in spatially resolved transcriptomics (Stereo-seq) data (bottom). (**B**) scATAC tracks of the *FOXP2* region in chicken (galGal6 chr1:26118953-26119453, left) and mouse (mm10 chr6:15369301-15369801, right) showing D1MSN specificity. (**C**) DeepMouseBrain1 and DeepChickenBrain contribution scores for their D1MSN classes for both the mouse (top) and chicken sequence (bottom); shown is the reverse complement of the sequence. Point mutations and insertions in the mouse sequence when aligned to the chicken sequence are indicated in red and blue respectively and vice versa in the chicken sequence. TFs that potentially bind to nucleotide features with high contribution scores are indicated and assigned based on their similarity to known TFBS motifs and cell-type specific expression. (**D**-**E**) GFP expression in the mouse brain driven by the D1/2MSN enhancer sequence within the mouse *Foxp2* intron (**D**) and the orthologous sequence within the chicken FOXP2 intron (**E**). The striatum is delimited by dashed lines.

### Enhancer codes suggest similarities between mammalian and avian excitatory neurons

In contrast to the conservation of GABAergic neurons, homologies between pallial avian and mammalian cortical excitatory neurons are highly debated (*48*). Building on our findings for non-neuronal and GABAergic neurons, we investigated whether the trained enhancer models provide additional insights into the conservation or divergence between excitatory neurons of birds and mammals.

We identified eight clusters of excitatory neurons in the chicken telencephalon that are localized to the hyper-, nido-, ento-, and medial pallium (Fig. 7A). The localization of these excitatory neuron cell types agrees with the subdivision of the avian pallium defined by the Avian Brain Consortium (*49*). The only exception being that we do not observe a split between hyper- and nidopallium in agreement with previous findings for zebra finch brains (*50*). Instead, the spatial transcriptomics data suggests two excitatory neuron cell types that are both distributed across the nido- and hyperpallium. Based on transcriptome comparisons using SAMap, excitatory neuron clusters of the chicken nido- and hyperpallium (GLU-1 and GLU-2) exhibit the highest similarities to mammalian IT neurons of cortical layers L2/3 and L4, excitatory neurons in the piriform cortex (PIR), and amygdala (Fig. 2C, fig. S1). The avian entopallium excitatory neuron transcriptome (GLU-3) shows a moderate similarity to mammalian L4 IT neurons and exhibits a characteristic expression of *RORB* as previously has been observed (*29*) (Fig. 5C). In contrast, transcriptomes of excitatory neurons of the chicken mesopallium (GLU-4/5/7) show the highest resemblances to excitatory neurons of the mammalian deeper layer cortical neurons including L5/6 IT neurons, L6 corticothalamic (CT) neurons, and neurons of the claustrum. Chicken excitatory neurons that are localized to the medial pallium (GLU-6) exhibit the highest similarities to mammalian hippocampal Cornu Ammonis (CA) neurons and the amygdala. The GLU-8 cluster in the chicken medial pallium shows the highest similarity to mammalian dentate gyrus (DG) neuroblasts (NBL). GLU-8 shows a characteristic expression of *SOX11*, a TF involved in neurogenesis in mammals (*51*) (Fig. 2B). A subpopulation of the GLU-8 cells localizes adjacent to the medial pallium while the entire cluster likely includes additional cells undergoing neurogenesis (Fig. 7A). As a positive control, we compared transcriptomes of mammalian cortical cell types between human and mouse. With the exception of L6 IT neurons, SAMap matches neuronal cell types of different cortical layers between human and mouse (fig. S1). While L2/3 IT neurons are properly matched they also exhibit some similarity to L5 extra-telencephalic (L5 PT) neurons. In contrast to mammalian excitatory neuron similarities, transcriptome comparisons by SAMap suggest 1:many or many:many homologies between excitatory neurons of the chicken and mammalian telencephalon. However, avian meso-, ento-, hyper/nidopallium, and medial pallium, show the highest similarities to either mammalian cortical deep layer, L4 IT, upper layer neurons or neurons of the hippocampus, respectively (Fig. 2C, fig. S1).

**Fig. 7.**
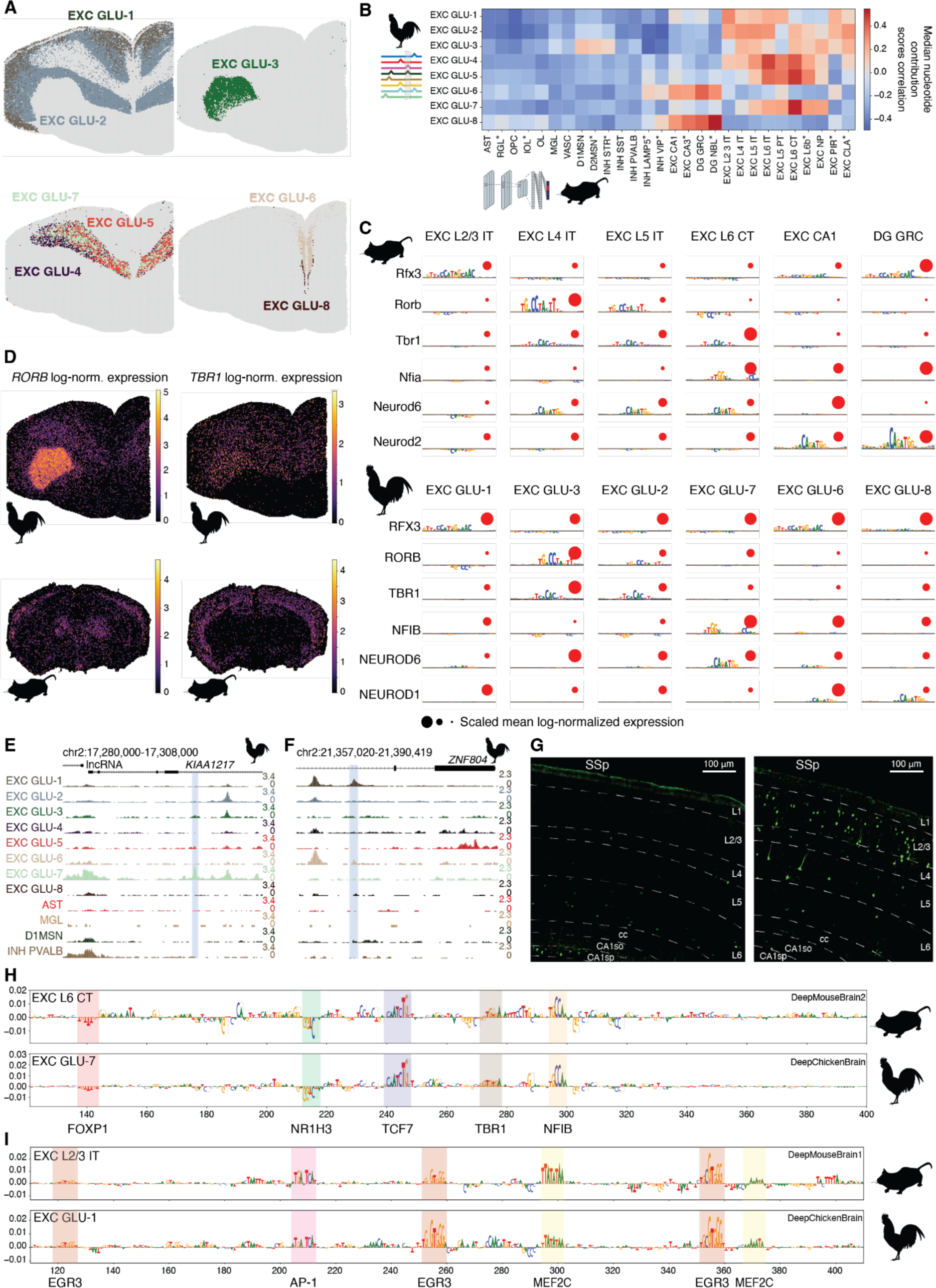
Enhancer code of excitatory neurons in chicken and mouse. (**A**) Chicken telencephalon excitatory neurons show a distinct localization to either the hyper- andnidopallium (top left), entopallium (top right), mesopallium (bottom left), or medial pallium (bottom right) in spatially resolved transcriptomics data (Stereo-seq). Spatial spots are assigned to cell types based on the highest density of mapped single cell clusters. (**B**) Median nucleotide contribution scores Spearman correlation heatmap for the top 100 DARs of the chicken glutamatergic neurons. Contribution scores were calculated by DeepChickenBrain for the glutamatergic classes and by the DeepMouseBrain models for all cell types. The correlation scores that were only scored with DeepMouseBrain2 are indicated with an asterisk. (**C**) Averaged contribution scores of characteristic sequence patterns are indicated for mouse (top) and chicken (bottom) excitatory neurons. The size of letters indicates their nucleotide contribution. Potential TFs that correspond to the sequence patterns are indicated based on known TFBS and their expression. The scaled mean expression of TFs per cell type is shown by red circles. (**D**) Expression of RORB (left) and TBR1 (right) is shown in chicken (top) and mouse (bottom) spatially-resolved transcriptomics data (Stereo-seq). RORB shows its highest expression in chicken entopallium and a characteristic expression in mouse layer 4 of the neocortex. Tbr1 expression is more widespread across the chicken pallium and neocortex but highest in chicken entopallium and mouse layer 6. (**E**) Chromatin accessibility profiles in the *KIAA1217* locus (galGal6 chr2:17294276-17294776) showing specific accessibility of EXC GLU-7. (**F**) Chromatin accessibility profiles in the *ZNF804B* locus (galGal6 chr2:21373470-21373970) showing specific accessibility of EXC GLU-1. (**G**) GFP expression driven by the *KIAA1217* (left) and the *ZNF804B* enhancer (right) in the mouse primary somatosensory cortex (SSp). The different cortical layers (L) are indicated, as well as the corpus callosum (cc) and the CA1 stratum oriens (CA1so) and pyramidal layer (CA1sp). (**H**) Nucleotide contribution scores for the chicken *KIAA1217* enhancer candidate scored by DeepMouseBrain2 and DeepChickenBrain for the classes of cell types EXC L6 CT and EXC GLU-7 respectively. TFs that potentially bind to nucleotide features with high contribution scores are indicated and assigned based on their similarity to known TFBS motifs and cell-type specific expression. **(I)** Nucleotide contribution scores for the chicken *ZNF804B* enhancer candidate scored by DeepMouseBrain1 and DeepChickenBrain for the classes of cell types EXC L2/3 IT and EXC GLU-1 respectively. TFs that potentially bind to nucleotide features with high contribution scores are indicated and assigned based on their similarity to known TFBS motifs and cell-type specific expression.

Next, we employed the DeepMouseBrain and DeepHumanCortex models to investigate whether enhancer codes suggest similar correspondences between excitatory neurons of the avian and mammalian pallium. For the top 100 DARs per excitatory neuron type in chicken, we calculated cross species predictions and nucleotide contribution score correlations with mouse and human cell types present in the DeepMouseBrain and DeepHumanCortex models (Fig. 7B, fig. S11A, fig. S15A). The correlation between nucleotide contribution scores largely recapitulates the evolutionary relationships between mouse and human excitatory neurons (fig. S15B and C). As was observed for excitatory neurons as a broad category (Fig. 4D), the enhancer code of chicken excitatory neuron clusters only correlates with mouse excitatory neurons. Chicken hyper- and nidopallium excitatory neurons (EXC GLU-1 and GLU-2) exhibit similarities to mammalian upper layer excitatory cortical neurons and neurons of the piriform cortex, whereas chicken mesopallium cluster EXC GLU-7 is most similar to mammalian deep layer neurons L6 CT and L6b (Fig. 7B, fig. S6). As we expected, chicken neurons of the medial pallium exhibit the highest similarity in nucleotide contribution score to mouse hippocampal or dentate gyrus cell types. Nucleotide contribution scores of chicken entopallium neurons (EXC GLU-3) are moderately correlated with mammalian L4 IT or L5 IT neurons, although this correlation is substantially lower than correlations observed between L6 CT and mesopallium cell types. As we observed for the transcriptome comparisons, the deep learning models suggest 1:many or many:many correspondences between the enhancer codes of most avian and mammalian excitatory neuron subclasses. In agreement with the transcriptome comparisons, the strongest correspondences of enhancer codes are between mammalian upper layer neurons or piriform cortex to avian hyper- or nidopallium; and L6 CT neurons to avian mesopallium neurons.

To investigate whether these correspondences are also reflected in learned TFBS, we identified TFBS patterns across species that are important for differentiating between excitatory neuron subclasses, in the same manner as we did for GABAergic and non-neuronal cell types above. Among others, the most characteristic learned motifs resemble binding sites of Rfx3, Rorb, Tbr1, and basic helix-loop-helix (bHLH) that likely correspond to the TCF-family, ATOH-family, or NEUROD-family TFs (Fig. 7C). Potential binding sites of Rfx3 are most characteristic for mammalian L2/3 IT, piriform cortex, avian hyper-/nidopallium neurons (EXC GLU-1), mammalian neurons of the hippocampus/dentate gyrus, and avian medial pallium, in agreement with their correlated enhancer codes (Fig. 7B, fig. S6). The learned importance of Rorb and Tbr1 motifs is correlated with their expression (Fig. 7C). *Rorb* expression is specific to L4 IT neurons in mice and to the chick entopallium, whereas *Tbr1* is more widely expressed across the mammalian neocortex and chicken pallium with highest expression in deeper layers of the neocortex and in avian entopallium (Fig. 7D). Potential binding sites of Tbr1 are learned to be most characteristic for L4/5 IT, L6 CT neurons, and excitatory neurons of the claustrum in mouse regions (fig. S16). For chicken regions, DeepChickenBrain learned potential TBR1 binding sites to be characteristic for the entopallium and nidopallial neurons of cluster EXC GLU-2. Both our DeepMouseBrain and DeepChickenBrain models learned bHLH TFBS likely corresponding to Neurod1/2 to be important for hippocampal/dentate gyrus and medial pallium neurons. While the DeepBrainChicken model suggest bHLH motifs, likely corresponding to TCF-family TFs, to be most characteristic for all excitatory neuron types of mesopallium, our mammalian models suggest them to be important for L4/5 IT and L6 CT neurons. Potential binding sites of Nfib are most characteristic for mammalian L6 CT and avian mesopallium neurons.

To validate the enhancer code-derived correspondences between GLU-7 and L6b and CT neurons, and between GLU-1 and L2/3 and PIR neurons, we inspected regions that are specifically accessible either in the avian mesopallium cell type GLU-7 or in the nido/hyperpallial excitatory neuron cluster GLU-1. A representative region for GLU-7 neurons is located approximately 10 kb upstream of the GLU-7 marker gene *KIAA1217* (Fig. 7E, fig. S17A). Characteristic of GLU-1, we identified a region that is specifically accessible in GLU-1 located near *ZNF804B* (Fig. 7F, fig. S18). We assessed the activity of both of these enhancer candidates with enhancer reporter assays in vivo in mouse brains (Fig. 7G). The *KIAA1217* enhancer shows a faint but specific expression in deep layer of the mouse neocortex, in agreement with the predicted correspondence to L6 CT and L6b cells by our models. In addition, the enhancer shows a strong activity outside of the cortex in the hippocampus and thalamus (fig. S12C). This activity pattern has been shown to be characteristic to AAV-PHP.eB infections and may not reflect cell type specific activity in the hippocampus (*52, 53*). In agreement with the patterns that we detected to be characteristic for mammalian L6 CT neurons and avian GLU-7 neurons, DeepMouseBrain and DeepChickenBrain suggest NFIB- and BHLH TFBS to be most important for the *KIAA1217* enhancer prediction (Fig. 7H). To verify that similar enhancer codes are characteristic for chicken mesopallium (GLU-7) and mouse L6 CT neurons, we investigated whether the ortholog of *KIAA1217* in mice, *Etl4*, contains a potential enhancer region that harbors similar TFBS as the potential *KIAA1217* chicken enhancer region. We indeed observed potential bHLH factor and Nfib binding site patterns similar to the ones in the chicken *KIAA1217* region inside of an intronic region of *Etl4* that is differentially accessible in mouse L6 CT neurons, and that has a high prediction score for the L6 CT class in DeepMouseBrain1&2 (fig. S17). Next, we tested the activity of the candidate GLU-1 enhancer near *ZNF804B* in the mouse brain. This chicken enhancer indeed drives the reporter GFP expression in L2/3 IT and PIR neurons (Fig. 7G, fig. S12D) as predicted by the enhancer models and transcriptome based comparisons GLU-1 to L2/3 IT and PIR correspondence. It contains several potential EGR1 and MEF2C binding sites, which are characteristic for excitatory neurons (Fig. 7I, Fig. 4F).

Overall, the analysis of enhancer codes and transcriptome comparisons suggest different degrees of similarities between particular avian and mammalian cell types. We present four approaches for computing cell type similarities: (i) transcriptome comparison, (ii) predictions from sequence-based deep learning models, (iii) correlation of derived nucleotide contribution scores, and (iv) similarities of TFBS motifs. To summarize the similarity between avian and mammalian cell types, we aggregated these four cell type similarity metrics into an average cell type similarity (Fig. 8, A and B, fig. S19). The combined similarity confirms one-to-one correspondences of non-neuronal cell types, MSN and interneurons. Excitatory neurons overall show many-to-many matches with the highest similarities found between GLU-7 and L6 CT, and GLU-1 and L2/3 IT neurons, suggesting a conserved enhancer code for these cell types.

**Fig. 8.**
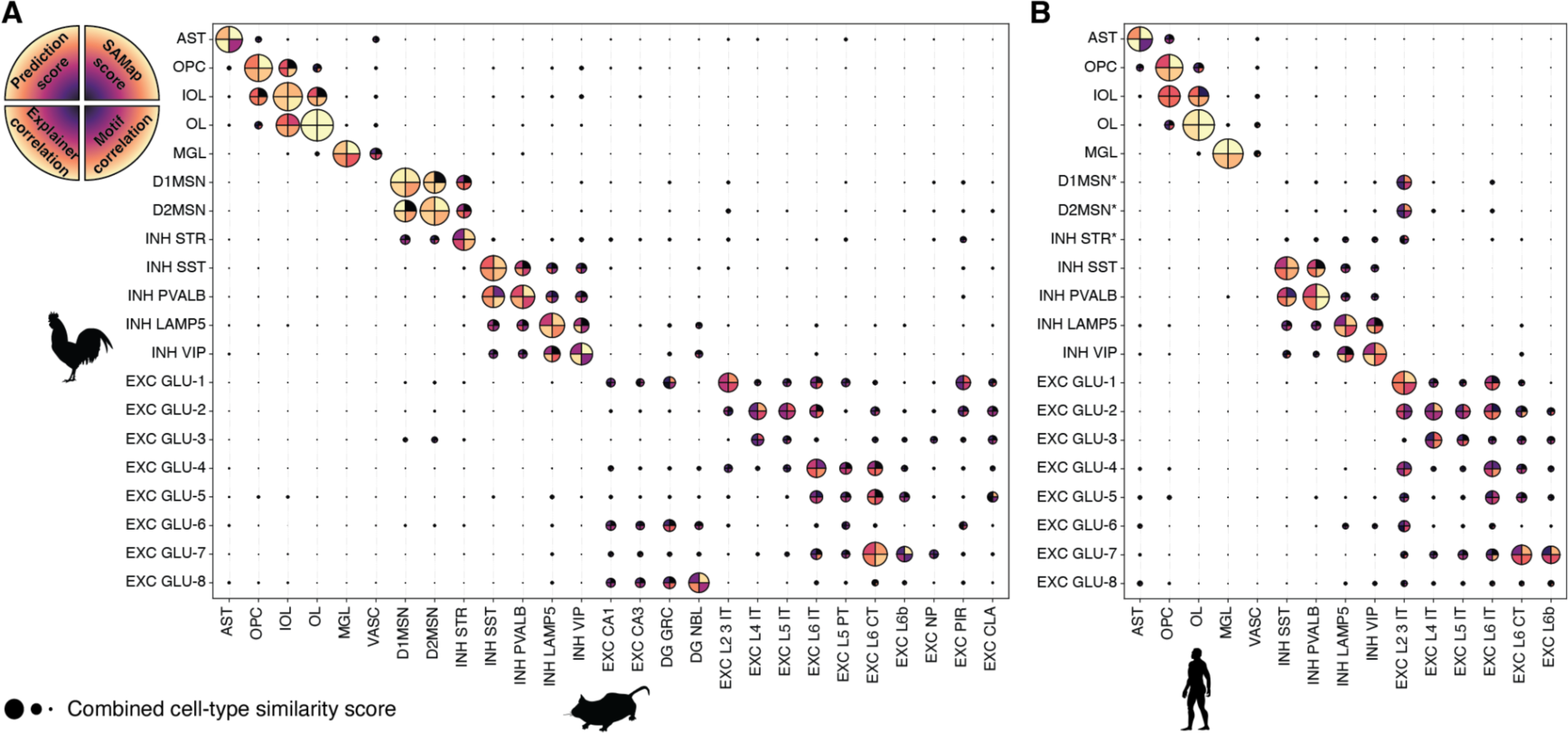
Overview of regulatory code matching methods between chicken-mouse and chicken-human brain cell types. (**A**) Combined cell-type similarity scores consisting of SAMap score, prediction scores, contribution score correlation and motif correlation between all available chicken and mouse cell types. The circle size depends on the mean score of all four metrics. SAMap scores are obtained from Fig. 2D. The predictions and contribution scores are the average median of scores on chicken regions with DeepMouseBrain models and scores on mouse regions with DeepChickenBrain. Motif correlation scores are merged for matching cell types in DeepMouseBrain1-DeepChickenBrain and DeepMouseBrain2-DeepChickenBrain. All scores were standardized between 0 and 1. Negative correlation scores were made zero. (**B**) Combined cell-type similarity scores between all available chicken and human cell types consisting of the same metrics as (A). SAMap scores are merged scores from the two human datasets against the chicken dataset. The predictions and contribution scores are the average median of scores on chicken regions with DeepHumanCortex models and scores on human regions with DeepChickenBrain. For VASC, the predictions and contribution scores correlations only come from DeepHumanCortex1 on chicken regions. Motif correlation scores are merged for matching cell types in DeepHumanCortex1-DeepChickenBrain and DeepHumanCortex2-DeepChickenBrain. All scores were standardized between 0 and 1. Negative correlation coefficients were set to zero. Asterisks indicate cell types which do not have correspondences to neocortical cell types based on our analysis. Consequently, enhancer code metrics only show low similarities to any cell type that is part of the human neocortex data sets.

## Discussion

We trained independent deep learning models on mammalian and avian genomic sequences to identify enhancer codes that are sufficient to identify major brain cell types, including astrocytes, microglia, oligodendrocytes, spiny neurons, interneurons, and excitatory neurons. These enhancer codes comprise TF binding site combinations and arrangements that are characteristic for the identity of the different cell types in the telencephalon and are conserved across mammals and birds. Our results agree with previous findings that TFs are the main components of conserved core regulatory circuits that define animal body plans and cell type identity (*31–33*). While gene expression and properties of homologous cell types may show larger variation between species, core sets of TFs have been found to show a high degree of conservation. For instance, developmental programs of cells in the retina are regulated by a set of key TFs that is conserved across vertebrates (*54, 55*). Also, the key TFs underlying heart development are conserved from the fruit fly to human (*56*). Genomic signatures of such TF combinations that form enhancer codes can be conserved across evolutionary distances as far as fishes and sponges (*57*). This conservation of enhancer codes is the foundation for training cell type specific deep learning models on accessible chromatin regions. Deep learning models have been successfully applied to predict chromatin accessibility for primate and mouse neocortical cell types indicating conservation of the mammalian neocortical enhancer code (*13, 14*).

We show here that enhancer codes are conserved across mammal and bird telencephalic cell types and present three computational strategies on how to utilize the conservation of enhancer codes to assess cell type similarities between species, as a complement to transcriptome-based similarities. These computational strategies only require the enhancer codes to be conserved, but the enhancer sequences themselves need not to be conserved. This enables our approach to be utilized for comparisons between evolutionary distant species such as mammals and birds even if very few or none of the enhancer regions are alignable (e.g., not projected via *liftOver*) between the species. While the applicability of our approaches is limited by the number of cells per cell type and the resulting resolution of clusters in scATAC-seq data, it shows a similar performance for cell types with a sufficient number of cells when compared to transcriptome-based comparisons (fig. S9). Advances in scATAC-seq technologies will resolve these limitations in future datasets, as the number of sequenced cells is increasing. Cell type annotation of single-cell data sets is often performed based on known marker genes, integration with reference atlases, or models trained on a subset of genes (*58*). For different species these analyses are limited to orthologous genes that either have to be specified or inferred with sophisticated tools such as SAMap or SATURN (*27, 59, 60*). For scATAC-seq data this usually requires relying on multiome data with both gene expression and ATAC-seq modalities. Our approach circumvents the need for an additional gene expression modality or known orthologs, once a model has been trained on cell types from an annotated reference species atlas. Hence, enhancer codes identified by deep learning models provide a means to compare cell types solely based on the genome sequence and chromatin accessibility.

While excitatory neurons, as a broad class, share characteristic enhancer codes between species, most of the excitatory neuron subclasses show 1:many or many:many homologies between mammals and birds based on their enhancer codes. Whether these correspondences of enhancer codes extend beyond brain regions that current data allows us to investigate such as the mammalian amygdala, entorhinal cortex, or avian arcopallium will be interesting to address in future studies. The similarities that we observed between avian and mammalian excitatory neurons neither agree with previous evolutionary models proposed based on shared developmental trajectories, nor with models based on vertebrate brain circuitry (*61–63*). Puelles et al. proposed the tetrapartite model for vertebrate brain homology by tracing the expression of a set of marker genes during development (*21*). The tetrapartite model suggests the avian hyperpallium (DPall) to be homologous to the mammalian neocortex but not the nidopallium (VPall or DVR) (*21*). In contrast, our enhancer code and transcriptome comparisons suggest that one of the excitatory neuron cell types of the chicken nido-/hyperpallium (EXC GLU-1) exhibits the highest similarity to mammalian L2/3 IT and PIR neurons. We validated this correspondence for a chicken enhancer, which shows activity in the upper layer of mouse neocortex and piriform cortex by enhancer reporter assays (Fig. 7G, fig. S12). The tetrapartite model has been questioned by Briscoe et al., who suggested that the cell types of the mammalian neocortex and non-mammalian pallium originated from a common amniote ancestor based on investigating bulk RNA-seq and in situ hybridization in late stage chick (E14) and alligator (stage 26) embryos, the closest extant bird relatives (*20, 64*).

The lack of cell types that resemble neocortical cell types in the amphibian telencephalon indeed suggests that the neocortical and reptilian/avian pallial neurons originated in the amniote ancestor (*65*). Our analysis of the chicken telencephalon supports the similarity between avian pallial and mammalian neocortical neurons. The evolutionary model by Briscoe et al. and previous transcriptome based comparisons suggest homology relationships between avian entopallium and mammalian L4 IT neurons (*20, 29*). While our analysis suggests some similarity between entopallium and L4 IT neurons, they are weaker than similarities observed between other pairs of cell types. The highest correspondence in enhancer code and transcriptome are surprisingly between a group of avian mesopallial neurons (EXC GLU-7) and mammalian L6 CT and L6b neurons. By contrast, Briscoe et al. suggested, based on gene expression and connectome similarities that the DVR, which includes the mesopallium, contains most similar neurons to the upper layer of the mammalian neocortex and the Wulst, which is part of the hyperpallium, to deep layer cell types in the neocortex (*64*). These differences to our findings may be in part attributed to investigating embryonic chicks and not juvenile chicken as analyzed in our study but are likely also due to the use of bulk RNA-seq. Bulk RNA-seq does not allow distinguishing between different cell types in the same tissue and their actual localization.

Three-dimensional imaging and tracing of pigeon brain fibers suggest that the avian telencephalon contains circuits that are similar to the ones of the mammalian neocortex (*19*). In contrast to enhancer code based similarities, circuit connectivities of mesopallial neurons resemble the ones of upper layer neocortical neurons, because neither have corticofugal projections (*19*). Given that the similarities between avian and mammalian telencephalic circuits do not correspond to assumed developmental origins, it has been suggested that they are a product of convergent evolution (*63*). Similar conclusions have been made for the vocal circuit of songbirds. By comparing single-cell RNA-seq and in situ hybridization, Colquitt et al. suggest that the songbird HVC in the posterior of the nidopallium resembles the mammalian piriform cortex, amygdala, and neocortex (*23*). While HVC excitatory neurons show widespread similarities to neurons of different neocortical layers based on the analysis by Colquitt et al., they also show their highest similarities to L2/3 IT neurons as our results suggest for nidopallial cell types.

Similar to our findings for birds, excitatory neurons of the turtle telencephalon do not have 1:1 homologs in mammalian neocortical cell types but their similarity is grouped by upper layer-like or deep layer-like neurons (*22*). Whereas we observe a clear separation of neocortex deep layer-like neurons in the mesopallium and upper layer-like neurons in the hyper- and nidopallium, turtle upper- and deep layer-like neurons are localized in the same layer of the reptilian dorsal cortex (*22*). As we find different degrees of conservation of enhancer codes across avian pallial and neocortical cell types that do not match the circuitry nor developmental origins, avian pallial excitatory neurons are likely either a product of convergent evolution or diversification and neofunctionalization as has been suggested for the evolution of the cerebellar nuclei in vertebrates and the hypothalamus in teleosts (*66, 67*). In line with the evolutionary model by Briscoe et al. (*20*), this means that neocortical neuron-like enhancer codes were present in cell types of the amniote ancestor and were either diversified or co-opted into different mammalian neocortical and avian pallial cell types.

In contrast to gene expression, enhancer codes can be directly traced across the genomes of related species to inform about evolutionary conservation. This provides a means to study cell type evolution through changes in candidate enhancers and the impact of genomic variants. As such, our models can be employed for studying how nucleotide changes are associated with cell type specificity as previously shown (*10, 15*). In past studies, we verified that enhancer codes for melanoma states are conserved between mammal and zebrafish cell lines and can be used to identify variants in the genomes of melanoma patients (*15, 68*). The models presented in this study can be used to complement efforts for studying the impact of genomic variants and their association with mental or cognitive traits and disorders (*69*). Ultimately, improved versions of our models hold the potential to screen genomes and annotate genomic loci by cell type specificity or to investigate the presence or absence of specific cell types or cell states.

## Methods

### Chicken telencephalon multiome

#### Tissue preparations

TRANSfarm (3360 Bierbeek, Belgium) provided chicken brains from healthy domestic chickens (P15, Gallus gallus/plofkip). Animal samples were handled according to KU Leuven ethical guidelines. The telencephalon was dissected from brains and either snap-frozen in liquid nitrogen or preserved for cryosectioning. To preserve the chicken telencephalons for cryosectioning, the whole telencephalon was frozen in isopentane precooled for 10 min in a beaker on dry ice. Coronal cryosections were dissected out at approximately 5 mm from the anterior of the brain and visually inspected for the presence of the lateral ventricle in parallel orientation to the longitudinal fissure.

#### Sample preparation

Two different sample types of chicken brain were used to isolate nuclei: dissected telencephalon snap-frozen in liquid nitrogen and a cryosectioned sample (12x 50 µm sections). The following procedure was followed to extract the nuclei: the tissues were resuspended in 300 µl nuclei lysis buffer comprising 10 mM Tris-HCl pH 7.4, 10 mM NaCl, 3 mM MgCl2, 0.1% Nonidet P40, 1 mM dithiothreitol and 1U/µl RNasin ribonuclease inhibitor (Promega) in nuclease-free water, pipeted until homogenized, 1 ml nuclei lysis buffer was added and samples were incubated on ice for 5 min. Nuclei were pelleted by centrifugation at 500 g for 5 min at 4°C and resuspended in 1x PBS, 1% BSA, 1U/µl RNasin ribonuclease inhibitor. Nuclei were centrifuged another time at 500 g for 5 min at 4°C and resuspended again in 1x PBS, 1% BSA, 1U/µl RNasin ribonuclease inhibitor. DAPI was added to the samples at a conc. of 0.5 µg/ml and incubated for 10 min. DAPI positive nuclei were sorted on a MA900 Sony cell sorter (Sony Biotechnology). The sorted nuclei were centrifuged at 500 g for 5 min at 4°C and resuspended in a 1 x nuclei buffer (10x Genomics) supplemented with 1 mM dithiothreitol and 1U/µl RNasin ribonuclease inhibitor.

#### Library generation

Single-cell libraries were generated using the 10X Chromium Single-Cell Instrument and NextGEM Single Cell Multiome ATAC+Gene Expression kit (10X Genomics) according to the manufacturer’s protocol. In brief, the single nuclei of chicken brains were incubated for 60 min at 37°C with a transposase that fragments the DNA in open regions of the chromatin and adds adapter sequences to the ends of the DNA fragments. After generation of nanoliter-scale gel bead-in-emulsions (GEMs), GEMs were incubated in a C1000 Touch Thermal Cycler (Bio-Rad) under the following program: 37°C for 45 min, 25°C for 30 min, and hold at 4°C. Incubation of the GEMs produced 10x barcoded DNA from the transposed DNA (for ATAC) and 10x barcoded, full-length cDNA from poly-adenylated mRNA (for GEX). This was followed by a quenching step that stopped the reaction. After quenching, single-cell droplets were broken and the transposed DNA and full-length cDNA were isolated using Cleanup Mix containing Silane Dynabeads. To fill gaps and generate sufficient mass for library construction, the transposed DNA and cDNA were amplified via PCR: 72°C for 5 min; 98°C for 3 min; seven cycles of 98°C for 20 s, 63°C for 30 s, 72°C for 1 min; 72°C for 1 min; and hold at 4°C. The pre-amplified product was used as input for both ATAC library construction and cDNA amplification for gene expression library construction. Illumina P7 sequence and a sample index were added to the single-strand DNA during ATAC library construction via PCR: 98°C for 45 s; 7–9 cycles of 98°C for 20 s, 67°C for 30 s, 72°C for 20 s; 72°C for 1 min; and hold at 4°C. The sequencing-ready ATAC library was cleaned up with SPRIselect beads (Beckman Coulter). Barcoded, full-length pre-amplified cDNA was further amplified via PCR: 98°C for 3 min; 6–9 cycles of 98°C for 15 s, 63°C for 20 s, 72°C for 1 min; 72°C for 1 min; and hold at 4°C. Subsequently, the amplified cDNA was fragmented, end-repaired, A-tailed, and index adaptor ligated, with SPRIselect cleanup in between steps. The final gene expression library was amplified by PCR: 98°C for 45 s; 5–16 cycles of 98°C for 20 s, 54°C for 30 s, 72°C for 20 s, 72°C for 1 min; and hold at 4°C. The sequencing-ready GEX library was cleaned up with SPRIselect beads.

Sequencing

Before sequencing, the fragment size of every library was analyzed using the Bioanalyzer high-sensitivity chip. All 10x scATAC libraries were sequenced on a NextSeq2000 instrument (Illumina) or on a NovaSeq6000 instrument (Illumina) with the following sequencing parameters: 51 bp read 1 – 8 bp index 1 – 24 bp index 2 – 51 bp read 2 and for 10x GEX libraries the following sequencing parameters were used: 28 bp read 1 - 10 bp index 1 - 10 bp index 2 - 90 bp read 2.

### Chicken telencephalon spatially resolved transcriptomics (Stereo-seq)

All the spatial analysis performed in work was done using BGI’s STomics platform. The sections below detail the methods used for the spatial library preparation and sequencing.

#### Tissue preparation for cryo-sectioning

All animal experiments were performed according to KU Leuven ethical guidelines. Healthy domestic chickens (P15, Gallus gallus/plofkip) provided by TRANSfarm (3360 Bierbeek, Belgium) were used for the experiments. Brains were dissected out and immediately snap-frozen in iso-pentane for 10 min. Afterwards, brains were embedded in Tissue-Tek OCT cryo embedding compound. 10 µm coronal cryosections were done at CT=14°C, OT=11°C. The used area of chicken telencephalon cryosections is the same as described above. 5-10 OCT scrolls of 70 um section thickness were collected into DNA lo-bind 2 ml Eppendorf tubes. Ice cold PBS was used for washing the tissue to remove the OCT matrix from the tissue. Total RNA was extracted from the washed tissue sections using the innuPREP mini RNA kit (Analytik Jen; Cat. No. AJ 845-KS-2040250). The RNA quality was assessed using RNA Nano kit (Agilent). Only tissue with RIN > 7 was used for the spatial analysis.

#### Optimization of tissue permeabilization

The tissue optimization was performed using the Stereo-seq Permeabilization kit (Cat. No. 111KP118) and Stereo-seq chip set P (Cat. No. 110CP118) according to the manufacturer’s protocol (Stereo-seq permeabilization set user manual, Ver A1). Briefly, 4 permeabilization chips were removed from the storage buffer and washed with nuclease free water and dried at 37*C. Next, 4 consecutive 10 µm tissue sections were prepared from the tissue cryo-block and placed on the permeabilization chip and thawed the tissue layer to attach to the surface of the chip. After drying the tissue on 37*C hot plate, the chip is then dipped into 100% methanol at -20*C and incubated for 30 mins to fix the tissue. Post fixation the tissue permeabilization test is performed on these chips by permeabilizing the tissue with PR enzyme prepared in 0.01N HCl (pH 2.0), at 4 different time points ranging from 6 mins to 30 mins. After the permeabilization, the chips were rinsed with 0.1X SSC buffer before reverse transcription. Reverse transcription is carried out at 42*C for 1 hr in the dark. Tissue removal is performed at 55*C for 1 hr using the TR enzyme to clear the tissue before imaging. Fluorescence imaging is performed in the TRITC channel with 10X objective, following the imaging guidelines provided by the manufacturer (Guide book for Image QC & microscope assessment and imaging, Ver A5). The optimal permeabilization time is assessed based on the strongest fluorescence signal with the lowest signal diffusion (crispness of the RNA footprint). Based on our assessment, we found the most optimal permeabilization time for the mouse brain to be X mins and for the chicken brain section to be Y mins.

#### Spatial Transcriptomics analysis

The spatial transcriptomics analysis was performed using the Stereo-seq Transcriptomics kit (Cat. No. 111ST114) according to the manufacturer’s protocol (Stereo-seq Transcriptomics set user manual, Ver A2). Briefly, similar to the permeabilization analysis, the T-chip was removed from the storage buffer and washed with nuclease free water and dried at 37*C. Next, a 10 µm tissue section from a desired region of interest was prepared from the tissue cryo-block and placed on the T-chip and thawed the tissue layer to attach to the surface of the chip. After drying the tissue on a 37°C hot plate, the chip is then dipped into 100% methanol at -20°C and incubated for 30 mins to fix the tissue. The fixed tissue is then stained using the Qubit ssDNA reagent (Thermo Cat. No. Q10212). Fluorescence imaging of the single stranded DNA staining is performed in the FITC channel with 10X objective, following the imaging guidelines provided by the manufacturer (Guide book for Image QC & microscope assessment and imaging, Ver A5). Prior to permeabilization, the ssDNA-stained image is also subjected to QC analysis using the imageQC software as per manufacturer’s recommendation. Like the permeabilization protocol, the tissue permeabilization is carried out with PR enzyme prepared in 0.01N HCl (pH 2.0) at 37*C. The optimal permeabilization time estimated from the tissue permeabilization analysis is used for the transcriptomics analysis. After washing the chip, reverse transcription mix is added to the chip and incubated at 42*C for at least 3 hrs. Tissue removal from the stereo seq chip is achieved by incubating the chip in TR buffer at 55*c for 10 minutes. cDNA release and collection is performed by incubating the chip in cDNA release mix overnight at 55*C and the release cDNA is purified with Ampure XP beads (Beckman Coulter; Cat. No. A63882) using the manufacturer’s recommendation. After the quality assessment using bioanalyzer (Agilent), sequencing library preparation is performed using transposase assisted tagmentation reaction. Indexed PCR and library purification is performed to prepare the final sequencing library as per manufacturer’s recommendations. Final Stereo-seq libraries were sequenced on MGI/BGI sequencing platforms and were sequenced at the MGI Latvia sequencing facility.

#### Analysis of chicken telencephalon single cell multiome data

Raw reads were mapped to the chicken genome assembly galGal6 (GRCg6a) with Cell Ranger ARC (v2.0.2, 10x Genomics) using an ENSEMBL 108 gene annotation as reference (*70*). Since ENSEMBL changed their main chicken assembly to bGalGal1 (GRC7b) and did not update all the gene names for the transcripts specified for the galGal6 assembly, we projected gene names from bGalGal1 to galGal6. In particular, we used liftOver to project exons from the bGalGal1 to GRCg6a assembly and assigned gene names based on the overlap of exons that were projected to the same strand (at least 85% of the nucleotides) (*71*). In addition, unclear assignments such as different gene names between the assemblies, splits, and merging of genes were manually curated.

The gene expression modality of the scMultiome was processed in SCANPY using the VSN pipelines (https://github.com/vib-singlecell-nf/vsn-pipelines, revision: 6a9c70769b) (*72*). Cells were filtered by a maximum 5% mitochondrial genes, a minimum of 200 and a maximum of 4,500 genes per cell. Genes that were present in less than three cells were removed. Doublets were detected with Scrublet and assigned based on the expected doublet rate for 10x Chromium protocols as implemented in the VSN Scrublet pipeline (*73*). Gene expression counts were normalized per cell to a target sum of 10,000 and log-transformed. 50 principal components were used after principal component analysis. Sample batch effects were corrected with Harmony (*74*). Clusters were identified with Louvain clustering using a resolution of 2 (*75*). Clusters were labeled as non-neuronal cell types, medium spiny neurons, and interneurons based on the expression of known marker genes. Clusters with less than 100 cells were removed. Additional clusters were annotated as excitatory based on expression of *SLC17A6* and inhibitory based on expression of *GAD1* or *GAD2*. Clusters that were assigned to the same cell type were merged. In addition, we used SAMap (version 1.0.2) to project cell types labels from mouse brain snRNAseq (see Methods below) to the chicken telencephalon scMultiome (*27, 30*). As input for SAMap, we compared the coding DNA sequences of all chicken transcripts against all mouse transcripts with tblastx and selected the transcript pairs with the highest BLAST score per gene pair (*76*). Cell types/clusters were specified as annotation keys and raw counts were used when running SAMap.

The ATAC-seq modality of the scMultiome was processed with pycisTopic (*4*). Pseudobulks were generated based on assigned clusters from the gene expression modality. Barcodes with less than 1,000 unique fragments, a fraction of reads in peaks (FRIP) lower than 20% and a TSS-enrichment below 2 were removed. In addition, only barcodes were kept that satisfied the filtering criteria of the gene expression modality and that were not considered doublets. 100 topics were selected for topic modeling based on topic metrics. Sample batch effects were corrected with Harmony (*74*). Up to the top 5,000 regions with the highest probability per topic were extracted after performing Otsu-thresholding. Differentially accessible regions (DARs) per cell type were computed with pycisTopic, i.e. a two-sided Wilcoxon rank-sum test on imputed chromatin accessibility. The imputed accessibility of all cells of per cell type is compared against the background composed of all other cells. Regions with a log-fold change of log_2_(1.5) and a Benjamini-Hochberg-adjusted p-value of at least 0.05 were considered for further analysis.

#### Analysis of chicken telencephalon Stereo-seq data

Stereo-seq data was processed with Stereopy (version 0.8.0) using bin-size=100 (*77*). Bins that express less than 20 genes and genes that were expressed in less than 3 bins were removed. Gene expression counts were normalized per cell to a target sum of 10,000 and log-transformed. Cell2location (version 0.1) was used to project labels of single cell clusters to bins in the Stereo-seq data (*30*). Raw counts were used as input and the single cell data was filtered with cell2location’s internal filtering function (using the following parameter: cell_count_cutoff=5, cell_percentage_cutoff2=0.03, and nonz_mean_cutoff=1.12). For mapping labels of single cell clusters to spatial bins, we set the number of expected cells per bin to 5.

## Analysis of mammalian brain data

### Mouse brain snRNA-seq

Available 10X Chromium mouse brain single nucleus was downloaded from https://cell2location.cog.sanger.ac.uk/tutorial/mouse_brain_snrna (Nov 2020) (*30*). Data was processed using VSN-pipelines (v0.24.0). Briefly, cells with at least 500 expressed genes, less than 7,000 expressed genes and less than 5% mitochondrial reads were kept. This resulted in a data set with 40,238 cells. Scanpy (v0.5.2) was run with default parameters, using 50 principal components, and using Leiden clustering with resolution 1.2. Clusters were manually annotated based on the expression of marker genes. IT neurons of the different layers were assigned based on known expression patterns from in situ experiments and by using a higher Leiden clustering resolution of 5 (*78*). Clusters were then merged per cell type and annotations were made consistent with the other analyzed single cell data sets.

### Mouse brain snATAC

Available 10X Chromium single nucleus ATAC-seq data of the mouse brain was obtained from the NeMO Data Archive (https://data.nemoarchive.org/biccn/grant/u19_cemba/cemba/epigenome/sncell/ATACseq/m <u>ouse</u>, Jun 2020) (*25*). Fragment files were processed with pycisTopic using the *L2cluster* annotation specified in the metadata for generating pseudobulk profiles and identifying consensus accessible regions (*4*). Barcodes with less than 1,000 unique fragments, a fraction of reads in peaks (FRIP) lower than 20% and a TSS-enrichment below 2.5 were removed. In addition, only barcodes with the following region identifiers were kept: 5D, 5E, 6D, 8B, 9J, 9H. This resulted in a data set with 106,462 cells. 225 topics were selected for topic modeling based on topic selection metrics. Up to the top 6,000 regions with the highest probability per topic were extracted after performing Otsu-thresholding. We then re-named cell type annotations from the *L2cluster* definition to match cell type annotations of the other data sets that we analyzed. DARs per cell type were computed with pycisTopic as mentioned above.

### Mouse brain scMultiome

We used previously generated 10X multiome data of mouse cortex, hippocampus and striatium; see Bravo González-Blas and De Winter et al. for details (*4*). scRNA-seq data were first analyzed using VSN (v.0.27.0). Briefly, cells with at least 100 genes expressed and less than 1% of mitochondrial reads were kept. Doublets were removed using Scrublet (v.0.2.3), with default parameters. 50 PCs were used as input for harmony, which was used to correct batch effects due to the sample preparation protocol and the corrected PCs were used for dimensionality reduction and Leiden clustering (resolution 1). This resulted in 41 clusters that were annotated based on marker gene expression. Clusters were per cell types and the nomenclature was made consistent with the other analyzed single cell data sets.

The ATAC-seq modality was processed in pycisTopic; see Bravo González-Blas and De Winter et al. for details (*4*). Briefly, the RNA-seq cell-type labels were used to create pseudobulks from which peaks were called with MACS2 (v.2.1.2.1) and consensus peaks were derived using the iterative-filtering approach. The data set was further filtered based on the scATAC-seq quality, keeping cells with at least 1,000 fragments, FRiP > 0.4 and TSS > 4. 125 topics were selected based on topic selection metrics. Up to the top 5,000 regions with the highest probability per topic were extracted after performing Otsu-thresholding. DARs per cell type were computed with pycisTopic as mentioned above.

### Mouse brain Stereo-seq

Available mouse brain Stereo-seq data was processed with Stereopy (version 0.8.0) using bin-size=100 (GEO-accession: GSE256319) (*77, 79*). Gene expression counts were normalized per cell to a target sum of 10,000 and log-transformed.

### Human motor cortex SNARE-seq

Available SNARE-seq2 data of the human motor cortex was downloaded from the NeMO Data Archive (*24*). High-quality cells (84,159) selected by Bakken et al. were used for the analysis. scRNA-seq data were analyzed using Seurat (v.4.0.3), using 47 PCs for dimensionality reduction and Leiden clustering (with resolution of 0.6). This resulted in 30 clusters (corresponding to 19 cell types) that were manually annotated based on marker gene expression and cell type nomenclatures were made consistent across analyzed single cell data sets.

The ATAC-seq modality was processed in pycisTopic; see Bravo González-Blas and De Winter et al. for details (*4*). 75 topics were selected for topic modeling based on topic selection metrics. Up to the top 5,000 regions with the highest probability per topic were extracted after performing Otsu-thresholding. We removed two topics which contained less than 100 regions after Otsu-thresholding from further analysis. DARs per cell type were computed with pycisTopic as mentioned above with the exception of adjusting the background. Since the fraction of cell types shows a high bias towards L2/3 IT neurons, which heavily affects the composition of the background for testing for differential accessibility, we randomly downsampled the background to a maximum of 3,000 cells per cell type when computing Wilcoxon rank-sum tests.

### Human prefrontal cortex multiome

Available 10X multiome data of the human prefrontal cortex was obtained from the NCBI Gene Expression Omnibus (GEO-accession: GSE207334) (*26, 80*). We limited our analysis to the following for samples: HSB8050, HSB5871, HSB6195, HSB6154, HSB8073. Raw reads were retrieved from the NCBI Sequence Read Archive using the SRA toolkit (https://github.com/ncbi/sra-tools) and fastq-dump with the following parameters: --include-technical --split-files Reads were mapped to the human genome assembly hg38 (GRCh38) with Cell Ranger ARC (v2.0.2, 10x Genomics) using an ENSEMBL 108 gene annotation (*70*). The gene expression modality of the scMultiome was processed in SCANPY using the VSN pipelines (https://github.com/vib-singlecell-nf/vsn-pipelines, revision: 6a9c70769b) (*72*). Cells with less than 500 genes and genes that were expressed in less than three cells were removed. Doublets were detected with Scrublet and assigned based on the expected doublet rate for 10x Chromium protocols as implemented in the VSN Scrublet pipeline (*73*). Gene expression counts were normalized per cell to a target sum of 10,000 and log-transformed. Sample batch effects were corrected with Harmony using 50 PCs as input (*74*). Clusters were identified with Leiden clustering using a resolution of 2 (*81*). Cell types were assigned to clusters based on the expression of known marker genes, following known expression from in situ experiments in the human cortex for neurons (*78*). To corroborate our assignments, we used SAMap (version 1.0.2) to project cell types labels from mouse brain snRNAseq (see Methods below) to the human cortex scMultiome (*27, 30*). Prior to running SAMap the mouse brain snRNAseq data set was subset to cell types that are present in the neocortex. As input for SAMap, we compared the coding DNA sequences of all human transcripts against all mouse transcripts with tblastx and selected transcript pairs with the highest BLAST score per gene pair (*76*). Cell types/clusters were specified as annotation keys and raw counts were used when running SAMap.

The ATAC-seq modality of the scMultiome was processed with pycisTopic (*4*). Pseudobulks were generated based on assigned clusters from the gene expression modality. Barcodes with less than 1,000 unique fragments, a fraction of reads in peaks (FRIP) lower than 25% and a TSS-enrichment below 3 were removed. In addition, only barcodes were kept that satisfied the filtering criteria of the gene expression modality and that were not considered doublets. 100 topics were selected for topic modeling based on topic metrics. Sample batch effects were corrected with Harmony (*74*). Up to the top 5,000 regions with the highest probability per topic were extracted after performing Otsu-thresholding. DARs per cell type were computed with pycisTopic as mentioned above with the exception of adjusting the background. Since the fraction of cell types shows a high bias towards L2/3 IT neurons, which heavily affects the composition of the background for testing for differential accessibility, we randomly downsampled the background to a maximum of 1,500 cells per cell type when computing Wilcoxon rank-sum tests.

### Correlation of gene ortholog expression

To compare the transcriptome of chicken and mouse telencephalon cell types with an alternative to SAMap, we evaluated the correlation of 1:1 gene ortholog expression following the approach by Tosches et al. (*22*). For this purpose, we obtained 1:1 orthologs of protein-coding genes between chicken and mouse genes from ENSEMBL that were annotated as *confident* orthologs (*70*). We then computed the mean log-normalized expression per cell type in the chicken gene expression data and mouse brain snRNAseq (*30*). We converted mean expression values into z-scores per the cell types and computed Spearman correlation coefficients of the z-scores over the 1:1 ortholog pairs for all pairwise combinations of chicken and mouse cell types. To evaluate whether TFs alone are meaningful for comparing cell types, we performed the same analysis with 1:1 TF orthologs instead of all orthologous genes.

## In vitro saturation mutagenesis

### Library generation

The saturation mutagenesis library used the mouse FIRE enhancer (mm10 chr18:61108596-61108855) as initial sequence. The 259 bp sequence was mutated in silico to generate all possible sequences with a single nucleotide mutation resulting in 777 sequences. Nine copies of the wild type FIRE sequence are included in the library to reduce the risk of losing it during cloning and MPRA. 500 shuffled sequences are used as negative controls. A unique 11 bp barcode generated via FreeBarcodes (*82*) was added in 5’ of each sequence and the adapters CCAGTGCAAGTGCAG and GGCCTAACTGGCCGG were added in 5’ and 3’ respectively. The library was produced by Twist Bioscience and PCR amplified with the Kapa HiFi HotStart ReadyMix (Roche) and the primers CCAGTGCAAGTGCAG and CCGGCCAGTTAGGCC. A barcoded version of the plasmid pSA351_SCP1_intron_eGFP (Addgene #206906) containing a 17 bp random barcode was linearized via inverted PCR with the Kapa HiFi HotStart ReadyMix and the primers GGCCTAACTGGCCGGCTGAGCTCCCTAGGGTACT and CTGCACTTGCACTGGCGACTCGAGGCTAGTCTC, followed by DpnI digestion and gel extraction in a 0.8% agarose gel. The amplified library and linearized plasmid were combined in an NEBuilder (New England Biolab) reaction with a vector to insert ratio of 1:2 for 45 min at 50°C then dialyzed against water in a 6 cm Petri dish with a membrane filter MF-Millipore 0.05 µm (Merck, Kenilworth, NJ) for 1 hour. The reaction was then transformed into Lucigen Endura ElectroCompetent Cells (Biosearch Technologies). Before culture for maxiprep, 1:100,000 of the transformed bacteria was plated onto an LB-agar dish with carbenicillin to estimate the complexity of the cloned library. A volume of bacteria corresponding to a complexity of 500 barcodes per enhancer was put in culture for maxiprep. Maxiprep was performed using the Nucleobond Xtra endotoxin-free maxiprep kit (Macherey-Nagel).

The enhancer to random barcode assignment was performed as described previously (*11*). Briefly, the cloned library was amplified with primers TCGTCGGCAGCGTCAGATGTGTATAAGAGACAGTCCCCAGTGCAAGTGCAG and GTCTCGTGGGCTCGGAGATGTGTATAAGAGACAGCTGGCCCTCGCAGACA. Illumina sequencing adapters were added during a second round of PCR with primers AATGATACGGCGACCACCGAGATCTACACNNNNNNNNTCGTCGGCAGCGTCAGATGTG and CAAGCAGAAGACGGCATACGAGATNNNNNNGTCTCGTGGGCTCGGAGATG*T. Reads were trimmed using Cutadapt (v.4.2) (*83*) with the options -g TCCCCAGTGCAAGTGCAG --discard-untrimmed -m 11 -l 11 for read 1 to extract the enhancer barcode and options -g CTGGCCCTCGCAGACA…GATCGGCGCGCCGGTCC -- discard-untrimmed -m 17 -M 17 for read 2 to extract the plasmid barcode. Reads were filtered to retain only those with quality > 30 using fastp (v.0.23.2) (*84*).

To produce lentivirus particles, HEK 293T cells cultured at ∼80% confluency in a 15 cm dish were transfected by using Lipofectamine 3000 reagent (Thermo Fisher Scientific) together with 30 μg cloned enhancer library, 20 μg psPax2 (Addgene #12260), 10 μg pMD2.G (Addgene #12259). Forty-eight and seventy-two hours post-transfection, medium was collected, combined, and spun down 5 min at 1,500 rpm. The supernatant was carefully collected with a blunt needle and a syringe and filtered through a 45 μm syringe disc filter (Millex - HV Millipore) into an Ultra-15 MWCO100 centrifugal filter (Amicon). The concentrator tube containing 15 ml of the supernatant was spun down at 4,000 rpm for approximately 45 min until the desired volume was reached (∼250 μl).

### Massively Parallel Reporter Assay

Mouse BV2 cells were grown in high glucose Dulbecco’s modified Eagle’s medium (Gibco) supplemented with 10% FBS (Gibco) and were maintained at 37°C and 5% CO_2_ atmosphere. Cells were seeded in a 10 cm dish and transduced with 100 µl of the lentivirus library once they had reached ∼80% confluency. 48 hours post transduction, cells are harvested with trypsin (Gibco). One-fifth of the cells are used to perform genomic DNA (gDNA) extraction with the DNeasy Blood & Tissue Kit (Qiagen). The rest of the cells are used to perform RNA extraction with the innuPREP RNA Mini Kit 2.0 (Analytik Jena), followed by mRNA isolation with the Dynabeads mRNA purification kit (Ambion) and cDNA synthesis using the GoScript RT Kit with oligo dT primers (Promega). The vector’s random barcode was amplified from the gDNA and cDNA via PCR with the Kapa HiFi HotStart ReadyMix and the primers TCGTCGGCAGCGTCAGATGTGTATAAGAGACAGTGCCTACGGACCGGCGC and GTCTCGTGGGCTCGGAGATGTGTATAAGAGACAGCTGGCCCTCGCAGACA. A second round of amplification was performed to add Illumina sequencing adapters with the same primers as for enhancer-barcode assignment. Three independent experiments were performed.

Following sequencing, random barcodes were extracted from gDNA and cDNA reads using Cutadapt (v.4.2) (*83*) with the options -g TGCCTACGGACCGGCGCGCCGATC…TGTCTGCGAGGGCCAGC -l 17 -m 17 --discard-untrimmed. Reads were filtered to retain only those with quality > 30 using fastp (v.0.23.2) (*84*). After assignments of the reads to enhancers based on the enhancer-barcode table, a count matrix with number of reads per enhancer is generated. Samples were processed using DESeq2 (v.1.34.0) (*85*), comparing the cDNA replicates versus the gDNA samples.

## In vivo validation of enhancer candidates

### Enhancer cloning in AAV vector

500 bp enhancer sequences with 15 bp adapters (GCCCTGCGTATGAGT and CTGAGCTCCCTAGGG in 5’ and 3’ respectively) were synthesized from Twist Bioscience. pSA358_pAAV-SCP1-Intron-eGFP-CS1 (Addgene #215513) was linearized via inverted PCR with the Kapa HiFi HotStart ReadyMix and the primers CTGAGCTCCCTAGGGTAC and ACTCATACGCAGGGCC, followed by DpnI digestion and gel extraction in a 0.8% agarose gel. The enhancer sequences and linearized plasmid were combined in an NEBuilder (New England Biolab) reaction with a vector to insert ratio of 1:2 for 45 min at 50°C. The reaction was then transformed into Stellar competent bacteria (Takara) and plated on LB agar plate with carbenicillin. Following maxiprep, the plasmid is sequenced with nanopore DNA sequencing (Oxford Nanopore Technologies).

### AAV production

HEK293T cells were cultured in DMEM high glucose + 10% FBS (Invitrogen). Transfection complex, containing PEI and OptiMEM (Invitrogen), was mixed and incubated with OptiMEM containing pΔF6 helper, pAAV-PHP.eB Rep/Cap (Addgene #103005) and pSA358_pAAV-SCP1-Intron-eGFP-CS1 with an enhancer for 20 min at room temperature (RT). Before adding the PEI:DNA complex, growth medium was replaced with DMEM + 1% FBS. 5 h post transfection DMEM + 10% FBS was added. 48 h post transfection, cells were harvested and centrifuged at 1,000 g at 4°C for 10 min. Cell pellets where lysed in lysis buffer (150 mM NaCl and 50 mM Tris-HCl pH 8.5 in endotoxin free H_2_O). Three freeze/thaw cycles were performed using a dry ice/ethanol mix, and a water bath at 37°C. Next, Supernatants were collected and Benzonase (Sigma) was added to a final concentration of 50 U/ml. After incubating 30 min at 37 °C, the lysates were centrifuged at 5,000 g for 20 min. Supernatant was filtered through a 0.45 μm filter (MillexHV) and carefully layered onto iodixanol gradients (15-25-40-60%) in 25 × 77 mm OptiSeal tubes (Beckman Coulter). Gradients were prepared using OptiPrep iodixanol (Sigma), 5 M NaCl, 5× PBS with 1 mM MgCl_2_ and 2.5 mM KCl (5× PBS-MK), and sterile H_2_O. The OptiSeal tubes containing gradient/viral load were centrifuged for 1 h 40 min at 50,000 RPM at 12°C in the Optima XE-100 Ultracentrifuge (Beckman Coulter). AAVs were collected with a 16 G needle from between the 40 and 60% layer, and 5 ml 1× PBS-MK was added. The diluted AAVs were desalted and concentrated by centrifugation at 5,000 g for 30 min at 20 °C in a pre-rinsed Amicon Ultra-15 filter (Millipore) in 1× PBS-MK. After 2 desalting rounds the concentrated AAV’s were washed with PBS containing 0.01% Pluronic F68 (ThermoFisher) by centrifugation at 5,000 g for 5 min at 20°C, aliquoted and stored at -80°C.

### Tail vein injection

C57B6/J mice of 6-8 weeks were used for this experiment. The mice were housed in individually ventilated cages in a room with a day-night cycle. Cages were enriched with housing material and extra cotton balls. Health of the mice was checked daily. Virus with 1x10^11^ multiplicity of infection was diluted up to a volume of 100 µl in saline and administered by tail vein injection using a 0.3ml syringe.

### Tissue sampling

21 days post-injection, the mice were sacrificed. The mice were flushed with 5 ml PBS and brains were dissected out. The brains were fixed overnight at 4°C in 4% PFA. After 3 washes with PBS, brains were incubated in 15% sucrose/PBS until saturated and next into 30% sucrose/PBS overnight at 4°C. 60 µm coronal sections were made using a microtome (Leica SM2010R) and stored in a cryoprotectant solution (30% glycerol, 30% ethylene glycol in PBS) at 4°C.

### Immunocytochemistry

Sections were washed with 0.1% triton X-100 in PBS and permeabilized in 0.25% triton X-100 in PBS for 30 min. Tissue was blocked with 2% BSA, 10% fetal bovine serum in 0.25% triton X-100-PBS for 1 h at RT. Primary antibody anti-GFP-Rabbit IgG (1/200; A-11122 Invitrogen) was incubated overnight at 4°C in 2% BSA, 10% goat serum in blocking solution. Secondary antibody AlexaFluor 488 Donkey anti-Rabbit IgG (H+L) (1/5,000; A-21206 Invitrogen) was incubated for 2.5 h at RT followed by 4’,6-diamidino-2-fenylindool (DAPI) (5 mM, Sigma, D9542) for 10 minutes at RT. In between steps, three wash steps of 10 minutes were performed with 0.1% triton-X in PBS. Brain sections were transferred to slides using ice cold 0.2% gelatin in 50 mM Tris-HCl pH7.5 and mounted in Mowiol-DABCO (25% Mowiol, Sigma, 81381, 2,5% DABCO, Sigma, D27802). Brain sections were imaged using a Nikon TiE A1R.

### Brain section image analysis and annotations

We used image processing software Fiji (https://imagej.net/software/fiji/) to filter the images such that the GFP signal was as clear as possible compared to the background (*86*). We did this by adjusting brightness thresholds, using the *Despeckle* noise filter. We also filtered the image with the *Remove background from image* option for the *KIAA1217 and ZNF804B enhancers.* Annotating the sections for the enhancer validation experiments was done by using the DAPI-stained nuclei to infer regions which could then be annotated manually and by usage of the Allen Brain Reference atlas (https://mouse.brain-map.org/static/atlas).

## Enhancer models

### Model training

The regions that were defined per topic were used as input data for deep learning models that can predict from a 500 bp sequence in which topic(s) it is accessible in a multi-class, multi-label manner, as has been done in earlier studies (*10, 11, 15*). The model’s architecture was updated and now consists of 5 convolutional layers, a dense layer and a classification layer using the Basset architecture as its foundation (*36*). We increased the number of filters per layer to 1024, 512, 512, 512, 512 respectively and added batch normalization and dropout to all convolutional layers and residual connections to the final two convolutional layers. Details of the model’s architecture for every dataset are provided in Supplementary Table S2-6. We used Adam Optimizer with a learning rate of 1e-3, decaying with a factor of 0.25 after no further increase in average validation auPR in 3 epochs. Early stopping was applied after 5 epochs without improvement. The batch size was 128. We trained such topic models for our five datasets: two mouse topic models, three human topic models and one chicken topic model, all with nearly the same architecture.

### Transfer learning to DARs

To obtain a per cell type classification output in our models, we applied transfer learning from the topic-based models to DAR-based models. For a cross-species model comparison, an output per cell type is more practical and beneficial compared to using topics: a cell type can have multiple topics, a topic can include multiple cell types and there is no guarantee that every cell type will have at least one topic. Therefore, for every cell type, we calculated a set of DARs (from pycisTopic (*4*) using the Wilcoxon rank sum test) and only retained regions which were above a certain log fold change threshold (a per model overview is provided in Supplementary Table S2-6). To remove potential promoter regions that may introduce a bias towards less cell type-specific TFBS motifs, we filtered out regions that were 500 bp up- or downstream of a transcription start site of any protein coding transcript. These DARs are then used to represent cell types.

For transfer learning to these DAR-based cell types, we loaded the weights from our topic model in the same architecture, froze all the convolutional layers, and replaced the dense and classification layer. We used a learning rate of 1e-4 to only train the weights of those two replaced layers while the convolutional layers remained frozen. Once the validation performance was saturated, we automatically selected the model with the highest validation auPR to further finetune. The convolutional layers were unfrozen and all parameters were now trained with a lower learning rate of 1e-6. Unlike during topic model training, there was no learning rate decay implemented. Early stopping after five epochs remained, the batch size was 32. The final model was selected based on the highest validation auPR score.

### Model validation

To validate the performance of our models, we split our training regions in a training, validation and test set with a 80-10-10% distribution. During training, the average validation auPR metric was used as the main measure of the model’s performance, and after training the weights at the epoch with the highest score were selected as the final model. This approach was used both in the topic model and DAR model training. An overview of the performance of all our models is provided in fig. S3.

### Nucleotide contribution scores

Nucleotide contribution scores were calculated as described in (*87*) using a neural network explainability tool called SHAP DeepExplainer (*34, 35*). We used 500 genomic regions to initialize the explainer.

### Pairwise sequence alignments

To visualize sequence changes in conserved regions, we computed pairwise sequence alignments with needle from the EMBOSS framework (*88*) using the default parameters (gapopen=10.0, gapextend=0.5, endopen=10.0, endextend=0.5, aformat3=pair).

### Detection of cell type-specific sequence patterns with TF-MoDISco

TF-MoDISco was used to infer characteristic sequence patterns separately for each cell type (*42*). We used DARs per cell type as input with a log-fold change of at least 1.5. We further removed promoter regions from the sets of DARs as described above and limited the input to regions that are in the center of pseudobulk ATAC-seq peaks. To identify regions that are in the center of peaks, we computed a peak imbalance score based on the pseudobulk ATAC-seq profile for each region per cell type. In particular, we computed the center of mass of the pseudobulk ATAC-seq profile per region and compared the average ATAC-seq signal up- and downstream of the center of mass. Regions with an imbalance score larger than 1.5 were removed. Before employing TF-MoDISco, we computed SHAP values for the filtered sets of DARs. For each cell type, this set of DARs was limited to regions with a prediction of at least 0.2. We then selected the top 5,000 or less DARs with the highest prediction score for the cell type for SHAP value computation. After computing SHAP values, we applied TF-MoDISco per type-specific DARs set with the following parameters: final_min_cluster_size=10, trim_to_window_size=15, initial_flank_to_add=5, final_flank_to_add=5, sliding_window_size=15, flank_size=5, target_seqlet_fdr=0.15, n_sample_null=5000, and max_seqlets_per_metacluster=50000.

To compare TF-MoDISco patterns with known TFBS motifs and against each other, we converted the patterns into position weight matrices (PWMs). For this purpose, we used TF-MoDISco’s internal function to convert contribution scores into a 2 bit information content scale and trimmed down the flanking regions on both ends using a threshold of 0.25. To compare the patterns to known TFBS motifs, we used tomtom from the MEME suite to compare the PWMs of the patterns against the cisTarget database using following parameters: no-ssc, oc=., min-overlap=5, dist=pearson, evalue, and thresh=10.0 (*4, 43*). Prior to running tomtom, we removed PWMs that had a width smaller than five. To remove PWMs with little information content, we computed the average Kullback-Leibler (KL) divergence per position against the genomic background frequencies and removed PWMs with a KL divergence smaller than 0.5. We employed tomtom in a similar fashion to compare all patterns of all cell types and data sets against each other. For comparing patterns against each other, we used tomtom with a threshold of 0.3.

For identifying sequence patterns that are shared between cell types, species, or data sets, we clustered TF-MoDISco patterns based on the e-value that we obtained from tomtom. For the graph-based clustering, we limited patterns to those that have a positive activity. First, we constructed a graph where verticesvertices are patterns and edges contain the -log_2_ e-value as weight describing the similarity of two patterns. We added a pseudocount of 10^-8^ to e-values before log-transformation. Edges were only added if the e-value was smaller than 0.01. In addition, we required the overlap between two matched patterns to be at least 70% of the size of the larger to avoid combining patterns to be matched if one of them was only a smaller sub-sequence of the larger pattern such as monomer and dimer patterns. We then performed Leiden clustering as implemented in the python *leidenalg* package using *ModularityVertexPartition* as partitioning method (*81*).

### Training and analyzing ChromBPNet models

ChromBPNet model training was performed as described in (*87*) using a pre-released version (*89*) from the ChromBPNet GitHub repository (https://github.com/kundajelab/chrombpnet/tree/v1.3-pre-release). We used pseudobulk MGL scATAC data obtained from pycisTopic from the DeepMouseBrain1 dataset (*4*) to make a scATAC profile track required for training. ChromBPNet takes 2114 bp regions as input. To score the 500 bp FIRE enhancer, we extended the sequence by 807 bp on both sides with its genomic flankings. The nucleotide contribution scores were obtained from the profile prediction head.

### Using the Enformer model

For calculating the FIRE nucleotide contribution scores from Enformer (*39*), we centered the enhancer in the 393 216 bp input sequence surrounded by its genomic flankings. To only measure the importance of the enhancer sequence without surrounding influences, we masked the genomic flanks before calculating the importance scores. We used mouse track 166 as MGL-specific ATAC track.

### Assigning transcription factors to potential binding sites

Annotating contribution score tracks with TFs at important motifs was done by first matching those motifs to a large TF-motif database (*4*), and by manually inspecting the expression of the candidate TFs in the cell type of interest to find the TF with the most specific expression.

## Enhancer-code based cell type similarity metrics

### Cross-species region predictions

From the DAR sets per cell type per dataset, we selected the top 100 regions (sorted on log fold change) after filtering (exclusion of promoters and non-peak-centered regions). We scored regions from cell types from one dataset with models trained different on datasets/species. From these scores per region we calculated the median prediction score per cell type. When consensus predictions were used from DeepMouseBrain and DeepHumanCortex models, we took the average of the prediction scores from both models for classes containing matching cell types. Also, for nucleotide contribution score Spearman correlation comparisons, the average of both models were used. For the DeepMouseBrain models, consensus cell type classes were AST, D1MSN, DG_GRC, EXC_CA1, EXC_L2_3_IT, EXC_L4_IT, EXC_L5_IT, EXC_L5_PT, EXC_L6_CT, EXC_L6_IT, EXC_NP, INH_PVALB, INH_SST, MGL, OL, OPC and VASC. For D2MSN, DG_NBL, EXC_CA3, EXC_CLA, EXC_L6b, EXC_PIR, INH_LAMP5, INH_STR, INH_VIP, IOL and RGL, predictions and contribution scores only came from DeepMouseBrain2 due to either an absence in the DeepMouseBrain1 dataset or too few cells (less than 350) leading to insufficient scATAC quality and worse prediction performance compared to DeepMouseBrain2. For the DeepHumanCortex models, consensus cell type classes were AST, EXC_L2_3_IT, EXC_L4_IT, EXC_L5_IT, EXC_L6_IT, INH_LAMP5, INH_PVALB, INH_SST, INH_VIP, MGL, OL and OPC. VASC predictions and contribution scores came from DeepHumanCortex1 due to absence of the cell type in the DeepHumanCortex2 dataset. For EXC_L6_CT and EXC_L6b, the same applies but in the opposite order.

### Nucleotide contribution score Spearman correlation

From the same set of cell-type specific regions per dataset, we computed the SHAP nucleotide contribution scores for every cell type (or output class) with the model that matches with the dataset and the other models. This results in vectors of 500 (sequence length) nucleotide contribution scores for all cell types per region and per model. We can use these nucleotide contribution scores from different models for the same region to compare how similar these are by using the Spearman correlation metric. Thus, for every region, we calculated the Spearman correlation between nucleotide contribution scores of all cell types of the model that matches with the dataset and the other models trained on different datasets or species. The correlation scores of the top 100 regions per cell type were aggregated by calculating the median to obtain a similarity score between all cell types on a nucleotide contribution level. To calculate Spearman correlations and their corresponding p-values, we used the *scipy.stats.spearmanr* function from Scipy (v1.5.3).

### Correlation of TF-MoDISco patterns

To compare the similarity of two cell types, we computed the Spearman correlation over the number of instances of TF-MoDISco pattern clusters (see above). In particular, for each deep learning model and for each cell type, we collected the number of instances of TF-MoDISco patterns (called seqlets) of each cluster and normalized the numbers by the number of cell-type specific DARs that contained any seqlet. By this way, we obtained vectors of the average occurrence of clustered TF-MoDISco pattern for each cell type and for each deep learning model. We then computed Spearman correlation coefficients between these vectors for each cell type of each model and each species. Here, we limited the clustered patterns to those that are detected in all three species. To cluster cell types based on the correlation of clustered TF-MoDISco patterns, we converted the pairwise Spearman correlation coefficients (SCCs) into a dissimilarity matrix (dissimilarity=1-SCC) and performed average linkage hierarchical clustering using the python package *scipy.cluster.hierarchy* (v1.5.3).

## Acknowledgements

We thank Eve Seuntjens and Veerle Darras for helping with the chicken brain dissections; Pierre Vanderhaeghen for advice; Bastienne Zaremba, Fernando Garcia-Moreno and Henrik Kaessmann for helpful comments on the manuscript. We further thank Kristofer Davie and the VIB Bioimaging Core for help with Stereo-seq data and the Vlaams Supercomputer Center for computing infrastructure. We also thank Carles Calatayud Aristoy, Blanca Lorente Echeverrìa, Gabriele Marcassa, Dan Dascenco, Danie Daaboul and Lars Borm for their help and advice on virus productions and injections, imaging, and image analysis.

## Authors contributions

N.H. and N.K are listed as co-first authors because they contributed equally to the manuscript. Conceptualization: N.H. and S.A. Experiments and sample preparation: D.M., R.V., S.D., E.L., R.M., V.C. and S.P. Computational analysis: N.H., N.K., D.A., I.S. and C.B.G. Data collection, processing, and curation: N.H., G.H., N.K., and C.B.G. Methodology: N.H., N.K. and S.A. Software implementation and testing: N.K., N.H., D.A. and I.S. Resources: S.P., L.L., J.D.W, and S.A. Visualization: N.H., N.K., D.M. and D.A. Writing - original draft: N.H., N.K., and S.A. Writing - review and editing: N.H., N.K., D.M., D.A., C.B.G., I.S., and S.A. Funding acquisition: S.A., N.H. and N.K.

## Data availability

The weights of the DeepMouseBrain, DeepHumanCortex and DeepChickenBrain models and code for making predictions, getting contribution scores and calculating cell-type contribution score correlation is available at https://github.com/aertslab/DeepBrain. A UCSC track hub of the scATAC profiles of all datasets is available at https://ucsctracks.aertslab.org/papers/brain_evo/hub.txt. Additional data is available on Zenodo (https://zenodo.org/records/10868679), containing all the TF-MoDISco pattern results, the code for the MPRA analysis and an extra storage location for the model weights. Raw and processed data of the chicken telencephalon 10x Single Cell Multiome ATAC + Gene expression, chicken telencephalon Stereo-seq, and MPRA in BV2 cell lines that was generated for this study are available at the NCBI Gene Expression Omnibus (GEO; https://www.ncbi.nlm.nih.gov/geo/): GSE262320 (MPRA), GSE262321 (single cell multiome), and GSE262322 (Stereo-seq).

## Supplementary information

- Supplementary_Information.pdf: Figure S1-S29, Tables S1-6 (captions)
- Supplementary_Tables.xlsx: Tables S1-6

