## Supplementary Information for "Enhancer-driven cell type comparison reveals similarities between the mammalian and bird pallium"

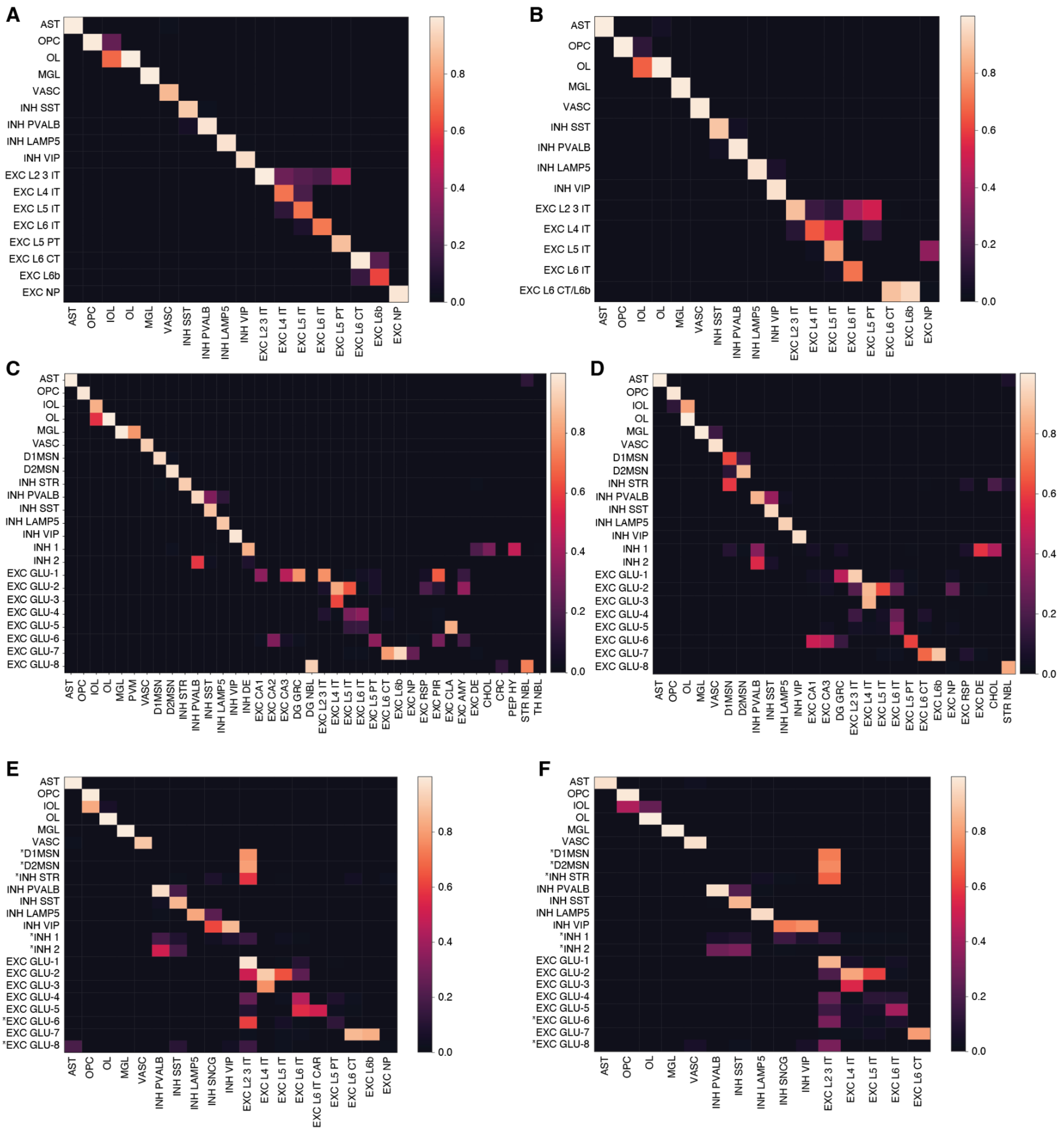


**Fig. S1. Transcriptome comparison using SAMap.** Heatmaps are showing cell type mapping scores computed with SAMap between (**A**) human motor cortex (*24*) and mouse neocortical cell types (*30*), (**B**) human prefrontal cortex (*26*) and and mouse neocortical cell types (*30*), (**C**) chicken telencephalon and mouse brain cell types (*30*), (**D**) chicken telencephalon and mouse brain cell types (*4*), (**E**) chicken telencephalon and human motor cortex (*24*) cell types, (**F**) chicken telencephalon and human prefrontal cortex (*26*) cell types. Asterisks indicate chicken telencephalon cell types that likely do not have neocortical correspondences based on mouse transcriptome comparisons and thus exhibit unreliable mappings in human neocortical data sets.

**A**

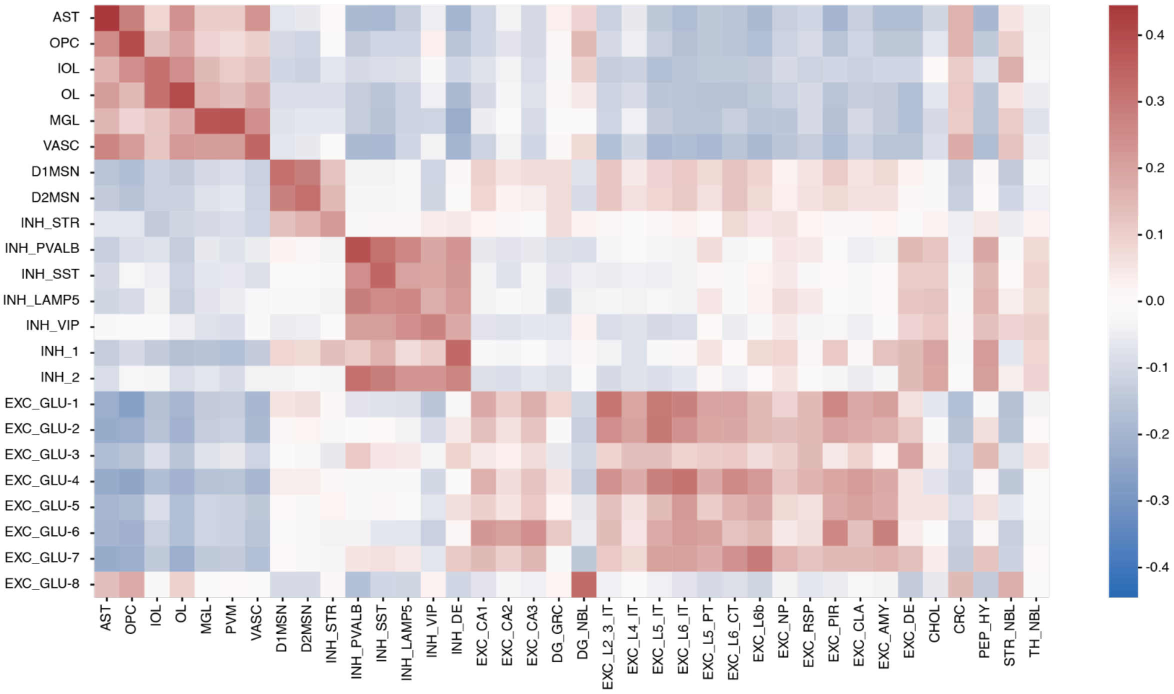


**B**

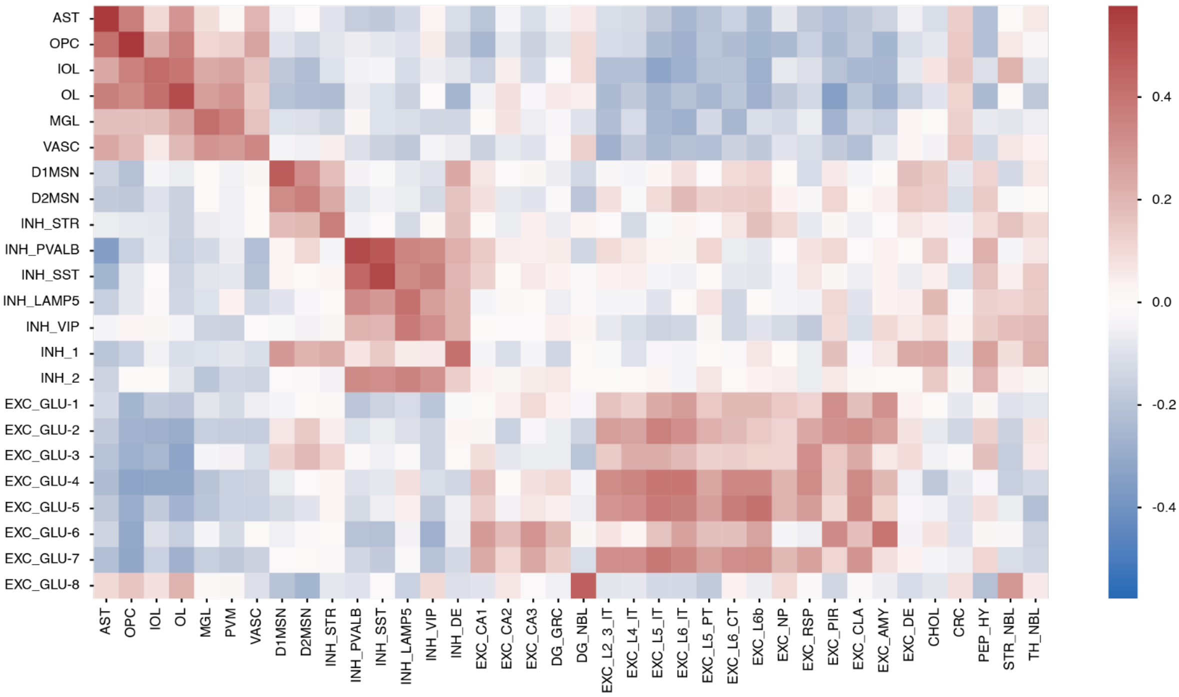


**Fig. S2. Gene expression correlation of 1:1 orthologs.** Heatmaps indicate the Spearman correlation between z-scores of mean gene expression values of (**A**) 1:1 chicken and mouse orthologs between chicken telencephalon and mouse brain cell types (*30*), and (**B**) 1:1 TF chicken and mouse orthologs between chicken telencephalon and mouse brain cell types (*30*). We limited the analysis to genes that were expressed in at least 25% of any cell-type and which mean log-normalized expression had a z-score of at least 1.65 for any cell type.

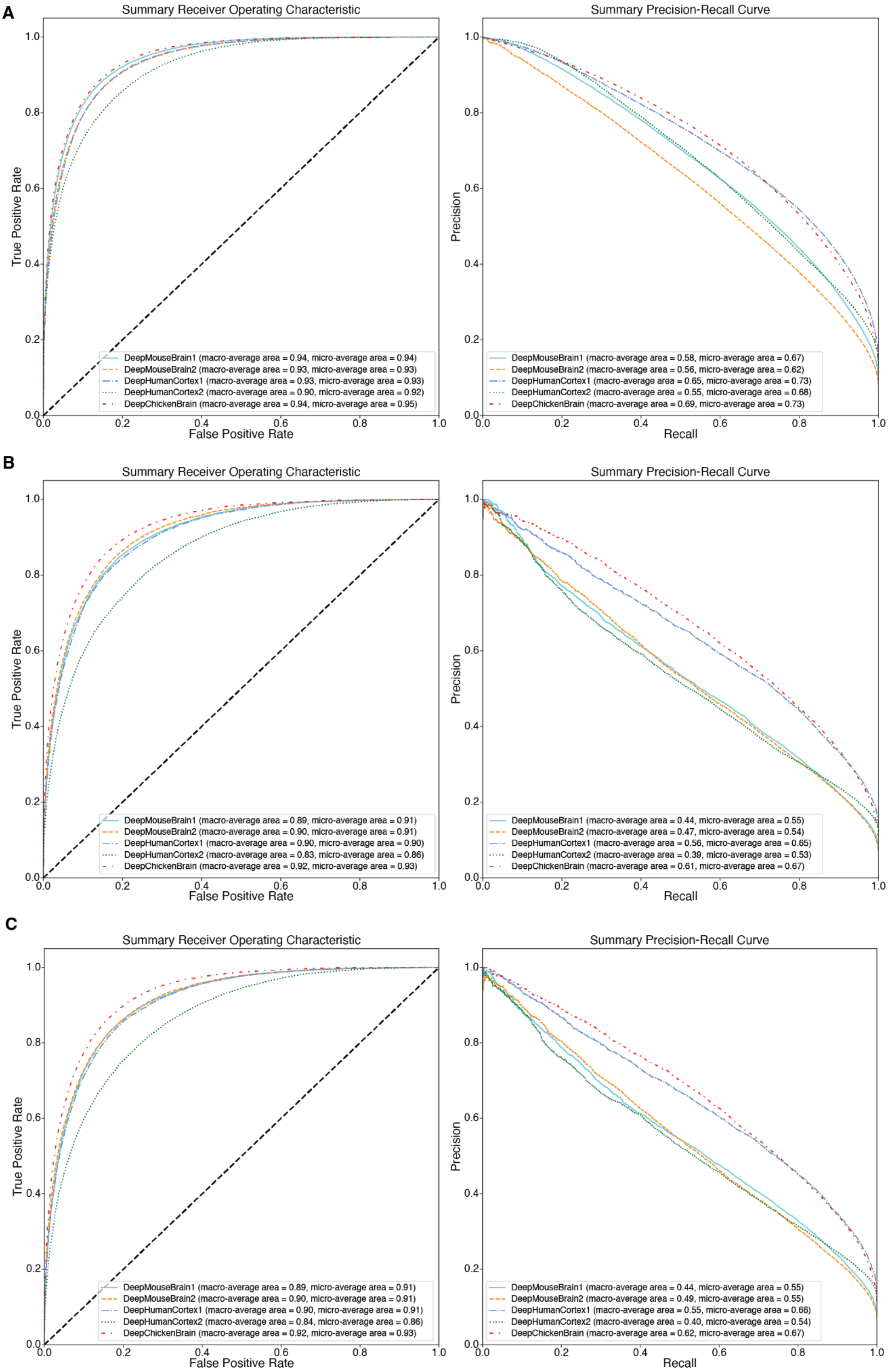

A
Summary Receiver Operating Characteristic
True Positive Rate
False Positive Rate
DeepMouseBrain1 (macro-average area = 0.94, micro-average area = 0.94)
DeepMouseBrain2 (macro-average area = 0.93, micro-average area = 0.93)
DeepHumanCortex1 (macro-average area = 0.93, micro-average area = 0.93)
DeepHumanCortex2 (macro-average area = 0.90, micro-average area = 0.92)
DeepChickenBrain (macro-average area = 0.94, micro-average area = 0.95)
Summary Precision-Recall Curve
Precision
Recall
DeepMouseBrain1 (macro-average area = 0.58, micro-average area = 0.67)
DeepMouseBrain2 (macro-average area = 0.56, micro-average area = 0.62)
DeepHumanCortex1 (macro-average area = 0.65, micro-average area = 0.73)
DeepHumanCortex2 (macro-average area = 0.55, micro-average area = 0.68)
DeepChickenBrain (macro-average area = 0.69, micro-average area = 0.73)
B
Summary Receiver Operating Characteristic
True Positive Rate
False Positive Rate
DeepMouseBrain1 (macro-average area = 0.89, micro-average area = 0.91)
DeepMouseBrain2 (macro-average area = 0.90, micro-average area = 0.91)
DeepHumanCortex1 (macro-average area = 0.90, micro-average area = 0.90)
DeepHumanCortex2 (macro-average area = 0.83, micro-average area = 0.86)
DeepChickenBrain (macro-average area = 0.92, micro-average area = 0.93)
Summary Precision-Recall Curve
Precision
Recall
DeepMouseBrain1 (macro-average area = 0.44, micro-average area = 0.55)
DeepMouseBrain2 (macro-average area = 0.47, micro-average area = 0.54)
DeepHumanCortex1 (macro-average area = 0.56, micro-average area = 0.65)
DeepHumanCortex2 (macro-average area = 0.39, micro-average area = 0.53)
DeepChickenBrain (macro-average area = 0.61, micro-average area = 0.67)
C
Summary Receiver Operating Characteristic
True Positive Rate
False Positive Rate
DeepMouseBrain1 (macro-average area = 0.89, micro-average area = 0.91)
DeepMouseBrain2 (macro-average area = 0.90, micro-average area = 0.91)
DeepHumanCortex1 (macro-average area = 0.90, micro-average area = 0.91)
DeepHumanCortex2 (macro-average area = 0.84, micro-average area = 0.86)
DeepChickenBrain (macro-average area = 0.92, micro-average area = 0.93)
Summary Precision-Recall Curve
Precision
Recall
DeepMouseBrain1 (macro-average area = 0.44, micro-average area = 0.55)
DeepMouseBrain2 (macro-average area = 0.49, micro-average area = 0.55)
DeepHumanCortex1 (macro-average area = 0.55, micro-average area = 0.66)
DeepHumanCortex2 (macro-average area = 0.40, micro-average area = 0.54)
DeepChickenBrain (macro-average area = 0.62, micro-average area = 0.67)

**Fig. S3. DeepBrain models ROC and PR curves on training, validation and test splits.**
(**A**) ROC and PR curves for all models scored on training set regions. Macro-averages are the average performances on all classes, micro-averages the average performance on all samples separately. (**B**) ROC and PR curves for all models scored on validation set regions. (**C**) ROC and PR curves for all models scored on test set regions.

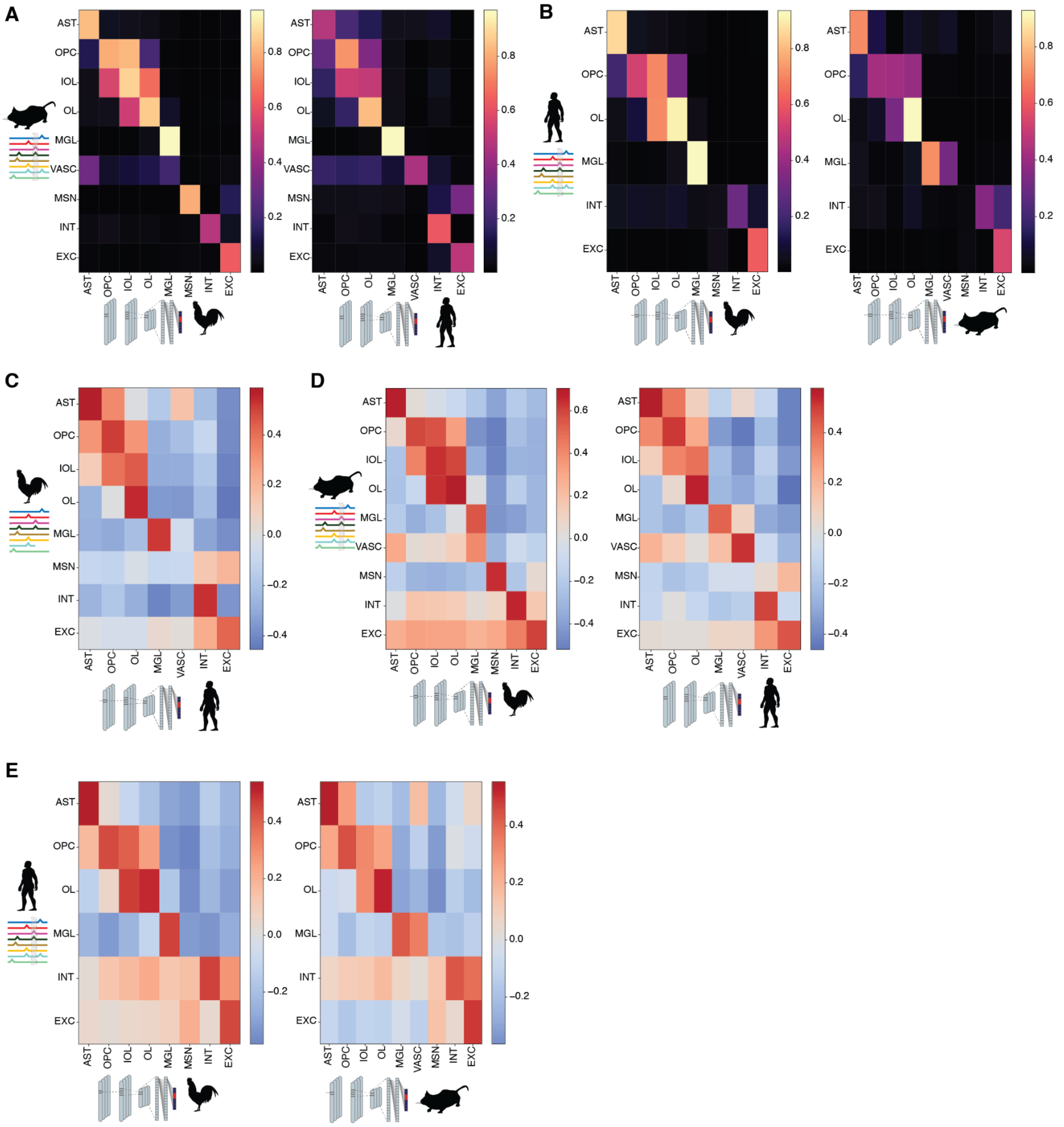


**Fig. S4. Grouped cross species predictions and nucleotide contribution Spearman correlation scores.** (**A**) Cross-species model predictions from DeepChickenBrain and DeepHumanCortex1&2 models on the top 100 DARs per mouse cell type from the DeepMouseBrain2 dataset. The median of the predictions per region is shown. Human predictions are the consensus of the DeepHumanCortex models. Excitatory neuron, MSN and interneuron cell types are grouped. (**B**) Same analysis for the top human DARs per cell type from the DeepHumanCortex2 dataset scored by DeepChickenBrain (left) and DeepMouseBrain1&2 (right). (**C**) Cross-species nucleotide contribution scores Spearman correlation between the DeepHumanCortex models and DeepChickenBrain for the top 100 DARs per chicken cell type. The median of the correlation per region is shown. The correlations are the consensus of two comparisons between DeepHumanCortex1&2 and DeepChickenBrain. Excitatory neuron, MSN and interneuron cell types are grouped. (**D**) Identical configuration as (B) but now for nucleotide contribution scores Spearman correlation between DeepChickenBrain and DeepMouseBrain2 (left) and DeepHumanCortex1&2 and DeepMouseBrain2 (right). (**E**) Identical configuration as (C) but now for nucleotide contribution scores Spearman correlation between DeepChickenBrain and DeepHumanCortex2 (left) and DeepHumanCortex1&2 and DeepHumanCortex2 (right).

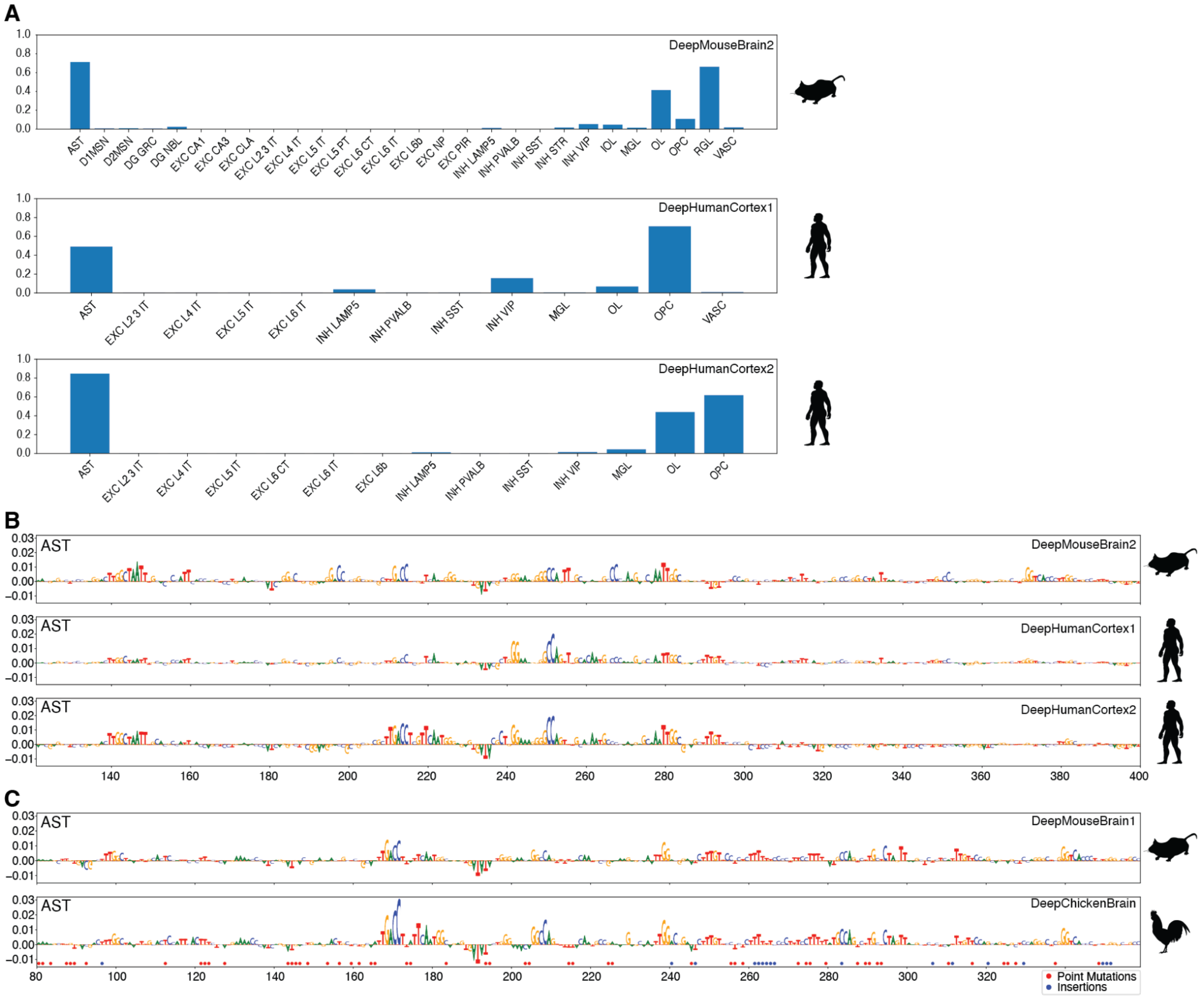


**Fig. S5. Model scoring on the identified *Prdm16a* region and its chicken homolog.**
(**A**) DeepMouseBrain2 and DeepHumanCortex1&2 predictions on the *Prdm16a* region. (**B**) DeepMouseBrain2 and DeepHumanCortex1&2 nucleotide contribution scores for predicting the *Prdm16a* region as astrocyte. (**C**) DeepMouseBrain1 and DeepChickenBrain nucleotide contribution scores for predicting the chicken *PRDM16a r*egion (galGal6 chr21:1148446-1148946) as astrocyte. Point mutations and insertions in the chicken sequence when aligned to the mouse sequence are indicated in red and blue respectively.

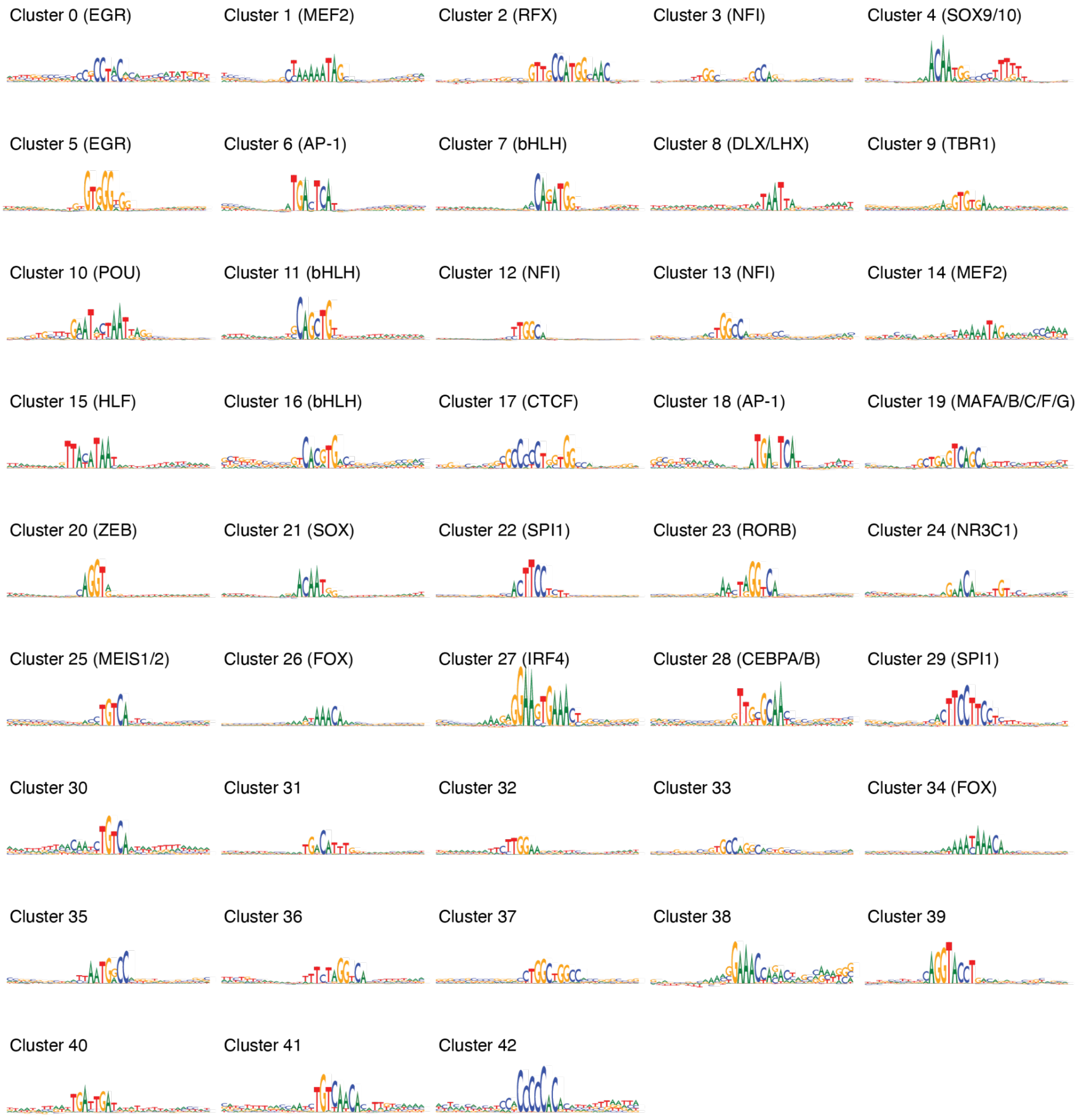


**Fig. S6. Detected clusters of TF-MoDISco motifs.** TF-MoDISco patterns were clustered across species and cell types, then clusters of motifs were identified using Leiden modularity-based clustering (Methods). Depicted are the patterns with the highest PageRank and corresponding TF families, gene families or gene names if it could be inferred by similarity to known motifs in the cisTarget database and expression in the analyzed brain data sets.

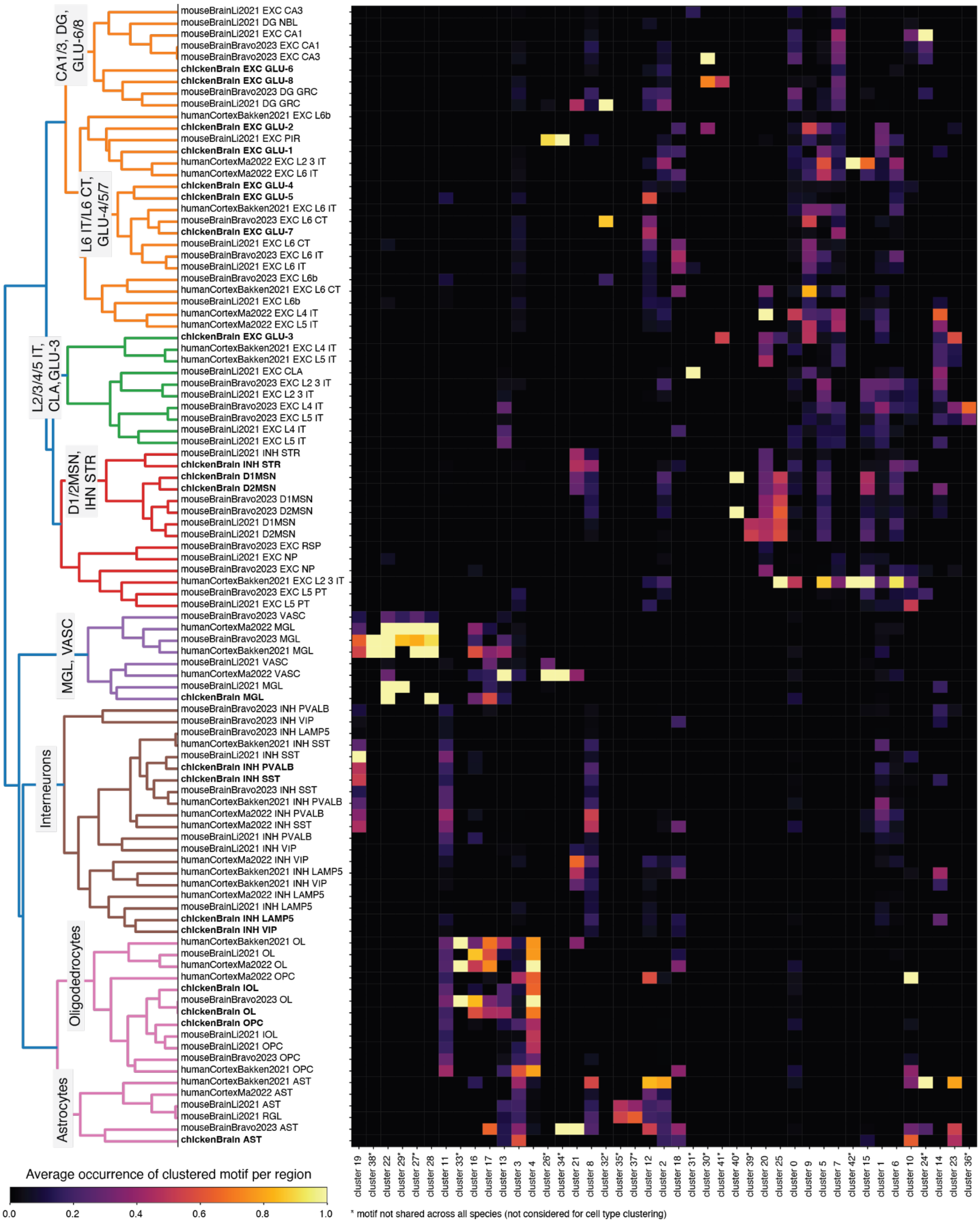


**Fig. S7. Cell types clustered by TF-MoDISco motif clusters.** Cell types were clustered across species based on the correlation of the average occurrence of motifs per region (Methods). For this purpose, the Spearman correlation between cell types was computed and converted into a dissimilarity matrix. Afterwards average linkage clustering was performed. For the comparison, we excluded patterns which were not found in all three species (human, mouse, and chicken); indicated by asterisks. The heatmap shows the average occurrence of motifs per TF-MoDISco input regions.

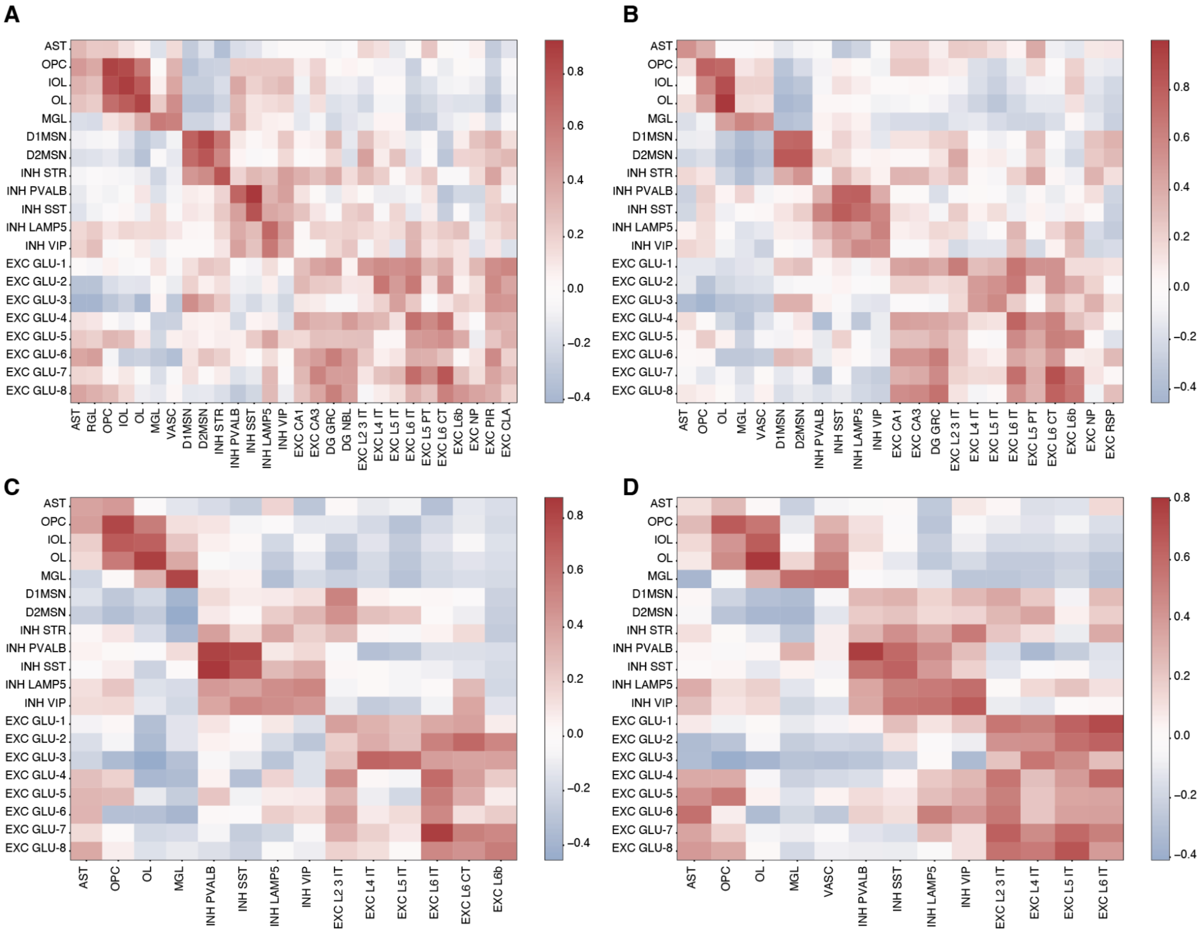


**Fig. S8. Correlation over TF-MoDISco motif clusters.** Heatmaps depict the Spearman correlation over the average occurrence of TF-MoDISco motif clusters per region (Methods) between the (**A**) chicken telencephalon and mouse brain (*25*) cell types, (**B**) chicken telencephalon and mouse brain cell types (*4*), (**C**) chicken telencephalon and human motor cortex (*24*) cell types, and (**D**) chicken telencephalon and human prefrontal cortex (*26*) cell types.

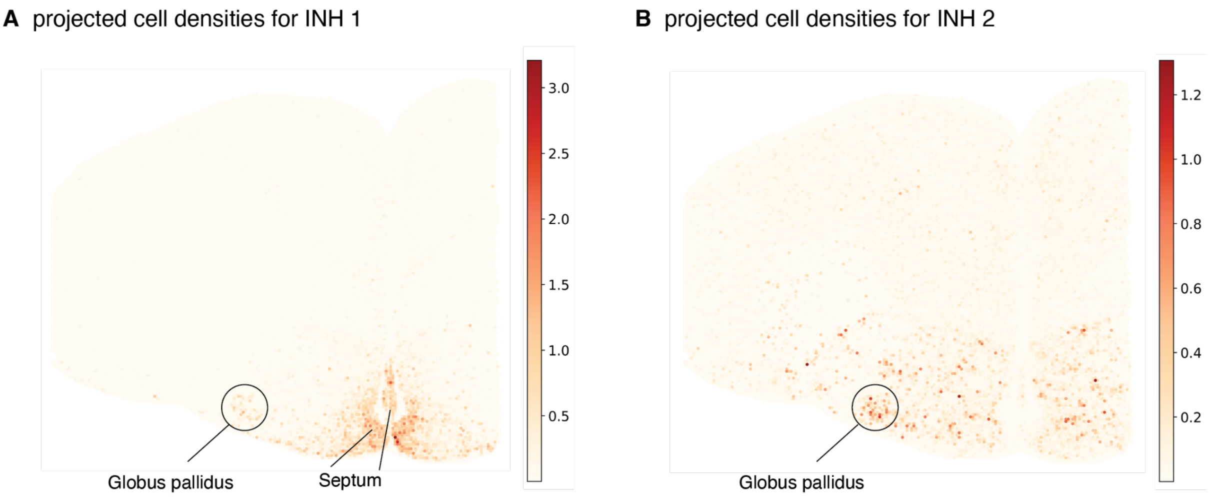


**Fig. S9: Localization of INH 1 and INH 2 clusters in chicken Stereo-seq data.** Depicted are cell densities obtained from cell2location (*30*) by projecting the chicken telencephalon single cell clusters to Stereo-seq data for cluster INH 1 **(A)** and INH 2 **(B)**. Expected anatomical structures that overlap projected labels are indicated.

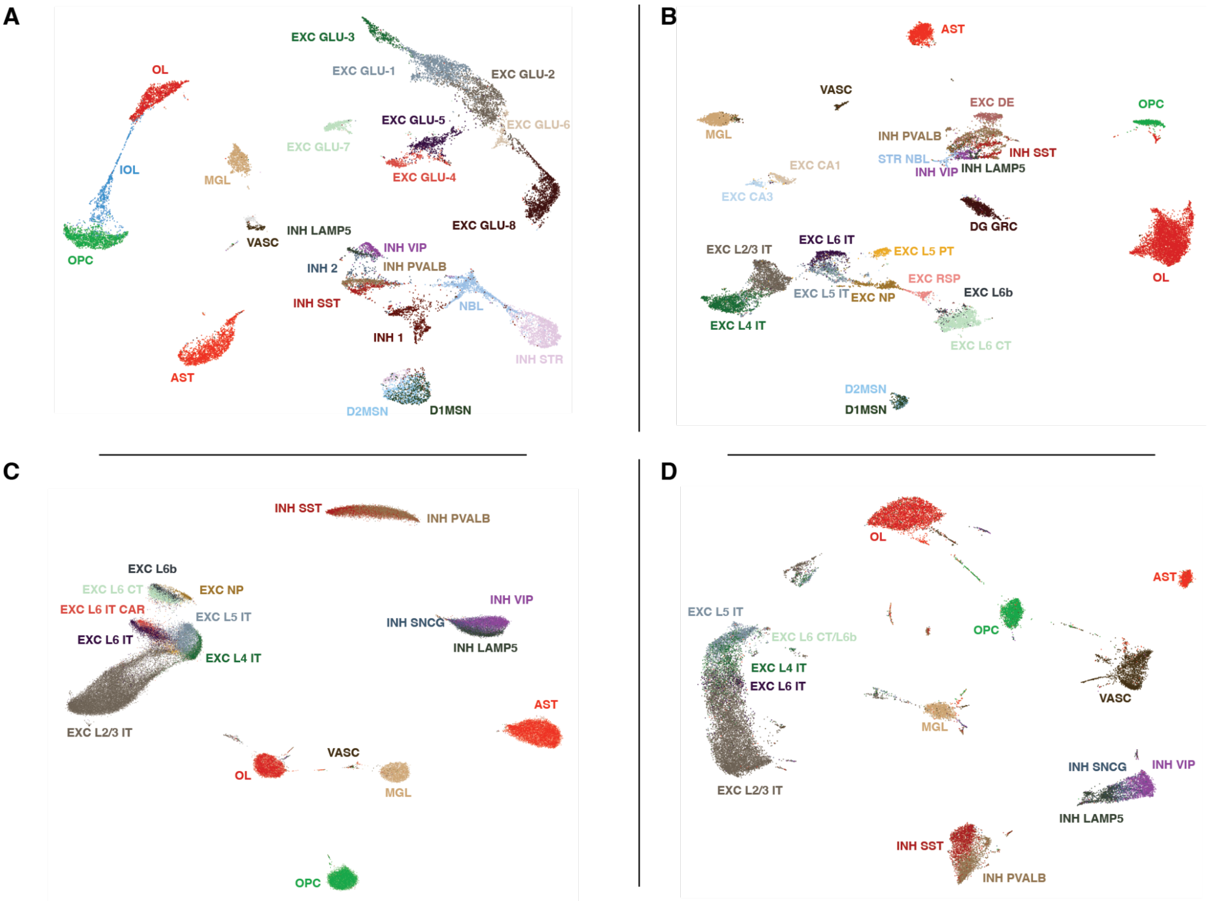


**Fig. S10. UMAPs of ATAC-seq modalities.** UMAP of (**A**) chicken telencephalon, (**B**) mouse brain (*4*), (**C**) human motor cortex (*24*), and (**D**) human prefrontal cortex cells (*26*).

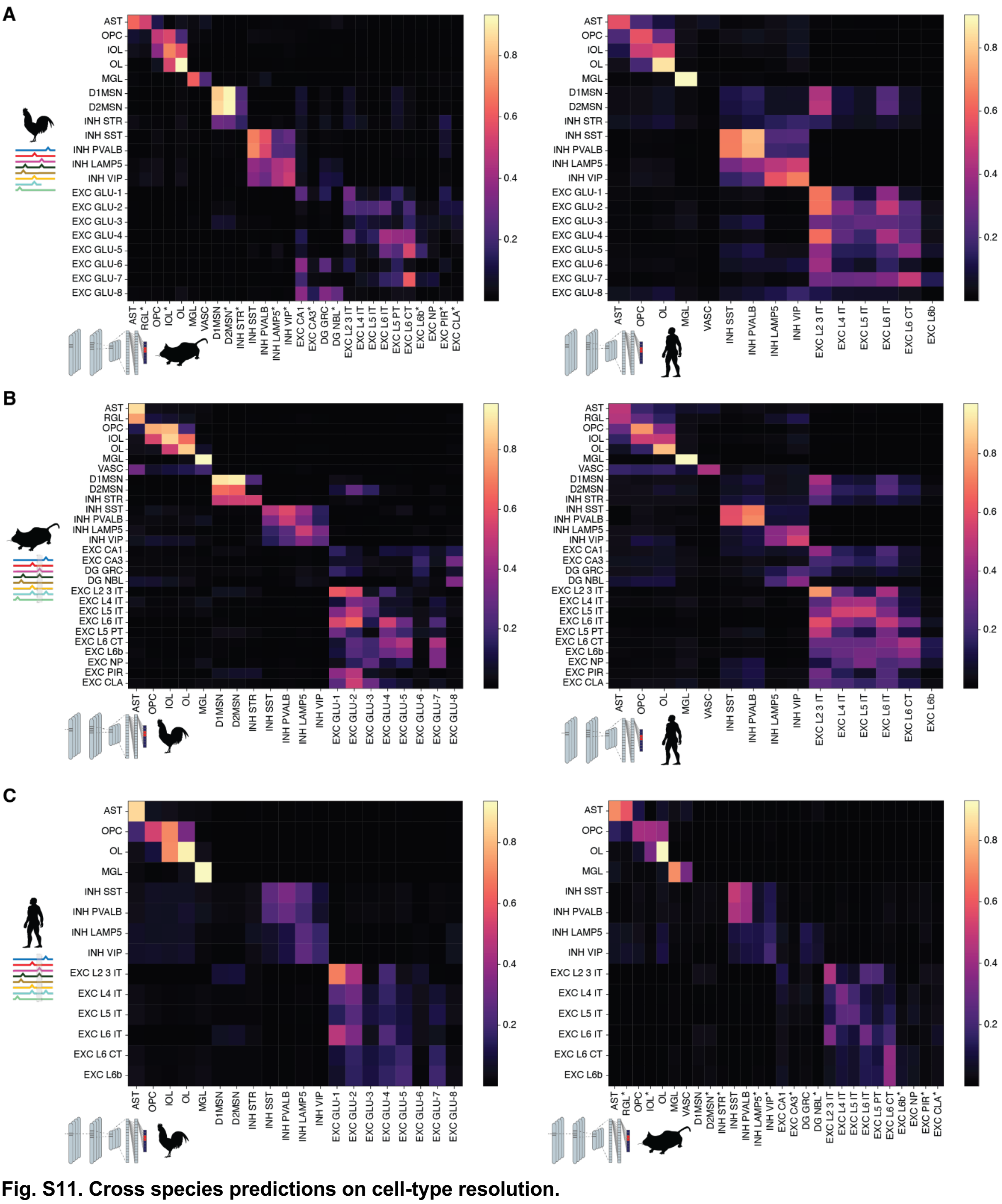

**Fig. S11. Cross species predictions on cell-type resolution.**
(**A**) Cross-species model predictions from the DeepMouseBrain (left) and DeepHumanCortex (right) models on the top 100 DARs per chicken cell type. The median of the predictions per region is shown. The prediction scores are the consensus of two comparisons between DeepMouseBrain1&2 and DeepChickenBrain (left) and DeepHumanCortex1&2 and DeepChickenBrain. (**B**) Same comparison for mouse regions from the DeepMouseBrain2 dataset showing DeepChickenBrain (left) and DeepHumanCortex1&2 (left) prediction scores. (**C**) Same comparison for mouse regions from the DeepHumanCortex2 dataset showing DeepChickenBrain (left) and DeepMouseBrain1&2 (left) prediction scores. In (A) and (C), mouse cell types annotated with an asterisk only contain predictions from DeepMouseBrain2.

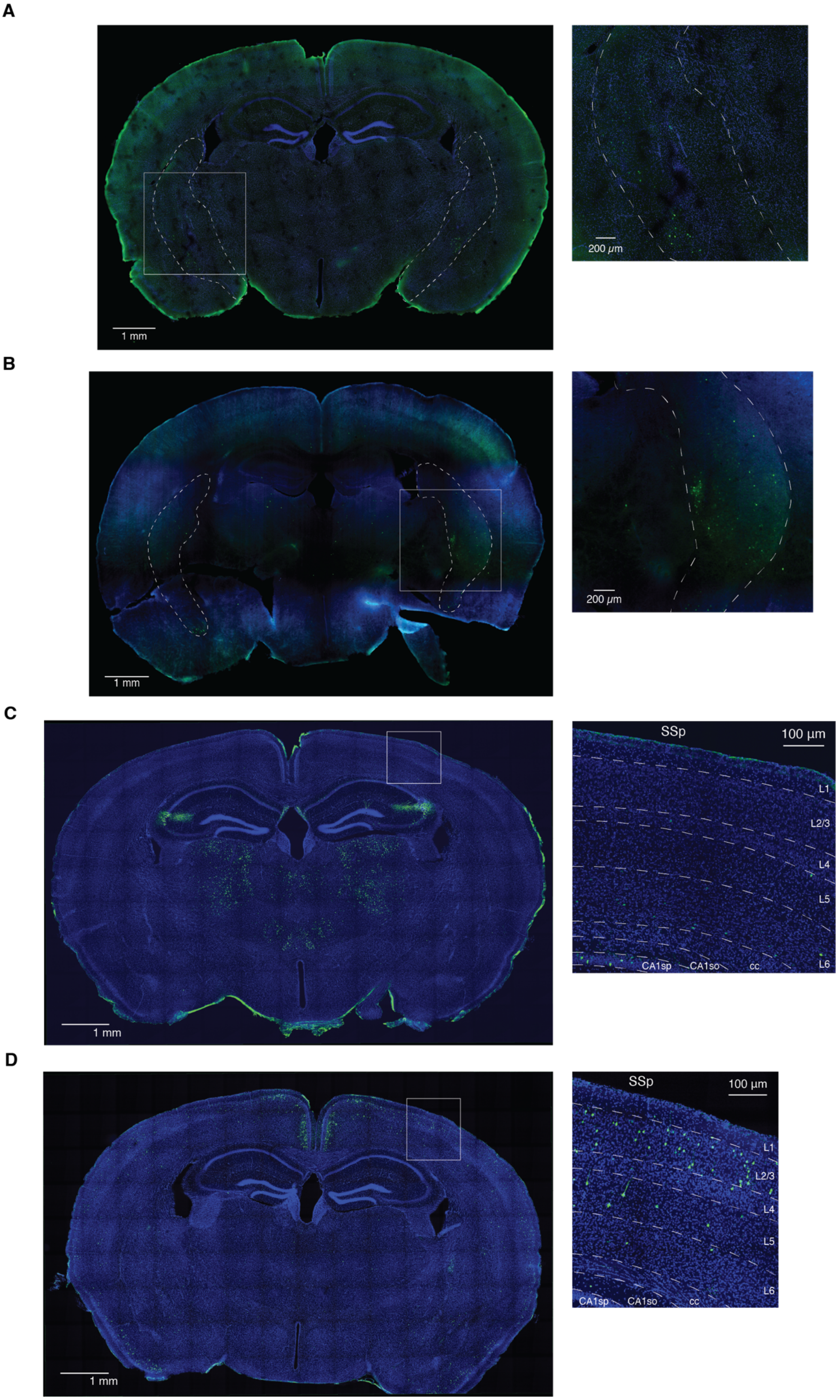
A
1 mm
200 μm
B
1 mm
200 μm
C
1 mm
SSp
100 μm
L1
L2/3
L4
L5
L6
CA1sp
CA1so
cc
D
1 mm
SSp
100 μm
L1
L2/3
L4
L5
L6
CA1sp
CA1so
cc

**Fig. S12. Whole brain images of enhancer-reporter assays.**
Whole mouse brain images (left) and zoomed in images (right) of the highlighted areas (white square) of DAPI-stained nuclei (blue) and enhancer-driven GFP expression (green) from the mouse *Foxp2* enhancer **(A)**, the chicken *FOXP2* enhancer **(B)**, the chicken *KIAA1217* enhancer **(C)** and the chicken *ZNF804B* enhancer **(D)**.

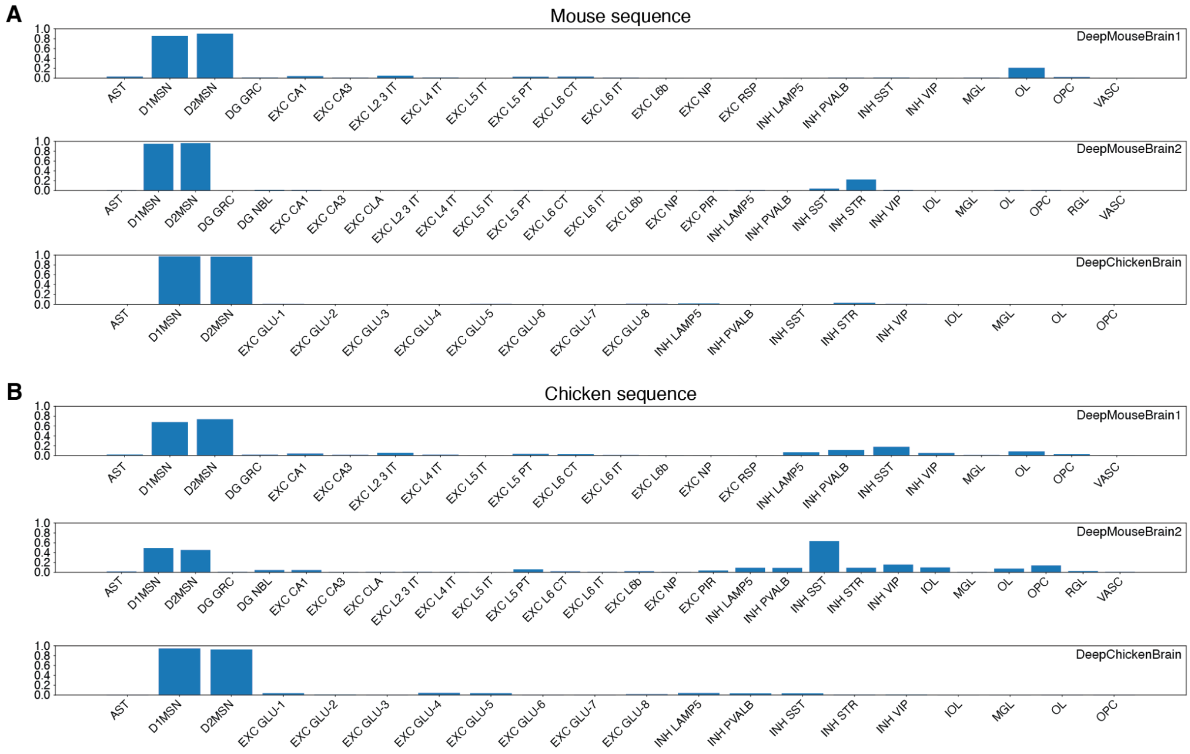


**Fig. S13. Model scoring on the identified *Foxp2* region and its chicken homolog.**
(**A**) DeepMouseBrain1&2 and DeepChickenBrain predictions on the mouse *Foxp2* region (reverse complemented). (**B**) DeepMouseBrain1&2 and DeepChickenBrain predictions on the chicken *FOXP2* region (reverse complemented).

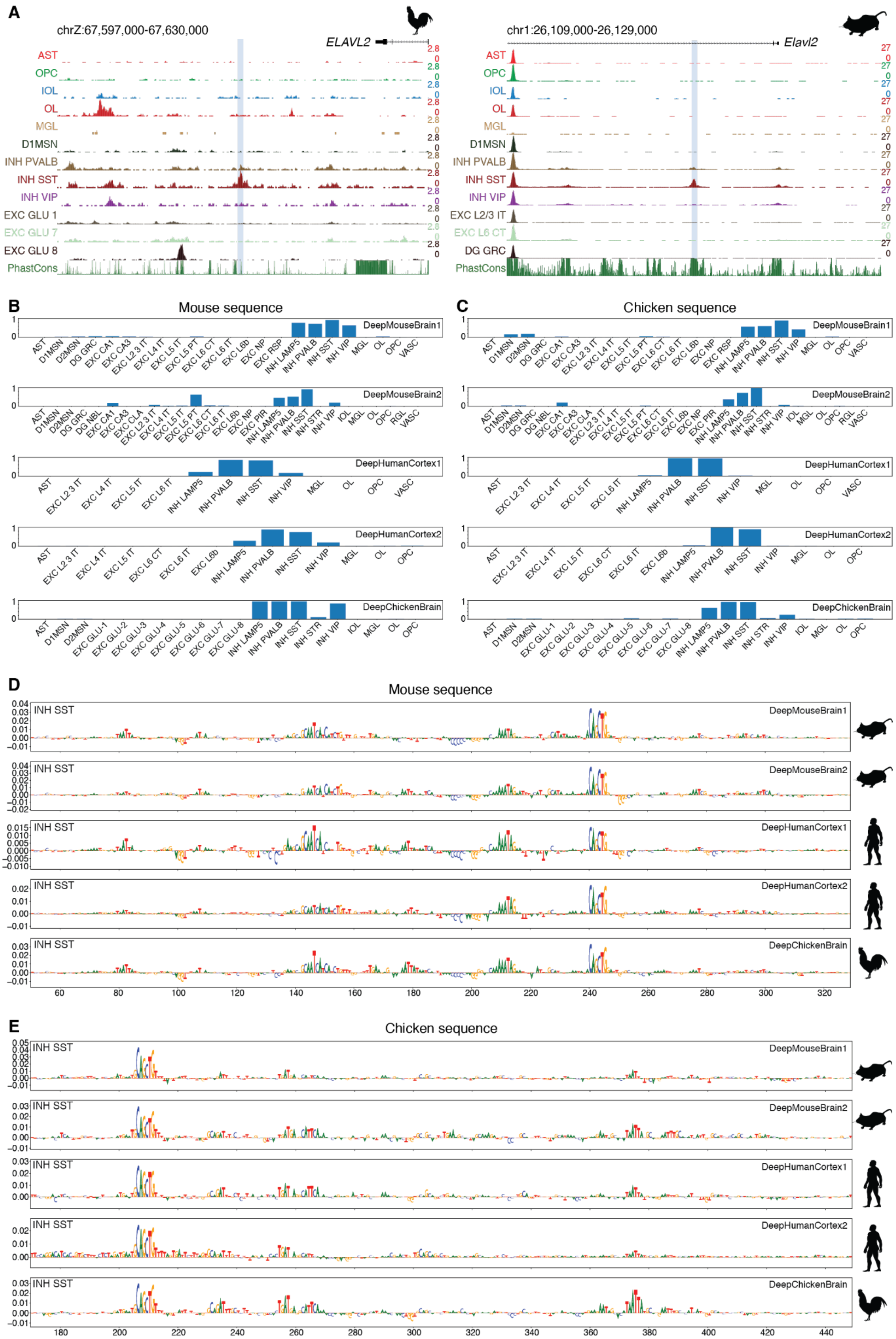

A
chrZ:67,597,000-67,630,000
ELAVL2
chr1:26,109,000-26,129,000
Elavl2
AST
OPC
IOL
OL
MGL
D1MSN
INH PVALB
INH SST
INH VIP
EXC GLU 1
EXC GLU 7
EXC GLU 8
EXC L2/3 IT
EXC L6 CT
DG GRC
PhastCons
B
Mouse sequence
C
Chicken sequence
DeepMouseBrain1
DeepMouseBrain2
DeepHumanCortex1
DeepHumanCortex2
DeepChickenBrain
D
Mouse sequence
E
Chicken sequence
INH SST

**Fig. S14. Conservation of SST enhancer code for a mouse and chicken candidate enhancer region near *Elavl2*/*ELAVL2.***

(**A**) scATAC tracks of the chicken (left) and mouse (right) *ELAVL2* enhancer candidates (galGal6 chrZ:67613064-67613564 and mm10 chr4:91392313-91392813) showing SST specificity. (**B**) DeepMouseBrain1&2, DeepHumanCortex1&2 and DeepChickenBrain predictions on the mouse *Elavl2* region. (**C)** DeepMouseBrain1&2, DeepHumanCortex1&2 and DeepChickenBrain predictions on the chicken Elavl2 region. (**D**) DeepMouseBrain1&2, DeepHumanCortex1&2 and DeepChickenBrain contribution scores for their SST predictions on the mouse *Elavl2* region. (**E**) DeepMouseBrain1&2, DeepHumanCortex1&2 and DeepChickenBrain contribution scores for their SST predictions on the chicken *ELAVL2* region.

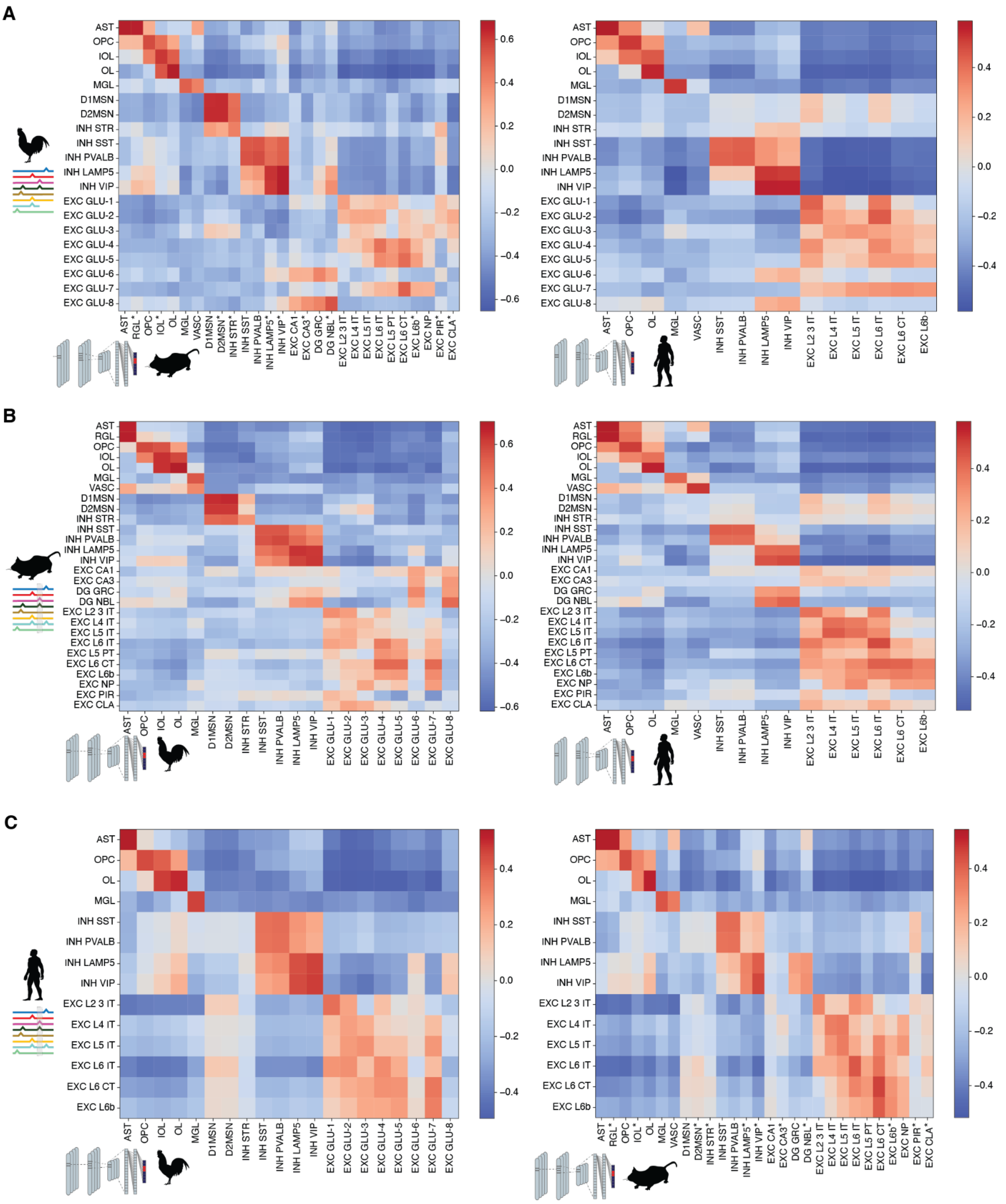


**Fig. S15. Cross species nucleotide contribution Spearman correlation on cell-type resolution.**
(**A**) Cross-species nucleotide contribution scores Spearman correlation between the DeepMouseBrain (left) and DeepHumanCortex (right) models and DeepChickenBrain for the top 100 DARs of all cell types. The median of the correlation per region is shown. The correlations are the consensus of two comparisons between DeepMouseBrain1&2 and DeepChickenBrain (left) and DeepHumanCortex1&2 and DeepChickenBrain. (**B**) Same comparison for mouse regions from the DeepMouseBrain2 dataset showing nucleotide contribution correlation between DeepChickenBrain and DeepMouseBrain2 (left) and DeepHumanCortex1&2 and DeepMouseBrain2 (right). (**C**) Same comparison for human regions from the DeepHumanCortex2 dataset showing nucleotide contribution correlation between DeepChickenBrain and DeepHumanCortex2 (left) and DeepMouseBrain1&2 and DeepHumanCortex2 (right). In (A) and (C), mouse cell types annotated with an asterisk only contain correlation scores from DeepMouseBrain2.

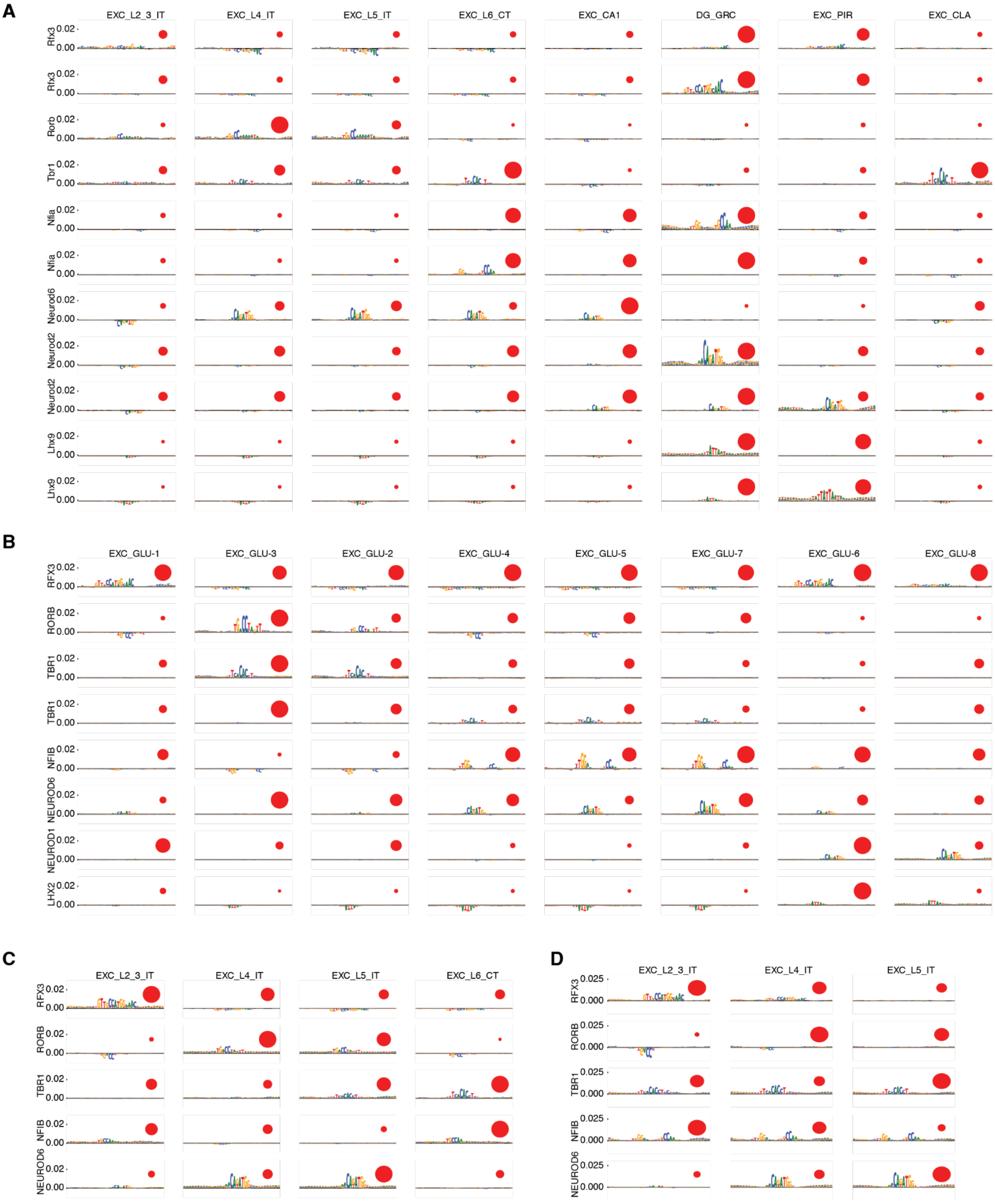


**Fig. S16. Nucleotide contribution scores of selected TF-MoDISco patterns for excitatory neurons.** TF-MoDISco was used to detect patterns across excitatory neurons based on DARs of the cell types. Depicted are nucleotide contribution scores for selected patterns of (**A**) mouse brain (*25*), (**B**) chicken telencephalon, (**C**) human motor cortex (*24*), and (**D**) human prefrontal cortex (*26*) cell types. The size of red circles indicates the mean log-normalized expression of TFs that may correspond to the shown patterns. Similar patterns are shown if different instances of the patterns (seqlets) result in different activity patterns across cell types, e.g. for Nfia-like and Neurod2-like patterns.

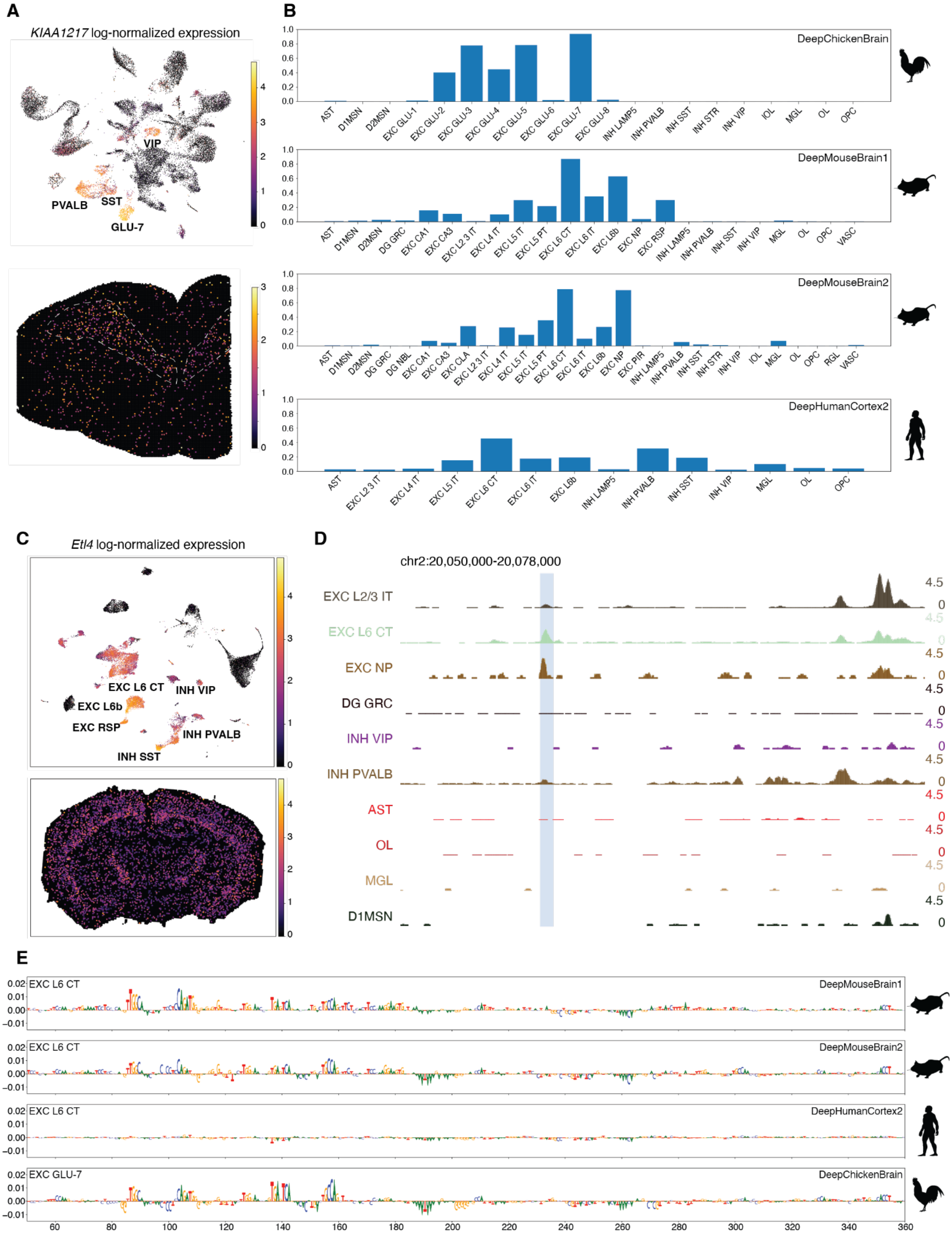


**Fig. S17. Conservation of GLU-7 and L6 CT enhancer code for a mouse and chicken candidate enhancer region near *KIAA1217*/*Etl4*.**

**(A)** scRNA-seq UMAP showing *KIAA1217* expression in chicken telencephalon single cell data (top), and its spatial expression in spatially resolved transcriptomics (Stereo-seq) chicken data (bottom). **(B)** DeepMouseBrain1&2, DeepHumanCortex1&2 and DeepChickenBrain predictions on the chicken *KIAA1217* region. **(C)** scRNA-seq UMAP showing *Etl4* expression in mouse telencephalon (top), and its spatial expression in spatially resolved transcriptomics (Stereo-seq) mouse data (bottom). (**D**) scATAC tracks of the mouse *Etl4* enhancer candidate (mm10 chr2:20063609-20064109). (**E**) DeepMouseBrain1&2, DeepHumanCortex2 and DeepChickenBrain contribution scores for their EXC L6 CT (DeepMouseBrain1&2 and DeepHumanCortex2) and GLU-7 (DeepChickenBrain) predictions on the mouse *Etl4* region.

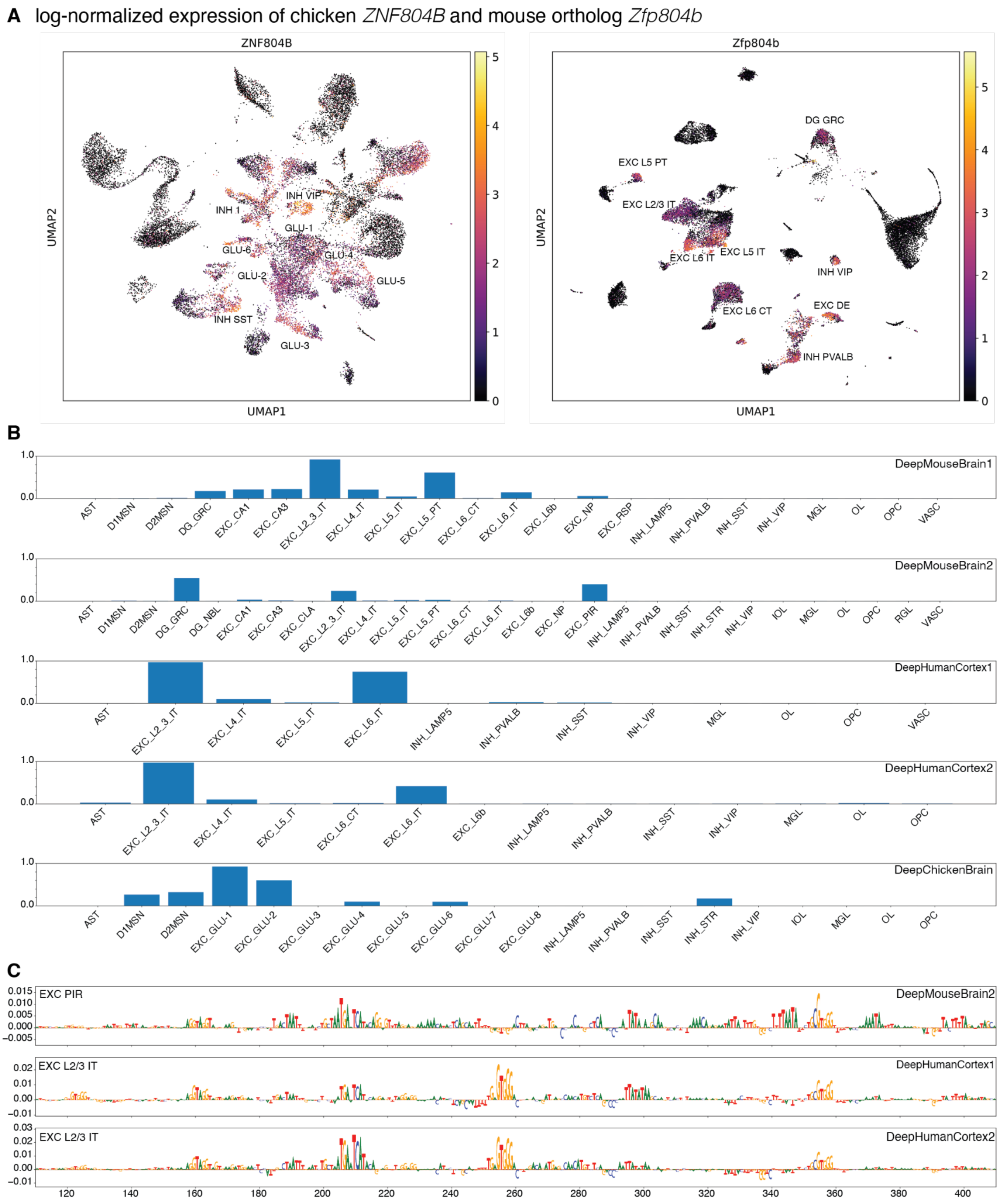


**Fig. S18. Predictions and contribution scores on the chicken *ZNF804B* enhancer region.**
**(A)** scRNA-seq UMAP showing *ZNF804B* expression in chicken telencephalon single cell data (left), and *Znf804B* expression in mouse telencephalon single cell data (right). (**B)** DeepMouseBrain1&2, DeepHumanCortex1&2 and DeepChickenBrain predictions on the chicken *ZNF804B* region (galGal6 chr2:21373470-21373970). (**C**) DeepMouseBrain2 and DeepHumanCortex1&2 contribution scores for their EXC PIR (DeepMouseBrain2) and EXC L2/3 IT (DeepHumanCortex1&2) predictions on the chicken *ZNF804B* region.

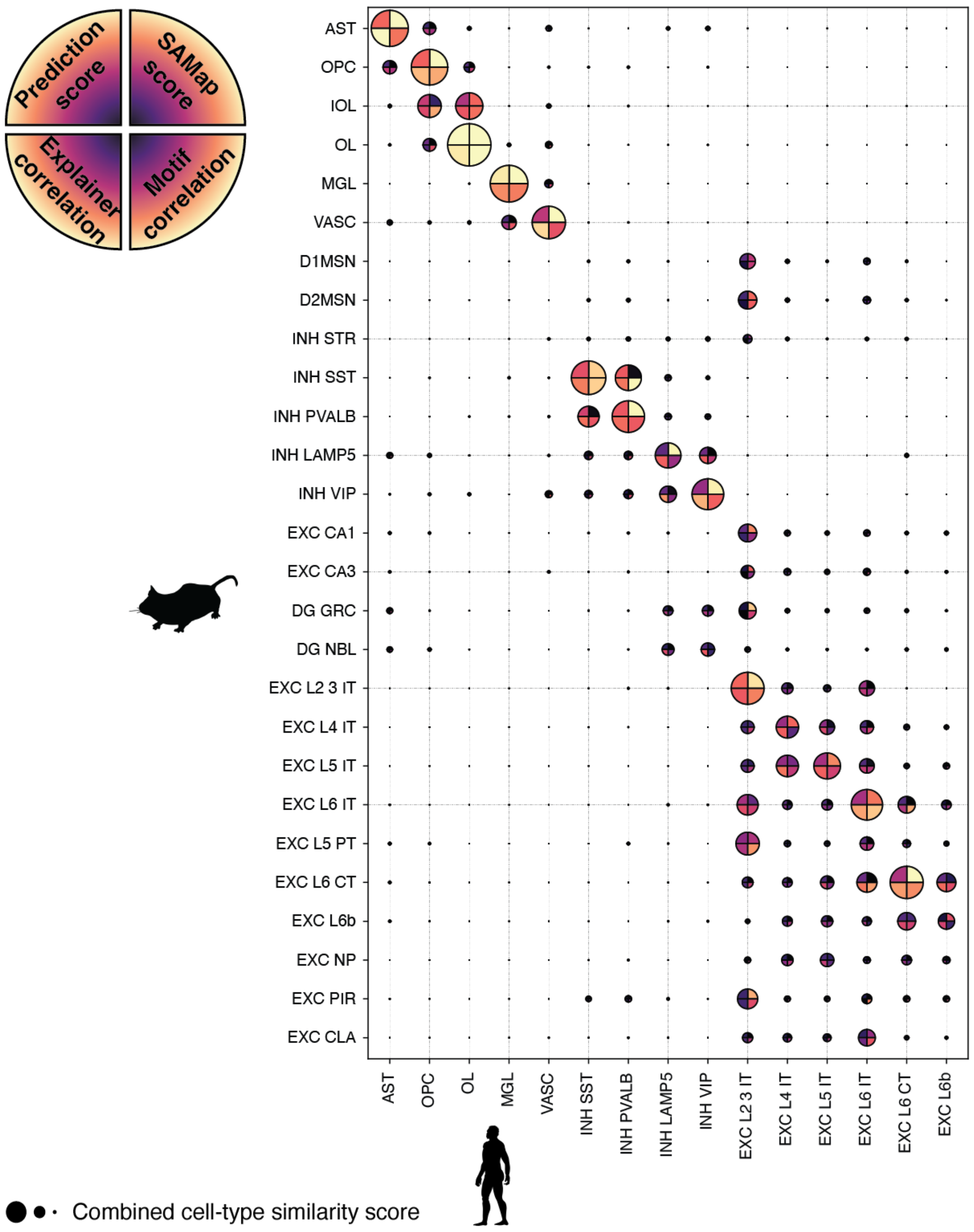


**Fig. S19. Summary of cell type comparison metric between human and mouse brain cell types.**
Combined cell-type similarity scores consisting of SAMap score, prediction scores, contribution score correlation and motif correlation between all available mouse and human cell types. The circle size depends on the mean score of all four metrics. SAMap scores are the average SAMap score from the DeepHumanCortex1 and DeepHumanCortex2 datasets against the complete mouse brain dataset (*30*). The predictions and contribution scores are the average median of scores on human regions (*24*) with DeepMouseBrain models and scores on mouse regions (*25*) with the DeepHumanCortex models. Motif correlation scores were merged for matching cell types in DeepMouseBrain1&2 and DeepHumanCortex1&2. All scores were standardized between 0 and 1. Negative correlation scores were set to zero.

### Supplementary tables

**Table S1: number of differentially accessible regions and cells per dataset.** Table containing the number of DARs after filtering steps, and number of regions used as input for TF-MODISCO per cell type.

**Table S2: DeepChickenBrain dataset and model information**
Table containing information on the number of identified DARs and the complete model architecture for the DeepChickenBrain dataset.

**Table S3: DeepMouseBrain1 dataset and model information**
Table containing information on the number of identified DARs and the complete model architecture for the DeepMouseBrain1 dataset.

**Table S4: DeepMouseBrain2 dataset and model information**
Table containing information on the number of identified DAR, and the complete model architecture for the DeepMouseBrain2 dataset.

**Table S5: DeepHumanCortex1 dataset and model information**
Table containing information on the number of identified DARs and the complete model architecture for the DeepHumanCortex1 dataset.

**Table S6: DeepHumanCortex2 dataset and model information**
Table containing information on the number of identified DARs and the complete model architecture for the DeepHumanCortex2 dataset.
